# Efficient and low-impact enzymatic glycosylation with robust sucrose synthase variants

**DOI:** 10.64898/2026.01.30.702896

**Authors:** Felipe Mejia-Otalvaro, David Delima, Mandy Hobusch, Brianna M. Lax, Catarina Mendonça, Gonzalo Bidart, Agata Matera, Alice Branger, Martina Escorial García, Carme Rovira, Ditte H. Welner

## Abstract

Glycosyltransferase-driven glycosylation enables environmentally mild synthesis of high-value chemicals, but industrial implementation is constrained by the cost of UDP-glucose. Sucrose synthase (SuSy) offers an attractive route for UDP-glucose recycling, yet inadequate operational stability has limited its use. Here, we report an integrated engineering workflow that overcomes the longstanding trade-off between efficiency and robustness in glycosyl donor recycling enzymes. We engineered *Gm*SuSy wild-type and obtained variants combining supra-wildtype activity (178%) with enhanced thermostability (ΔTm_app_ = 13.3 °C), solvent tolerance (70% retained activity in 25% DMSO) and a 123-fold longer half-life. The variants achieved total turnover numbers of ∼1 million (60 °C), supporting their industrial relevance. Mechanistic analyses revealed that long-range residue communication networks couple oligomeric interfaces with active sites, shifting conformational populations toward stable, catalytically competent states while increasing hydrophobic packing and reducing solvent accessibility. This enhanced robustness enabled significant process-level gains, including >90% conversion yields in indoxyl and MANT glycosylation. Techno-economic and life-cycle assessments indicate the potential to halve reaction costs and reduce environmental impacts threefold, establishing SuSy robustness as a key lever for sustainable industrial glycosylation.

## 1. Introduction

Glycosylated natural products constitute an industrially important class of compounds whose glycosyl moieties frequently confer beneficial physicochemical and biological properties compared to the aglycons^1^. These enhanced properties expand their application across the food, pharmaceutical, and agricultural sectors, as exemplified by glycosylated saponins used as vaccine adjuvants^2,3^, phenylethanoid glycosides employed for the treatment of neurodegenerative disorders^4^, steviol glycosides that serve as natural sweeteners^5^, and enzyme-based denim dye^6^.

Among the available routes to glycosylated natural products, glycosylation catalyzed by uridine diphosphate-dependent glycosyltransferases (UGTs) has been widely investigated, resulting in a diverse repertoire of UGTs with favorable operational properties, including thermostability^6–11^ and solvent tolerance^12,13^. However, the high cost of the sugar donor substrate, uridine diphosphate glucose (UDP-Glc) remains a challenge to industrial implementation. UDP-Glc regeneration with sucrose synthase (SuSy; EC 2.4.1.13) offers an attractive solution to this challenge. SuSy catalyzes the reversible interconversion of sucrose and UDP with UDP-Glc and fructose, and has been exploited in UGT-SuSy enzyme cascades^14,15^. However, industrial implementation requires a SuSy combining high catalytic UDP-Glc recycling efficiency with high operational stability.

Plant SuSys such as those from *Arabidopsis thaliana* (*At*SuSy) and *Glycine max* (*Gm*SuSy), efficiently regenerate UDP-Glc but lack operational stability^14–21^. In contrast, bacterial homologs such as *Ac*SuSy from *Acidithiobacillus caldus* are substantially more thermostable but display poor UDP specificity^14,22,23^. Low solvent tolerance further hampers glycosylation of commonly used hydrophobic substrates^24–27^. Strategies explored to address this, such as enzyme immobilization^28^ and solvent engineering ^26,29^, underscore the need for SuSys that are intrinsically robust in process conditions.

This limited operational robustness and efficiency of currently available SuSys restricts industrial glycosylation. For example, nothofagin production requires prolonged operation times at elevated temperatures, under which SuSy is unstable^19,26,30^. Similarly, methyl anthranilate (MANT) glycosylation has relied on stoichiometric UDP-Glc because of insufficient SuSy stability^31^, while indican biosynthesis remains limited to ∼65% conversion despite the use of large SuSy excesses^6^. Consequently, the regeneration enzyme is a major contributor to process cost and environmental impact.

Bridging the longstanding divide between catalytic efficiency and operational stability has therefore become a central challenge in developing industrial sugar donor-recycling enzymes. Previous studies have addressed this through enzyme discovery and protein engineering of both bacterial and plant SuSys^22,24,32–34^. For instance, *N*-terminal truncation of the plant *Gu*SuSy to resemble bacterial SuSys doubled its half-life at 50 °C to reach 100 min^25^. Similarly, consensus engineering enhanced the half-lives of both bacterial *Nm*SuSy and plant *Sr*SuSy at 55 °C, from 1.2 to 51 min and from 1.39 to 3.55 h, respectively^35,36^. Despite these advances, however, the thermostability of these engineered variants remains substantially below that of *Ac*SuSy, which displays a half-life of 4 h at 60 °C^22^.

A major obstacle to further improvement is our limited understanding of the molecular determinants governing SuSy stability and activity. Available structures reveal a tetrameric enzyme in which each monomer, comprising approximately 800 residues, is organized into four domains^37,38^. In plant SuSys, these include a cellular targeting domain (CTD), an ENOD40 peptide-binding domain (EPBD), and two GT-B fold domains (Figure S1). These contain the active site and form a hinge-latch system that controls active site access and catalytic activity^37,38^. These structural features have provided insight into possible catalytic mechanisms and cellular interactions^37–41^. However, beyond the general notion that oligomerization has the potential to enhance protein stability^33,42,43^, the structural determinants of SuSy stability remain largely unexplored.

In this work, we developed SuSys that unite high operational robustness with efficient UDP-sugar regeneration, addressing the key limitations of UGT-SuSy cascades. To this end, we integrated consensus sequence mutagenesis, ancestral sequence reconstruction (ASR), and deep learning-based protein sequence design. The resulting SuSy variants retained wild-type activity while reaching melting temperatures up to 76 °C, half-lives of 7.4 h at 60 °C, twofold higher soluble expression, and total turnover numbers (TTN) up to one million at 60 °C, making SuSy suitable even for bulk chemical production. Molecular dynamics analyses indicated strengthened hydrophobic packing and reduced solvent-accessible surface area (SASA) to be the main drivers of robustness in our variants. Residue-correlation network analysis further linked changes at the oligomeric interface with the active site, suggesting an allosteric mechanism that modulates both activity and thermostability. We further demonstrate how SuSy robustness translates directly to process productivity, doubling the conversion yields for MANT and indoxyl glycosylation and reaching up to 96% conversion. Preliminary techno-economic and life-cycle assessments indicate that these improvements can reduce reaction costs and environmental impacts by up to 6.4-fold compared with literature-reported systems. These findings establish robust SuSy as a central biocatalyst for low-impact glycosylation of natural products and enzyme robustness as a key lever for sustainable industrial enzymology.

## 2. Results and discussion

### 2.1. Robust SuSy variants were generated through multi-strategy enzyme engineering

To alleviate the bottleneck of UDP-Glc recycling in enzymatic cascades, we set out to engineer the stability of SuSy. *Gm*SuSy wild-type (hereafter referred to as WT) was the selected template based on its extensive characterization and superior cascade performance^19,26,30,44^. We implemented an integrated semi-rational design strategy combining rational engineering, consensus-guided engineering, ancestral sequence reconstruction, and deep learning-based protein sequence design (Figure 1a). The resulting variants were screened for thermostability and activity, using WT as the baseline reference and *Ac*SuSy_M_ (L637M-T640V), which displays improved UDP affinity compared to the bacterial wild-type^45^, as a thermostable benchmark. A detailed description of the engineering strategy can be found in Methods, and a brief overview is given below.

**Figure 1.**
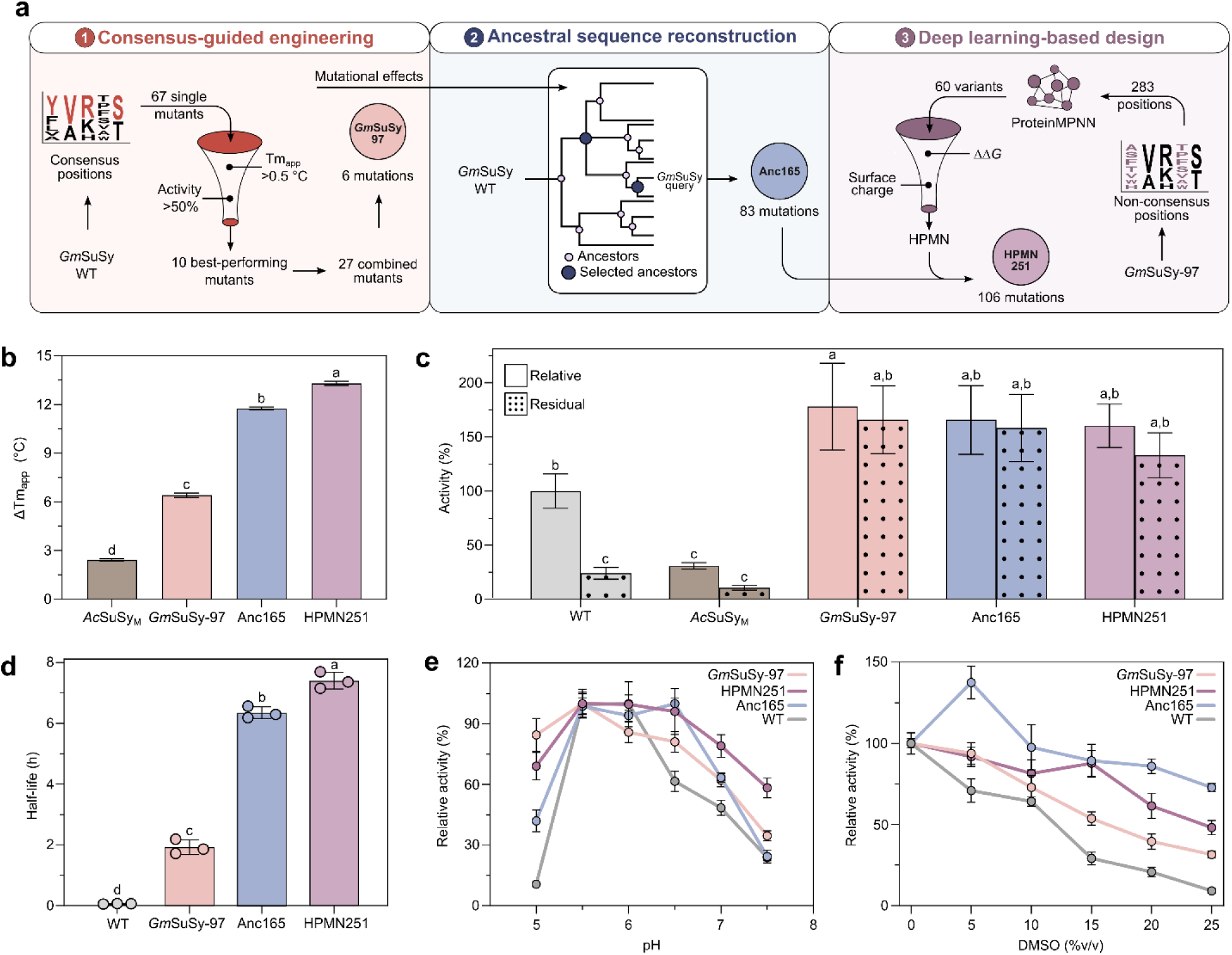
Integrated stability engineering improves SuSy robustness across thermal and chemical stress conditions. **a**, Overview of the integrated engineering workflow for robust SuSy design. Screening and combination of consensus-guided mutations resulted in *Gm*SuSy-97. Ancestral sequences were modified according to detrimental and beneficial mutations found in the consensus approach, yielding Anc165 as the top performer. Non-consensus positions were modified through inverse folding, yielding HPMN. Further incorporation of HPMN mutations into Anc165, resulted in HPMN251. **b**, Thermodynamic stability was assessed by differences in Tm_app_ (ΔTm_app_) relative to WT. **c,** Activities were quantified with reference to WT. Dotted bars refer to residual activity after 5 min incubation at 60 °C. **d**, Half-lives calculated at 60 °C showed improved operational stability. **e**, pH-dependent activity profiles for SuSy variants. Relative activity was calculated separately for each variant by assigning its highest activity across the tested pH range a value of 100%. **f**, Enhanced DMSO tolerance was assessed by comparing the relative activities of solvent-treated samples to those of untreated samples. All measurements were performed by triplicate (*n*=3), bar height reflect means and error bars refer to standard deviations including error propagation for relative values such as ΔTm_app_, relative and residual activities. Bars with different superscripts differ statistically significantly between them according to analysis of variance (ANOVA) with Tukey’s post hoc test (*p* < 0.05).

First, 67 semi-rationally designed single mutants generated from a consensus sequence approach were screened and ranked according to apparent melting temperature (Tm_app_). The maximum increase was 2.71 °C, and the top 10 variants retaining >50% activity were combined into 27 higher-order mutants (Tables S1-S3). The sextuple variant *Gm*SuSy-97 was the top performer, with a 6.4 °C increase in Tm_app_ and 178% activity (Figure 1b,c; Figure S2).

We then turned to ASR, as ancestral enzymes are often more robust than their extant counterparts^46^. Three phylogenetic trees were constructed with varying positional restrictions, from which six ancestral sequences were selected. These were further modified by introducing and reverting beneficial and detrimental mutations previously identified in consensus designs. Four ancestral variants were produced in soluble form and characterized, leading to Anc165 as the top performer with 83 single-point mutations (including the six mutations in *Gm*SuSy-97), 11.8 °C increase in Tm_app_, and 166% activity (Figure 1b,c; Tables S4 and S5).

To further diversify the SuSy sequence, we turned to deep learning-based protein sequence design. 60 variants were generated by modifying non-conserved positions using an inverse folding approach (ProteinMPNN^47^). These sequences were characterized *in silico*, evaluating their predicted structural and functional integrity, and four sequences were selected for experimental characterization. Only one of these, HPMN, could be expressed and characterized. HPMN, carrying 47 single point mutations (Table S4), showed around 5-fold higher soluble expression than the previous variants (Table S6), but had reduced Tm_app_ (−3.4 °C) and a loss of 79% in activity compared to Anc165 (Tables S5 and S7). We opted to combine the sequences of HPMN and Anc165, hypothesizing that this might give rise to a SuSy variant with excellent performance in soluble expression, stability, and activity. Indeed, the resulting HPMN251 variant, carrying 106 single point mutations (Table S4), displayed a 13.3 °C increase in Tm_app_, 160% activity, and twofold expression level (Figure 1b,c; Figure S3; Tables S6 and S7).

Overall, the screening of 108 SuSy mutants resulted in three outstanding variants: *Gm*SuSy-97, Anc165, and HPMN251. They exhibited progressively increased Tm_app_ ranging from 68.9 °C to 75.9 °C without loss of activity relative to the WT (Figure 1b,c). This was achieved by incorporation of mutations distal from the active site (> 5 Å) and distributed all over the SuSy domains, starting from 6 mutations in *Gm*SuSy-97, adding additional 77 in Anc165 to a total of 83, and finally adding 23 more to a total of 106 mutations in HPMN251 (Figure S4; Table S8). These three variants outperformed the benchmark, *Ac*SuSy_M_, in thermostability, activity and soluble expression (Figure 1b,c; and Table S6).

### 2.2. Robust SuSy variants are operationally and chemically stable

To further assess the robustness of our variants, we measured residual activity after 5 min incubation at 60 °C, a temperature close to the Tm_app_ of WT and therefore suitable for identifying variants with enhanced kinetic stability. *Gm*SuSy-97, Anc165, and HPMN251 retained statistically alike activity after incubation, while WT retained only 24% (Figure 1c). Consistent with this observation, their half-lives at 60 °C increased by 32-, 107-, and 123-fold, respectively, changing from approximately 3.7 min (WT) to 7.4 h (HPMN251) (Figure 1d; Figure S5). HPMN251 reached a half-life nearly twofold higher than the one of the highest previously reported value for a SuSy at 60 °C, *Ac*SuSy (4 h)^22^, despite being assayed in the absence of sucrose. At 55 °C, a temperature that enables direct comparison with other literature-reported SuSys, Anc165 and HPMN251 retained half-lives greater than 20 h, exceeding WT (2.5 h), and engineered *Nm*SuSy (51 min)^36^ and *Sr*SuSy (3.6 h)^37^ (Figure S6).

Because process performance depends not only on operational stability but also on catalytic activity, we next evaluated the kinetic and turnover properties of our thermostable SuSy variants (Table 1 and Figure S7). We found increases in k_cat_ of up to threefold accompanied by increases in K_M_ for sucrose of up to fourfold, resulting in a twofold reduction in catalytic efficiency (k_cat_/K_M_). However, the substantially prolonged half-lives outweighed this trade-off, enabling TTNs of up to 1 million, three orders of magnitude higher than WT (Table 1), thus positioning these variants as high-performing biocatalysts.

**Table 1.** Kinetic parameters for *Gm*SuSy variants determined in the sucrose cleavage reaction.

| Parameter* | WT | *GmSuSy*-97 | Anc165 | HPMN251 |
| --- | --- | --- | --- | --- |
| $k_{\text{cat}}$ ($\text{s}^{-1}$) | $10.7 \pm 0.2^{\text{a}}$ | $24.5 \pm 1.3^{\text{b}}$ | $31.0 \pm 1.0^{\text{c}}$ | $10.6 \pm 0.5^{\text{a}}$ |
| $K_{\text{M}}$ (mM) | $79.4 \pm 4.0^{\text{a}}$ | $294.1 \pm 16.1^{\text{b}}$ | $324.3 \pm 38.9^{\text{b}}$ | $173.9 \pm 7.5^{\text{c}}$ |
| $k_{\text{cat}}/K_{\text{M}}$ ($\text{s}^{-1} \text{M}^{-1}$) | $134.7 \pm 7.4^{\text{a}}$ | $83.4 \pm 6.3^{\text{b}}$ | $95.7 \pm 11.9^{\text{b}}$ | $61.1 \pm 3.7^{\text{c}}$ |
| $k_{\text{d}}$ ($\text{s}^{-1}$) | $3.1 \times 10^{-3} \pm 4.9 \times 10^{-4}^{\text{a}}$ | $1.0 \times 10^{-4} \pm 1.2 \times 10^{-5}^{\text{b}}$ | $3.0 \times 10^{-5} \pm 9.1 \times 10^{-7}^{\text{b}}$ | $2.6 \times 10^{-5} \pm 9.5 \times 10^{-7}^{\text{b}}$ |
| TTN ($k_{\text{cat}}/k_{\text{d}}$) | $3.4 \times 10^3 \pm 5.4 \times 10^2^{\text{a}}$ | $2.4 \times 10^5 \pm 3.2 \times 10^4^{\text{b}}$ | $1.0 \times 10^6 \pm 4.4 \times 10^4^{\text{c}}$ | $4.1 \times 10^5 \pm 2.3 \times 10^4^{\text{d}}$ |

\* All measurements were performed in triplicate ( $n=3$ ) and standard deviations include error propagation for relative values such as $k_{\text{cat}}/K_{\text{M}}$ and TTN. Means with different superscripts differ statistically significantly, according to analysis of variance (ANOVA) with Tukey's post hoc test ( $p < 0.05$ ). $K_{\text{M}}$ is for sucrose. $k_{\text{d}}$ refers to thermal deactivation constant at 60 °C. TTN indicates total turnover number. All reactions were run under optimized conditions for each variant.

Thermostabilization was also coupled with a higher pH tolerance (Figure 1e). Compared with WT, the three variants retained higher activity under both acidic and alkaline conditions, with the largest gains observed above pH 6.0, a regime that is particularly relevant for UGT-coupled glycosylation cascades^48^ (Figure 1e). Among the variants, HPMN251 displayed the broadest pH tolerance, maintaining >60% relative activity across the pH range tested.

Finally, a notable feature of our engineered variants is enhanced tolerance to DMSO, a key cosolvent in glycosylation cascades involving poorly soluble substrates. *Gm*SuSy-97, Anc165, and HPMN251 all showed improved DMSO tolerance relative to WT at 60 °C. Anc165 retained up to 70% activity in the presence of 25% (v/v) DMSO, compared to only 9% for WT, and it even displayed a higher activity (137.5%) in the presence of 5% DMSO (Figure 1f). Increasing DMSO concentrations reduced ΔTm_app_, but the decline was substantially slower for Anc165 and HPMN251 than for WT (Figure S8), indicating increased resistance to solvent-induced unfolding. By contrast, tolerance to methanol and acetonitrile was not improved, suggesting a different mode of action for these solvents (Figure S8).

### 2.3. The molecular basis of enhanced SuSy robustness

To elucidate how the identified mutations collectively enhance robustness and catalysis, we employed structural analyses and molecular dynamics (MD) simulations of *Gm*SuSy-97 and Anc165, which showed the largest increases in Tm_app_ respective to the previous best variant (Figure 1b). Simulations were performed at 60 °C, corresponding to the temperature optimum of WT and providing conditions suitable for resolving structural and dynamic differences relevant at stress conditions.

Tetrameric models revealed no major structural rearrangements, with C_α_ RMSDs of 0.002 Å and 1.132 Å relative to WT for *Gm*SuSy-97 and Anc165, respectively (Figure S9). Electrostatic surface analysis suggested that Anc165 displays more extensive positively charged surface patches than WT at its optimal pH, including at the tetramer interfaces (Figure S10). This indicates that electrostatic remodeling may contribute to its enhanced robustness^49,50^, consistent with previous reports linking positive surface charge to solvent tolerance in enzymes^49,50^.

The variants differed primarily from WT in their dynamic structural organization. During 3.6 μs of MD simulations, *Gm*SuSy-97 and Anc165 formed 366 and 774 additional hydrophobic contacts, respectively (Figure 2a), accompanied by reductions in SASA of 210 and 820 Å² (Figure 2b). The number of hydrogen bonds remained essentially unchanged, whereas salt bridges decreased slightly (Figure S11). Together, these observations indicate that the engineered variants adopt more compact conformational ensembles through enhanced hydrophobic packing.

**Figure 2.**
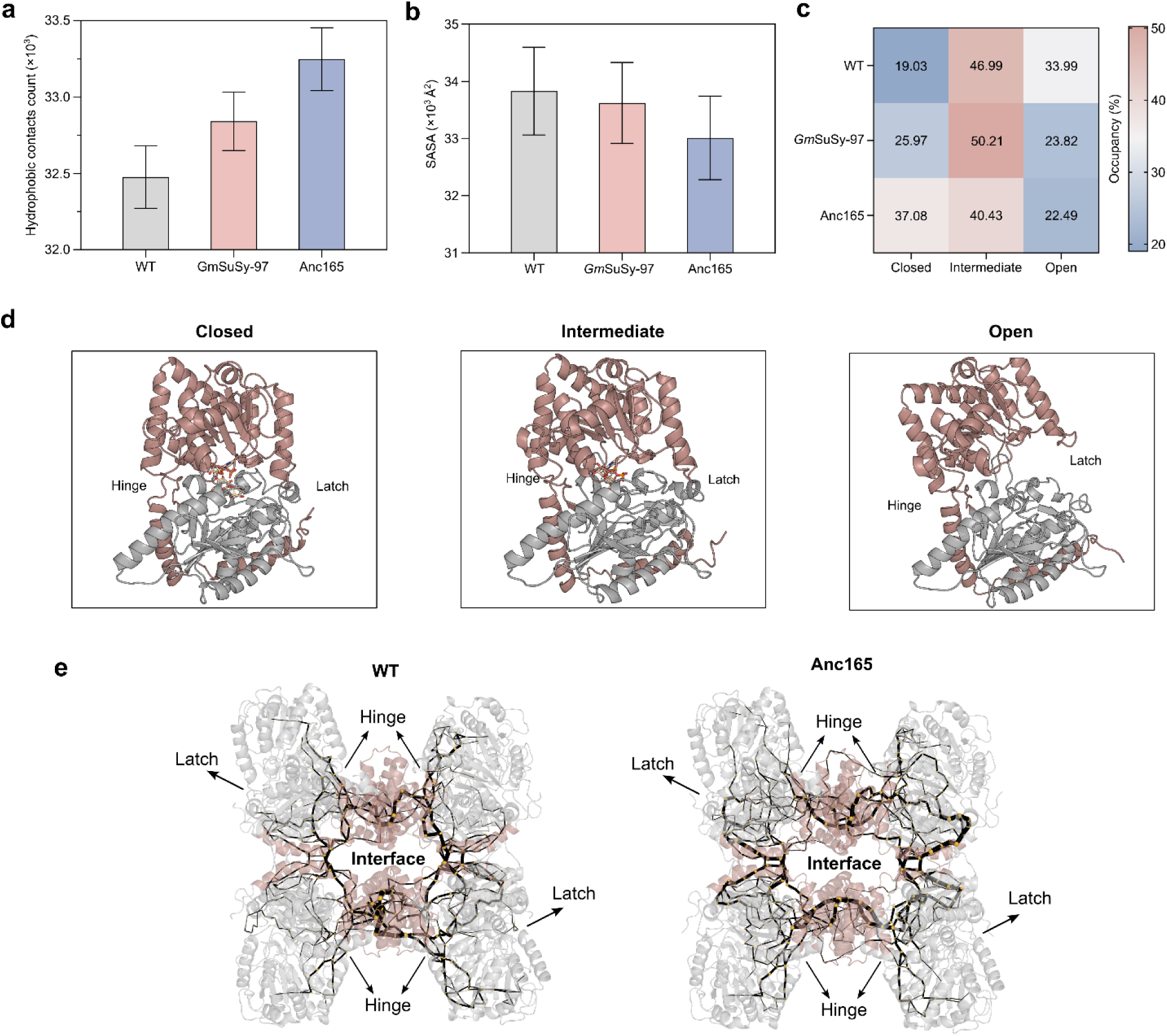
Enhanced SuSy robustness correlates with increased hydrophobic contacts, reduced SASA, and long-range residue communication that shifts conformational populations toward closed states. Differences between WT, *Gm*SuSy-97, and Anc165 after analyzing the trajectories from a 3.6 µs MD simulations of SuSy as a tetramer. **a,** Count of hydrophobic interactions. **b**, SASA of whole protein. **c**, Occupancy (%) of closed, intermediate, and open conformational states determined by conformational clustering analysis. **d**, Representative structures of Anc165 from each cluster. The C-terminal and N-terminal domains of the GT-B fold are colored rosy-beige and gray, respectively. Other SuSy domains are omitted to better illustrate the changes in the GT-B fold and hinge-latch system. **e**, SPM representing protein dynamics as a network in which residues are nodes connected by edges. Residues that contribute strongly to global protein dynamics are depicted as larger spheres, whereas thicker edges denote highly correlated conformational changes between pair of nodes. The tetramer interface is colored rosy-beige. Bar heights and error bars refer to means and standard deviations of the count along all the simulation (2 ns per frame; *n* = 1800), respectively.

Clustering analysis of the MD trajectories resolved three dominant conformational states in accordance with the opening and closing of the GT-B fold^37,38^ (Figure 2c-d). Of these, only one cluster represented a catalytically competent conformation (closed), exhibiting productive catalytic bonds and binding free energies for both sucrose and UDP that correlated with the experimentally determined K_M_ values for sucrose (Figures S12 and S13). By contrast, substrate binding was prohibitively unfavorable (> −1 kcal mol⁻¹) in the intermediate and open states clusters. The occupancy of the active conformation was substantially increased in both engineered variants relative to the WT (Figure 2c-d), possibly mediated by the 3 and 11 mutations in the hinge-latch system of the GT-B fold in *Gm*SuSy-97 and Anc165, respectively (Table S8 and Figure S14). This suggests that redistribution of the conformational ensemble toward the catalytically competent state is the mechanistic basis for the enhanced activity of the variants.

Although these analyses did not identify a dominant stabilizing region, shortest-path map (SPM) analysis^51,52^ suggested an association between SuSy oligomerization and stability (Figure 2e). The strongest communication pathways converged at the tetramer interface and involved more residues in Anc165 than in WT (393 versus 254, Table S9). These pathways connect the tetramer interface to the active site (Figure 2e), linking oligomerization and catalysis. This link is further supported by our attempts to engineer the tetrameric interface, where we found increased Tm_app_ by 1.6-3.5 °C but reduced catalytic activity by 34-66% relative to WT (Figure S15; Tables S10 and S11). Incidentally, 3 and 17 residues implicated in the WT paths are mutated in *Gm*SuSy-97 and Anc165, respectively (Table S9), providing a mechanistic explanation for their increased k_cat_ despite all mutations being >5 Å from the active site.

### 2.4. Robust SuSy variants reduce the cost and environmental footprint of UGT-SuSy cascades

To assess the impact of enhanced SuSy robustness on process performance and sustainability, we tested our variants in two glycosylation cascades known to be limited by SuSy performance^6,31^.

First, we evaluated the glycosylation of MANT (Figure 3a), a food additive (grape flavor), and bird deterrent^53,54^. Previous attempts to implement WT in a MANT glycosylation cascade coupled with UGT72B68 were abandoned due to WT inhibition by MANT and/or the cosolvent DMSO^31^. We hypothesized that our variants would perform better, since they are more tolerant to DMSO (Figure 1f). Initial tests showed significantly increased MANT-*N*-glucose yields (Figure S16), with only small differences between *Gm*SuSy-97, Anc165, and HPMN251, whereas WT comparatively underperformed over time.

**Figure 3.**
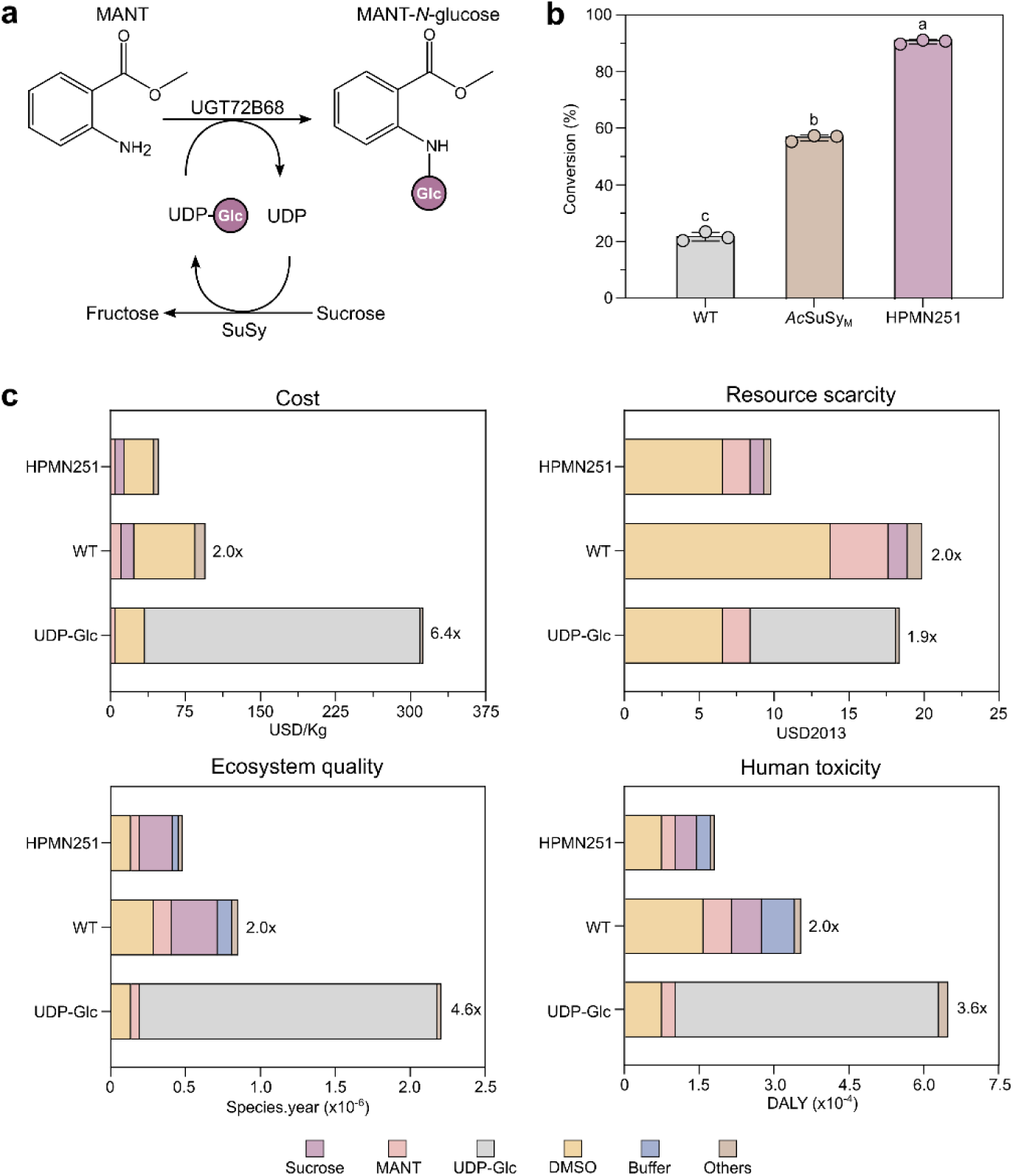
Robust SuSy enhanced MANT glycosylation efficiency. **a**, Schematic depiction of the reaction cascade for MANT glycosylation using UGT72B68 and SuSy for *in situ* UDP-glucose regeneration. Glc stands for glucose. **b**, MANT-*N*-glucose formation under optimized conditions for HPMN251, WT, and *Ac*SuSy_M_. Measurements were performed in triplicate (*n*=3), bar height reflects means and error bars refer to standard deviations. Bars with different superscripts differ statistically significantly according to analysis of variance (ANOVA) with Tukey’s post hoc test (*p* < 0.05). **c**, Preliminary life cycle assessment and cost analysis of MANT-*N*-glucose production under three different systems: UDP-Glc/Literature^31^, WT, and HPMN251. Impact increases compared to HPMN251 are given next to the bars. The “Others” contributor category refers to reaction components with a minimum contribution (<5%), including water, UDP, enzymes, and waste treatment. Buffer contribution was also considered in Others in the UDP-Glc system.

Since HPMN251 showed higher soluble expression, which impacts the process cost positively, this variant was chosen for further optimization of the MANT glycosylation cascade. Optimization of sucrose concentration, pH, and temperature, coupled with increased glycosyltransferase loading (Figure S17), resulted in 90% conversion, substantially outperforming *Ac*SuSy_M_ and WT while matching the performance of the original process without requiring stoichiometric UDP-Glc (Figure 3b).

In addition to improved conversion, HPMN251 approximately tripled titer, rate, and specific yield, compared to WT (Table 2). Remarkably, HPMN251 achieved titers and production rates comparable to those obtained with stoichiometric UDP-Glc while increasing the specific yield by 30%, despite the addition of an extra enzyme. This improvement was enabled by a lower optimal UGT loading (Figure S17). One possible explanation is that the lower UDP concentration in the reaction reduced product inhibition of the UGT^48^.

**Table 2.** Techno-economical metrics for the production of MANT-*N*-glucose and indican across different biocatalytic scenarios.

| Product | Scenario | Titer (g L<sup>-1</sup>) | Rate (g L<sup>-1</sup> h<sup>-1</sup>) | Specific yield (g<sub>p</sub> g<sub>e</sub><sup>-1</sup>) |
| --- | --- | --- | --- | --- |
| MANT-*N*-glucose | HPMN251 | 5.6 | 0.3 | 60.7 |
|  | WT | 1.9 | 0.1 | 17.5 |
|  | UDP-Glc | 5.6 | 0.3 | 46.7 |
| Indican | HPMN251 | 28.5 | 4.7 | 20.1 |
|  | WT | 10.3 | 1.7 | 7.3 |
|  | Literature | 8.4 | 1.4 | 2.7 |

g<sub>p</sub> and g<sub>e</sub> refer to grams of product and grams of enzyme, respectively.

To determine whether this improved process performance translates into economic and environmental benefits, we assessed the implications of three scenarios for the enzymatic synthesis of 1 kg MANT-*N*-glucose: stoichiometric use of UDP-Glc as described previously^31^, and *in situ* recycling of UDP-Glc with WT or HPMN251 (Figure S18). The corresponding inventories, cost analysis, and life-cycle impact assessment can be found in Tables S12-S14.

We found the HPMN251 scenario to have the potential to reduce raw material cost by 6.4-fold compared to stoichiometric UDP-Glc, and to halve the cost compared to WT (Figure 3c). Additionally, this robust variant has the potential to reduce resource scarcity, human toxicity, and ecosystem quality impacts by two to fivefold (Figure 3c). Implementing SuSy mitigated the high costs and environmental burden associated with UDP-Glc, shifting the dominant contribution to sucrose and DMSO (Figure 3c). Suggestions to mitigate these contributions can be found in Figure S19, and proof of concept for using SuSy-expressing biomass, which is closer to the assumption applied in the LCA framework, is also demonstrated (Figure S20). Overall, HPMN251 has the potential to reduce the environmental impact of enzymatic MANT glycosylation by 50% compared to UDP-Glc and WT systems (Table S14).

To further evaluate the impact of SuSy robustness on process performance, we investigated the glycosylation of indoxyl to produce indican, a challenging reaction involving a three-enzyme cascade and a substrate concentration of 100 mM (Figure 4a). This system has previously been reported to reach 65% conversion after 32 h with WT, in which SuSy was identified as the bottleneck^6^.

**Figure 4.**
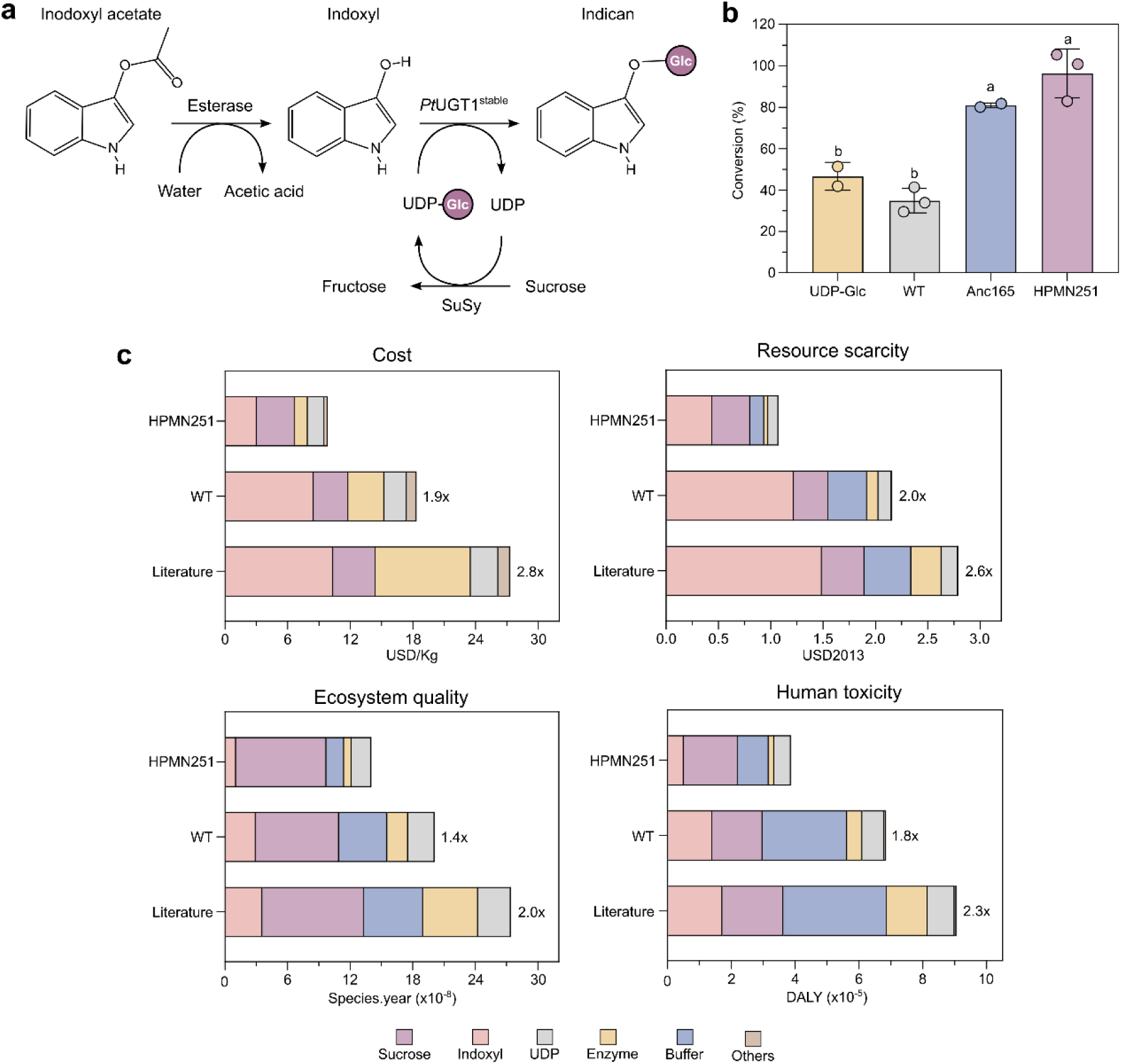
Robust SuSy enhances indican production efficiency. **a**, Schematic representation of the three-enzyme reaction cascade for indican biosynthesis from indoxyl acetate implementing SuSy for UDP-Glc regeneration. **b**, Indican formation under optimized conditions for HPMN251, Anc165, WT, and UDP-Glc. Conversion was measured in triplicate (*n*=3) for WT and HPMN251 and in duplicates (*n*=2) for Anc165 and UDP-Glc. Bar height reflect means and error bars refer to standard deviations. Bars with different superscripts differ statistically significantly according to analysis of variance (ANOVA) with Tukey’s post hoc test (*p* < 0.05). **c**, Preliminary life cycle assessment and cost analysis of 1 kg indican biosynthesis using three different systems: HPMN251, WT, and a previously reported system based on WT and denoted as *Literature*^6^. Impact increases compared to HPMN251 are given next to the bars. The “Others” contributor category refers to reaction components with a minimum contribution (<5%), including water and waste treatment.

After cascade optimization, including increasing sucrose concentration from 200 to 600 mM and temperature from 30 to 54 °C (Table S15), 96% conversion within 6 h was achieved with HPMN251, surpassing the yield with stoichiometric UDP-Glc and WT by approximately 2-fold (Figure 4b). Implementation of HPMN251 improved titer and rate by threefold and the specific yield by approximately an order of magnitude, corresponding to a reduction of enzyme load from 364 g to 50 g per kg of indican (Table 2).

Importantly, this translates directly into halved reaction cost and environmental footprint across endpoint categories (Figure 4c; Tables S16-S18). Notably, resource scarcity and cost, which are dominated by substrate and enzyme in the WT system, are now evenly balanced by multiple factors (Figure 4c). In the optimized HPMN251 system, sucrose contributes significantly to all impact categories, accounting for 62% of ecosystem quality, 44% of human toxicity, and 34% of resource scarcity impacts, making it the key target for process improvement. Sensitivity analysis shows that with one sucrose recycling cycle these effects can be reduced by 15-25% (Figure 19).

## 3. Conclusions

Efficient UDP-Glc regeneration remains a central requirement for the industrial implementation of UGT-mediated glycosylation. Although SuSy has long been recognized as an attractive biocatalyst for this purpose, its use has been constrained by limited operational robustness under the reaction conditions needed for industrial implementation. Here, we show that this limitation can be overcome by engineering SuSy variants that combine high UDP-Glc recycling activity with substantially improved thermodynamic stability, kinetic stability, solvent tolerance and soluble expression. The resulting variants outperformed both the *Gm*SuSy template and a thermostable bacterial benchmark, and enabled TTNs up to 1 million. These properties place SuSy in a performance range compatible with industrial biocatalysis, including applications in which enzyme cost and lifetime are decisive process parameters.

A notable outcome of this work is that robustness was improved without the activity losses commonly associated with enzyme stabilization. The mutations in the best-performing variants were distributed across the protein, distant from the active site (> 5 Å), and did not produce major global structural rearrangements. Instead, the enhanced robustness was associated with progressive structural compaction, reflected by increased hydrophobic contacts and reduced SASA. These features provide a plausible molecular basis for both improved thermostability and DMSO tolerance, as tighter hydrophobic packing and reduced solvent penetration have previously been linked to solvent-resistant enzymes^55^. In Anc165, an increased positive surface potential may further contribute to solvent resistance, suggesting that both internal packing and surface electrostatics can be exploited to improve SuSy robustness^49,50^.

The molecular analyses further indicate that SuSy stability and activity are coupled through long-range dynamic communication. Residue-correlation networks revealed communication pathways connecting the tetrameric interface with the GT-B fold hinge-latch system and the active site. These pathways were strengthened in the engineered variants and coincided with a shift in the conformational ensemble toward catalytically competent closed states. This provides a mechanistic explanation for the increased activity observed despite the absence of active-site mutations. It also helps explain why direct interface engineering produced gains in thermostability but accompanied by substantial activity losses, whereas the integrated engineering strategy used here improved stability and activity simultaneously. Thus, the tetrameric interface appears not only to contribute to SuSy structural integrity, but also to participate in the dynamic regulation of catalysis. Our results not only provide a mechanistic explanation for the activity changes observed in *Gm*SuSy-97 and Anc165, but also offer broader insight into widely observed effects in other SuSy homologs, where changes in the EPBD and CTD domains may influence activity and oligomeric state^39–41^.

The practical impact of SuSy robustness was demonstrated in two challenging glycosylation cascades. In MANT glycosylation, HPMN251 enabled conversion comparable to stoichiometric UDP-Glc supply while maintaining the economic advantage of *in situ* co-substrate recycling. Remarkably, this was achieved without compromising process performance: titers and rates matched those obtained with UDP-Glc, and the specific yield increased despite the introduction of an additional enzyme. This illustrates that enzymatic co-substrate recycling can provide benefits beyond replacing an expensive sugar nucleotide, including improved cascade efficiency and reduced enzyme demand. In indican biosynthesis, HPMN251 removed SuSy as the process bottleneck, enabling high conversion within substantially shorter reaction times and reducing the enzyme loading required per kilogram of product. Across both cascades, these improvements translated directly into lower projected costs and reduced environmental impacts.

The techno-economic and life-cycle analyses highlight enzyme robustness as a major lever for sustainable biomanufacturing. By increasing enzyme lifetime and enabling higher conversions under process-relevant conditions, the engineered SuSy variants reduced the relative contribution of enzyme and UDP-Glc to cost and environmental burden. At the same time, the analyses revealed new improvement targets that become visible only once SuSy is no longer limiting. In particular, sucrose and DMSO emerged as important contributors in the optimized processes. Therefore, future development should focus on efficient reuse or valorization of sucrose and fructose, improved co-solvent strategies, and solvent recycling. Further increases in soluble SuSy expression will also be important to fully exploit the cost and sustainability potential of the engineered variants.

Overall, this work establishes robust SuSy as an enabling platform for *in situ* UDP-Glc regeneration in UGT-mediated glycosylation. By transforming SuSy from a cascade bottleneck into a stable and efficient regeneration module, the engineered variants expand the operational window of enzymatic glycosylation and improve its economic and environmental viability. More broadly, these findings demonstrate that robustness-focused engineering of co-substrate recycling enzymes can unlock process-level gains that extend beyond the engineered enzyme itself, providing a general strategy for advancing sustainable industrial biocatalysis.

## 4. Methods

### 4.1. Materials

All chemicals and reagents used in this study were of analytical grade and were purchased from commercial suppliers unless otherwise stated. Primers were synthesized by Integrated DNA Technologies (IDT) and are listed in Table S19. Synthetic genes were obtained from Biomatik or Twist Bioscience, and the amino acid sequences they encode are listed in Table S4.

### 4.2. Consensus and rational sequence designs

The *Gm*SuSy wild-type sequence (UniProt accession number P13708) was selected as the template. A three-dimensional structure model of monomeric WT was submitted to the FireProt web server^56^, which generated a set of 66 candidate stabilizing single mutations based on a consensus sequence analysis and free-energy calculation (ΔΔG). All mutations were manually inspected in PyMOL v.2.5.4, and 12 mutations were excluded as they could potentially disrupt molecular interactions, exhibited a high risk of steric clashes, or were located within the catalytic active site and tetramer interface (Tables S20 and S21). Additional single mutations with the potential to strengthen molecular interactions were included, based on visual inspection of the structural model (Table S21). After activity and thermostability characterization (described in subsection 4.6), ten mutations (S152Y, H534Y, A395M, L453F, S523P, V671F, N775K, C660A, N694H, and Q754K) were selected based on changes in Tm_app_ of the second (>0.5 °C) and first thermal transition (>1.5 °C), and residual activity (Figure S2 and Tables S1-S3). After combinatorial screening of 27 progressive combinations, the *Gm*SuSy-97 variant was obtained involving six mutations: S152Y, A395M, L453F, V671F, N775K, S523P.

### 4.3. Ancestral sequence reconstruction

The *Gm*SuSy wild-type sequence (UniProt accession number P13708) was input to the FireProtASR web server^57^ with standard settings, including filters of minimal and maximal sequence identity of 30 and 90%, respectively. Three independent phylogenetic trees were generated using 1) no positional restriction, 2) restriction of residues comprising the UDP-binding site (Table S20), and 3) restriction of the conserved QN motif residues Q646, R649, R651, and N652^45^. Two ancestors were selected per phylogenetic tree: one corresponding to the closest ancestor leading toward the query sequence and a second representing the following closest ancestor that represented a significantly divergent clade (Figure S21). Then, beneficial and detrimental mutations previously identified in consensus and rational designs were introduced and reverted, respectively. The integrity of the resulting sequences was verified *in silico*, based on predicted solubility^58^, sequence identity, surface charge^59^, and structural similarity (RMSD) relative to WT (Table S22). Only Anc180, Anc165, Anc111, Anc101 could be expressed in soluble form and therefore were subsequently characterized (Table S5). Anc294 and Anc290 could not be expressed.

### 4.4. ProteinMPNN variants

ProteinMPNN-based variants were designed using a modified pipeline adapted from Sumida *et al* (2024), which integrates hotspot identification based on Jensen-Shannon divergence with sequence design using ProteinMPNN^60^. Briefly, Hotspot Wizard^61^ was run with *Gm*SuSy-97, with residues belonging to the catalytic active site selected as essential and excluded from mutagenesis (Table S20). Standard settings were applied and the maximum number of sequences was set to 10,000 to maximize sequence diversity. Based on a mutability score threshold (>6), 283 positions were identified as candidate hotspots. From these hotspots, three design strategies were implemented: Non-conservative strategy: All 283 positions were submitted to ProteinMPNN^47,62^ using three different sampling temperatures (T = 0.1, 0.2, and 0.3), and ten sequences were designed per sampling temperature. For each design sequence, AlphaFold2 using MMseqs2 structural models were generated^63,64^ and *in silico* characterization was performed based on sequence identity, structural similarity (RMSD) relative to WT, surface charge^59^, solubility score (Soluprot)^58^ and FoldX folding energies^65,66^. Based on a balanced evaluation of these parameters, one variant per temperature was selected for experimental validation, named SuSy-T1, SuSy-T2, and SuSy-T3 (Table S4). Unfortunately, none of these variants could be expressed and recovered, and SuSy activity could not be detected.

Conservative strategy: Residues located within 5 Å of amino acids at the active site were excluded from the 283 previously identified hotspots. Additionally, hotspots with mutability scores of 6 and 7 were removed, resulting in a final set of 62 positions subjected to ProteinMPNN. As in the non-conservative strategy, the top ten designs per sampling temperature were evaluated *in silico*. Most designs generated under this strategy were highly similar, thus, a consensus sequence was generated per sampling temperature. At positions without a clear consensus, manual inspection of proposed mutation was performed, and residues predicted to improve molecular interactions were selected. Due to the high sequence similarity among designs, a single variant was selected for experimental validation, designated as HPMN (Table S23). HPMN, carrying 47 single-point mutations (Table S4), showed around fivefold higher soluble expression than the previous variants, but had reduced Tm_app_ (−3.4 °C) and a 79% loss in activity compared to Anc165 (Tables S5-S7).

Conservative strategy + Ancestral sequence reconstruction: Based on these observations, we combined the sequences of HPMN and Anc165. Mutations derived from the HPMN variant were incorporated into Anc165. At positions where both Anc165 and HPMN differed from WT but encoded different amino acids, the Anc165 residue was retained. The resulting sequence was designated HPMN251.

### 4.5. Modified oligomeric interface variants

Mutations targeting the SuSy oligomer-oligomer interface were explored using a tetrameric structural model of WT. The PRODIGY web server (PROtein binDIng enerGY prediction)^67–69^ was used to estimate binding affinities and assess the impact of interface mutations on tetramer stability. Mutations predicted to enhance tetrameric interactions were designed through rational inspection of molecular interactions at the interface and by comparison with the crystal structure of *At*SuSy^38^. In addition, potential intermolecular disulfide bonds were designed using the Disulfide by Design 2 tool^70^. Selected single amino acid variants were generated and experimentally evaluated. Computational analyses are described in Table S24 and experimental results are provided in Tables S10 and S11.

### 4.6. Cloning, expression and protein purification

SuSy sequences were obtained already inserted into the expression vector pET28a+ at NcoI and XhoI sites. All sequences include a His-tag and a tobacco etch virus (TEV) site in the N-terminal. Desired amino acid mutations were introduced in the nucleic acid sequence encoding the SuSy enzyme by PCR and USER cloning^71^, and primers used are listed in Table S19. The resulting mutant sequences were verified by Sanger sequencing (Eurofins) after transformation into chemically competent *E. coli* BL21 Star (DE3).

5 mL pre-cultures of *E. coli* BL21 star (DE3) carrying the corresponding expression plasmid were grown overnight in 2xYT media containing kanamycin (50 μg mL^-1^) at 37 °C and 200 rpm. These pre-cultures were used to inoculate 500 mL of 2xYT media containing kanamycin (50 μg mL^-1^), which were grown at 37 °C and 200 rpm until reaching an optical density (OD) between 0.5 and 0.8. Protein expression was induced by adding isopropyl-β-D-thiogalactopyranoside (IPTG) at a final concentration of 0.2 mM and the cultures were incubated at 20 °C and 180 rpm for 16-20 h.

Cells were harvested by centrifugation at 4 °C and 4000 g for 15 min, and cell pellets were resuspended in buffer A (20 mM imidazole, 50 mM phosphate buffer, 300mM NaCl, and pH 7.5) containing DNase I (15 μg mL^-1^). The cell suspension was subjected to sonication with a VCX130 sonicator and a 6 mm probe (Sonics & Materials Inc.). An 85% amplitude was used, with a cycle of 30 seconds on and 30 seconds off, for a total sonication time of 6 minutes. During sonication all samples were placed in an ice bath. Sonicated samples were centrifuged at 12000-14000 g at 4 °C for 45 min and supernatants were filtered using a single-use filter unit of 0.45 µm.

Filtered supernatants were loaded onto a Histrap^TM^ FF 1mL column using an ÄKTA pure system (Cytiva). Briefly, columns were first equilibrated with buffer A prior to cell extract injection. Elution was carried out with a buffer B solution (500 mM imidazole, 50 mM phosphate buffer, 300 mM NaCl, and pH 7.5) using a linear gradient of buffer B from 2.5% to 60%. Fractions corresponding to a peak at 280 nm were analyzed using SDS-PAGE and fractions containing a band corresponding to SuSy molecular weight were pooled. To remove imidazole, buffer exchange was carried out with a 50,000 MWCO Amicon Ultra-15 Centrifugal Filter Unit (Sigma-Aldrich) using buffer C (50 mM phosphate buffer, 150 mM NaCl, and pH 7.5). The protein was concentrated, and its final concentration was determined using a Nanodrop spectrophotometer at 280 nm, applying the theoretical molecular weight and extinction coefficient. Concentrated proteins were aliquoted and stored at −70 °C until further use.

### 4.7. Apparent melting temperature determination

The apparent protein melting temperature was determined through DSF using the Protein Thermal Shift Dye kit (Thermo Fisher). Briefly, protein samples were prepared at a 2x concentration (0.8-1.0 mg mL^-1^) in reaction buffer (50 mM phosphate buffer, pH 7.5). A 50 µL aliquot was mixed with 50 µL of a 2x SYPRO™ orange dye solution, and 20 µL of the resulting solution was transferred to a MicroAmp optical 96-well reaction plate. Samples were incubated at 25 °C for 2 min and fluorescence measurement was recorded in a QuantStudio5 qPCR system as the temperature increased to 99 °C at a ramp rate of 0.05 °C/second. For ancestral sequences and ancestral derived variants, the Tm_app_ was quantified with nanoDSF in the Prometheus Panta system. Proteins were prepared at 1.0 mg mL^-1^ concentration in reaction buffer, and a 10 µL aliquot was transferred to a capillary tube. The program was run using an excitation power of 20% and a temperature slope of 1-2 °C per minute starting from 25 °C to 100 °C. Tm_app_ quantifications using DSF and nanoDSF were carried out by quadruplicates and triplicate, respectively.

DSF analysis of the WT displayed two distinct thermal transitions, and both were used during the mutant screening and combinatorial analysis. Final variants *Gm*SuSy-97, Anc165, and HPMN251 exhibited a single thermal transition. In the case of comparison between variants with single and double thermal transitions, the Tm_app_ of the second thermal transition was chosen.

Changes in Tm_app_ as a function of DMSO concentration were determined through nanoDSF using 1 mg/mL of protein with concentrations of DMSO varying from 5% to 80% (v/v) incubated at room temperature for approximately 1 h. The ratio of folded and unfolded tryptophan fluorescence (F350/F330) was plotted as a function of DMSO concentration at temperatures ranging from 25 °C to 70 °C as reported by Sorgenfrei et al., 2024^72^.

### 4.8. Relative and residual activity measurement

Relative activities were assessed by incubating 90 µL of a reaction solution (50 mM phosphate buffer, pH 7.5, 200 mM sucrose, and 1 mM UDP) with 10 µL of a 1 µM protein sample (0.1 µM final concentration). *Ac*SuSy_M_ activity assay was carried out with 1 µM final concentration. Reactions were conducted at 30 °C for 5 minutes and terminated by heat inactivation at 95 °C. The formation of fructose, as a reaction product, was quantified using the D-Fructose/D-Glucose Assay Kit (Megazyme) following the manufacturer’s protocol, with a fructose standard curve ranging from 0 to 2 µg (Figure S22). One unit of specific activity (U mg^-1^) was defined as 1 µmol of fructose produced per minute per mg of total protein, and relative activity was calculated by normalizing to the specific activity of the WT, which was set at 100%.

Residual activity was assessed similarly, except that each protein sample was incubated for 5 min at 60 °C prior to addition to the reaction mixture. Residual activity was expressed relative to the activity of the WT under non-heat-treated conditions. Residual activity reflects the kinetic stability of the protein, indicating its enzymatic activity after heat exposure. This parameter is compared to the non-heated WT activity to facilitate the identification of mutants with enhanced kinetic stability. All relative and residual activity quantifications were carried out by triplicate and considered a blank consisting of all reaction components except for enzyme, and storage/reaction buffer was used instead.

### 4.9. Kinetic parameters, solvent tolerance and soluble expression determination

All specific activity measurements for SuSy variants characterization were determined with fructose quantification as described above. Reaction times (1 to 3 min) and enzyme concentrations (0.01 to 0.05 µM) were adjusted to maintain initial-rate conditions. All experiments were carried out by triplicate.

Optimal pH values were determined by measuring specific activity using buffers prepared at a final concentration of 50 mM, including citrate buffer (pH 5.0-5.5) and phosphate buffer (pH 6.0-8.0). Temperature optima (20-80 °C) were subsequently determined at the optimal pH identified for each enzyme variant (Figure 1e; Figure S23).

Kinetic stability was assessed under optimal conditions established for each variant. Briefly, 50 µL of each SuSy variant (0.05 mg mL^-1^) diluted in their corresponding optimal pH buffer were incubated at 55 and 60 °C for different time intervals. After incubation, the aliquots were placed at 4 °C, spun, and specific activity was quantified at the optimal temperature. Residual activity over time was fitted to one-phase decay or plateau followed by on-phase decay models using GraphPad Prism v10.6.0 to determine half-life (t_1/2_) and thermal deactivation constants (k_d_) (Figures S5 and S6).

Kinetic parameters were determined from sucrose cleavage reactions conducted at the optimal pH and temperature for each SuSy variant using sucrose concentrations from 25 - 1000 mM. Data were fitted to the Michaelis-Menten model using GraphPad Prism v10.6.0 (Figure S7). As substrate inhibition was observed at high sucrose concentrations, only pre-inhibition data were used to estimate K_M_ and k_cat_ apparent values. Blanks without UDP and with enzyme were also performed to account for sucrose hydrolysis under increasing sucrose concentration.

Solvent tolerance was evaluated by measuring specific activity under optimal conditions in the presence of 5-25% (v/v) DMSO, methanol (MeOH), and acetonitrile (ACN) (Table S25). For MeOH and ACN, reactions were initiated on ice by enzyme addition and subsequently transferred to the optimal temperature to minimize solvent loss by volatilization.

### 4.10. Molecular dynamics simulations

The initial structures for all molecular dynamics (MD) simulations were derived from Protenix web server^73,74^, generated as tetramers for WT, *Gm*SuSy-97, and Anc165 in complex with sucrose and UDP. Ligand binding poses were validated against the ligands-bound crystal structures of *At*SuSy (PDBs: 3S29 and 3S28)^38^. Protonation states of titratable residues were first predicted at pH 6.0 using H++^75^ and PropKa^76,77^, and subsequently refined by manual inspection of local hydrogen-bonding patterns and electrostatic microenvironments following short 50-ns MD simulations. Final protonation assignments, including Asp, Glu, and His residues displaying atypical protonation states, are reported in Table S26. All Lys and Arg side chains remained protonated. Each protein-ligand complex was solvated in an OPC^78^ water truncated cubic box providing a 12 Å buffer, and neutralized with counterions. The FF19SB^79^ force field (for proteins), GLYCAM06^80^ (for sucrose), and GAFF2^81^ (for UDP) were applied.

System construction was performed in AMBER 22 using the LEaP^82^ module, and all subsequent MD simulations employed the CUDA-accelerated PMEMD engine in AMBER 24^83^. Energy minimization proceeded in two stages^84^: 2,500 steps of steepest descent followed by 2,500 steps of conjugate-gradient descent, with solute heavy atoms restrained by a harmonic potential of 50 kcal mol⁻¹ Å⁻². The systems were then heated from 0 to 333.15 K (60 °C) using the Berendsen thermostat (1.0 ps coupling constant), followed by a series of five 500 ps equilibration stages under NPT conditions (1 atm) during which positional restraints on solute heavy atoms were linearly tapered to 5 kcal mol⁻¹ Å⁻². Production simulations consisted of three independent 1.2 μs NPT trajectories at 333.15 K and 1 atm. Temperature control employed a Langevin thermostat (2.0 ps⁻¹ collision frequency) and pressure control used a Berendsen barostat (1.0 ps coupling constant). A 2.0-fs timestep was enabled by constraining bonds to hydrogen with SHAKE^85^ algorithm. Long-range electrostatics were treated using the particle-mesh Ewald (PME)^86^ method, applying a 9 Å cutoff for nonbonded interactions. Snapshots were recorded every 20 ps, and all production runs were performed without positional restraints. All properties of the protein complexes were quantified using the CPPTRAJ module in AMBER 22^87^. Hydrogen bonds were computed using a donor-acceptor distance cutoff of 3.9 Å and an angle cutoff of 90°, following the suggested criteria^88^. Salt bridges were identified using a distance cutoff of 4.0 Å, hydrophobic contacts using a cutoff of 5.0 Å, and SASA was calculated using default parameters.

Clusters were generated with TTClust v4.10.4^89^ using default settings. The binding energies across the identified clusters were calculated using MM/PBSA in AMBER^83^ based on a total of 14.4 µs of sampling of monomeric complex conformations. Dynamic correlation analysis was carried out using the SPM method^51,52^. SPMs were calculated with significance thresholds of 0.1 and a distance cutoff of 6.0 Å. SPMs were reconstructed from three independent MD replicates of 1.2 µs each and from the combined 3.6 µs ensemble. Electrostatic potentials were calculated using the APBS webserver^90^ with standard settings at pH 5.5 and 7.5 for Anc165 and at pH 6.0 and 7.5 for WT.

### 4.11. MANT glycosylation

SuSy variants (WT, *Gm*SuSy-97, Anc165, and HPMN251) were evaluated for MANT glycosylation in a coupled cascade with UGT72B68 from *Solanum lycopersicum* (UGT16; UniProt D7S016)^31^. Screening reactions (final volume of 100 µL) contained 200 mM sucrose, 50 mM phosphate buffer (pH 7.5), 0.351 mM UDP, 1 µM UGT16, and 0.75 µM SuSy. Reactions were initiated by addition of MANT dissolved in DMSO, resulting in a final concentration of 20 mM MANT and 10% (v/v) DMSO. Reactions were incubated at 40 °C and sampled after 1, 2, 3, 4, 5, and 22 h. Reactions were quenched by heating at 95 °C for 1 min, diluted with Milli-Q water, and analyzed by HPLC for quantification of MANT and MANT-*N*-glucose relative peaks as described in Gharabli *et al* (2025)^31^ (Figure S24).

Reaction optimization was performed using HPMN251. An experimental design was first applied to evaluate the effects of sucrose concentration (100-300 mM), pH (50 mM phosphate buffer 7.5-8.5), UDP concentration (0.125-0.5 mM), temperature (35-45 °C), and the UGT:SuSy molar ratio (1-2, with UGT fixed at 1 µM). A 2 full factorial design with center points was employed with triplicate, and reactions were stopped after 5 h. Substrate conversion (%) was used as the response variable. All factors were found to significantly influence conversion, although the UGT:SuSy ratio showed only a minor effect (Table S27; Figure S17).

Based on the optimization results, selected reaction conditions (300 mM sucrose, 50 mM phosphate buffer pH 7.0, 0.5 mM UDP, 45 °C) were used to further evaluate the influence of enzyme loading by varying UGT concentration (1.2, 1.5, and 2.0 µM) and UGT:SuSy ratios (5 and 10) (Figure S17) after 20 h incubation. Benchmarking with *Ac*SuSy_M_ under optimal conditions was also carried out.

Whole-cell reactions for MANT glycosylation were performed under optimized conditions, substituting purified proteins with *E. coli* BL21 star (DE3) wet biomass expressing UGT or SuSy. The biomass was prepared as previously described, but after centrifugation, it was resuspended at either 100 or 150 g L^-1^ and frozen at −70 °C. To ensure that the protein concentration in the biomass matched the enzyme concentration used in the purified form, an SDS gel with biomass and a standard curve generated with purified protein was run. Bands corresponding to the molecular sizes of UGT and SuSy in the cells were quantified using an iBright 1500 imaging system (Invitrogen) and data analysis was conducted with Image Lab v6.1 (Biorad) (Figure S25).

All reactions were performed with triplicate, and blanks without enzyme were run to verify effects of reaction conditions on conversion. Additionally, blanks with whole cell biomass lacking either UGT or SuSy were considered to account for side reactions.

### 4.12. Indoxyl glycosylation

WT, Anc165, and HPMN251 were evaluated for glycosylation of indoxyl yielding indoxyl-β-glycoside (Indican) in a three-enzyme cascade with stabilized *Pt*UGT1^stable^ ^6^ from *Polygonum tinctorium* and esterase from porcine liver (15 U/mg from Sigma-Aldrich).

Cascade reaction optimization using 100 mM indoxyl acetate was carried out through a 2^4^ central composite design with five levels using Anc165. Effects of sucrose concentration (50-600 mM), phosphate-citrate buffer (pH 5-8), temperature (30-60 °C) and UDP concentration (0.001-2 mM) were tested with a fixed UGT:SuSy 1:1 mass ratio of 50 µg (Figure S26). The reaction set-up was adapted from Bidart et al. (2024)^6^, using a final volume of 400 µL. The reaction was initiated by adding 4U of esterase, and it was incubated at the corresponding temperatures under magnetic stirring (100 rpm) and anaerobic conditions. After 6 h, the reaction was quenched by incubation at 95 °C for 1 min, and subsequently diluted for HPLC indican quantification (Figure S24). Samples were analyzed at 270 nm using a gradient of acetonitrile (B) with 0.1% TFA in RP-HPLC (1 min at 15% B, ramp up to 35% B at 6 min, increase to 85% B at 12 min, 100% B at 13 min, hold at 100% B until 14 min, decrease to 5% B at 15.1 min, and re-equilibrated until 18 min), with a flow rate of 1 mL/min. The full design was performed with biological triplicate (Table S15).

Based on optimization results, selected reaction conditions (100 mM indoxyl acetate, 600 mM sucrose, 90 mM citrate phosphate buffer pH 8.0, 2 mM UDP and 54 °C) were used to assess a 1:5 UGT:SuSy mass ratio for Anc165 and HPMN251 in triplicate. Comparison with WT was also run using 90 mM phosphate-citrate buffer (pH 8.0), 1 mM UDP, 200 mM sucrose, 50 µg *Pt*UGT1-87 and 50 µg SuSy.

### 4.13. Preliminary techno-economic and life-cycle assessment

Preliminary life cycle assessment (LCA) and techno-economic analysis (TEA) were performed to evaluate the reductions on the environmental impacts and costs associated with the implementation of robust SuSy in the two study cases. For MANT glycosylation, the comparative analyses were performed using HPMN251 and WT from this study, and UDP-Glc system from Gharabli et al., (2025)^31^. For indoxyl glycosylation the comparison was based between WT, HPMN251 and UDP-Glc systems presented this study, and the conversion achieved by Bidart et al., (2024)^6^.

The functional unit for each study case was defined as the production of 1 kg of MANT-*N*-glucose and 1 kg indican, respectively. System boundaries were defined as cradle-to-gate and limited to the glycosylation reaction step, including upstream synthesis of UDP and UDP-Glc^31^ (Figures S18 and S26). Energy inputs, infrastructure, solvent recovery, and downstream purification of intermediates and products were excluded from the system boundary, consistent with the early-stage nature of the process.

The life cycle inventory was constructed from reaction mass balances from reported studies and from experimental data generated here. Mass balances and assumptions are provided in Tables S12-S14 and S16-S18. Life cycle impact assessment was performed using openLCA v2.1 with background data sourced from the ecoinvent v3.8 database. Environmental impacts were calculated using the ReCiPe 2016 Endpoint (H) method^91^. Normalization of endpoint categories was done using World (2010) H/H as weighting set. The normalization allows direct comparison by assigning a single score (Pt) to each system. Given the preliminary scope and laboratory-scale data used, results are intended to highlight relative differences between scenarios, rather than provide definitive environmental footprints.

The reaction costs for 1 kg of MANT-N-glucose and 1 kg indican were estimated using indicative bulk market prices obtained from industrial suppliers (Alibaba) and complemented with literature-reported values for the components included in the reaction mass balance (Tables S13 and S17). Unit prices were set to 10 USD kg⁻¹ for MANT^92^, 5 USD kg⁻¹ for indoxyl acetate^6^, 0.5 USD kg⁻¹ for sucrose^93^, 84.9 USD kg^-1^ for UDP-Glc^6^, 50 USD kg⁻¹ for UDP^6^, 0.76 USD kg⁻¹ for phosphate buffer^94^, 0.1 USD kg⁻¹ for citrate-phosphate buffer^6^, 25 USD kg⁻¹ for enzymes^6^, 1.5 USD kg⁻¹ for DMSO^95^, and 0.0077 USD kg⁻¹ for water^6^.

### 4.14. Statistical analysis

All measurements were performed by triplicate unless otherwise specified. Statistical significance of reaction optimization and comparison between samples was evaluated at a 5% significance level using analysis of variance (ANOVA), followed by Tukey’s post hoc test. Normality and homoscedasticity assumptions were verified for reaction optimization and statistical analyses were performed using R v4.5.2^96^, RStudio v2026.01.0 Build 392^97^ and GraphPad Prism v10.6.0.

## Author contributions

FMO designed the enzyme engineering strategies and performed screening, variant characterization, process optimization for methyl anthranilate glycosylation, and preliminary techno-economic and life-cycle assessments with contributions from BML, CM, GB, AM, AB, MEG, and DW. Indoxyl glycosylation optimization was carried out by MH with assistance from FMO and DW. Molecular dynamics studies were led by DD and CR, with contributions from FMO and DW. The project was conceptualized by FMO and DW. Finally, FMO and DW wrote the manuscript with input from all authors.

## Acknowledgements

This work was supported by The Novo Nordisk Foundation through grants NNF20CC0035580, NNF24SA0100980, and NNF24OC0088001. FMO received funding from the Copenhagen Bioscience Ph.D. Programme (Grant No. NNF22SA0078231). DD acknowledges the support of la Caixa foundation: The project that gave rise to these results received the support of a fellowship from “la Caixa” Foundation (ID 100010434). The fellowship code is LCF/BQ/DR24/12080011.” The authors express gratitude to Alexander Kai Büll and Lars Knuth Satoshi Boyens-Thiele for training and assistance with the use of the Prometheus Panta (NanoTemper). The authors thank David Harding-Larsen, Ruben de Boer, Hani Gharabli, Onur Kırtel, Carlos Lozano, Patricia Arias, Maria Isabel Rodriguez-Torres and John M. Woodley for their support and guidance during the course of this work.

## References

1. Mrudulakumari Vasudevan, U. & Lee, E. Y. Flavonoids, terpenoids, and polyketide antibiotics: Role of glycosylation and biocatalytic tactics in engineering glycosylation. Biotechnology Advances 41, 107550 (2020).

2. Pifferi, C., Fuentes, R. & Fernández-Tejada, A. Natural and synthetic carbohydrate-based vaccine adjuvants and their mechanisms of action. Nat Rev Chem 5, 197– 216 (2021).

3. Reed, J. et al. Elucidation of the pathway for biosynthesis of saponin adjuvants from the soapbark tree. Science 379, 1252–1264 (2023).

4. Huang, W. et al. Complete pathway elucidation of echinacoside in *Cistanche tubulosa* and de novo biosynthesis of phenylethanoid glycosides. Nat Commun 16, 882 (2025).

5. Wang, Y. et al. Characterization of *Sr* UGT76G4 reveals a key residue for regioselectivity and efficient Reb M synthesis. Proc. Natl. Acad. Sci. U.S.A. 122, e2504698122 (2025).

6. Bidart, G. N. et al. Chemoenzymatic indican for light-driven denim dyeing. Nat Commun 15, 1489 (2024).

7. Pinkas, Z. et al. Glycosylated cannabinoids in *Cannabis sativa* and enzyme design to modulate their synthesis. Proc. Natl. Acad. Sci. U.S.A. 122, e2515688122 (2025).

8. Liu, D. et al. Rational design of a bifunctional glycosyltransferase for enhanced substrate promiscuity and thermostability. Journal of Biotechnology 410, 285–297 (2026).

9. Go, S.-R., Lee, S.-J., Ahn, W.-C., Park, K.-H. & Woo, E.-J. Enhancing the thermostability and activity of glycosyltransferase UGT76G1 via computational design. Commun Chem 6, 265 (2023).

10. Li, G. et al. Engineering C-glycosyltransferase UGT708A60 to enhance the activity and thermostability for the highly efficient biosynthesis of nothofagin. Food Bioscience 74, 107952 (2025).

11. Ding, J. et al. Semi-rational design of a thermostable *O*-glycosyltransferase from *Glycyrrhiza uralensis* for efficient conversion of protopanaxadiol. Journal of Biotechnology 410, 217–227 (2026).

12. Kırtel, O., Strother, L. H., Putkaradze, N. & Welner, D. H. Halophytic C-Glycosyltransferases Enable C-Glycosylation in Organic Solvents. ACS Omega 10, 55909–55919 (2025).

13. Wang, H. et al. Characterization of an organic solvent tolerance C-glucosyltransferase from *Phyllostachys edulis* and development of a whole-cell biocatalyst for the efficient and sustainable production of Nothofagin. Molecular Catalysis 584, 115308 (2025).

14. Schmölzer, K., Gutmann, A., Diricks, M., Desmet, T. & Nidetzky, B. Sucrose synthase: A unique glycosyltransferase for biocatalytic glycosylation process development. Biotechnology Advances 34, 88–111 (2016).

15. Mejia-Otalvaro, F., Lax, B. M., Kırtel, O. & Welner, D. H. Sustainable Natural Product Glycosylation: A Critical Evaluation of Biocatalytic and Chemical Approaches. ChemSusChem 18, e202501094 (2025).

16. Klotz, K. L., Finger, F. L. & Shelver, W. L. Characterization of two sucrose synthase isoforms in sugarbeet root. Plant Physiology and Biochemistry 41, 107–115 (2003).

17. Morell, M. & Copeland, L. Sucrose Synthase of Soybean Nodules. Plant Physiol 78, 149–154 (1985).

18. Baroja-Fernández, E. et al. Sucrose synthase activity in the sus1/sus2/sus3/sus4 *Arabidopsis* mutant is sufficient to support normal cellulose and starch production. Proceedings of the National Academy of Sciences 109, 321–326 (2012).

19. Liu, H., Tegl, G. & Nidetzky, B. Glycosyltransferase Co-Immobilization for Natural Product Glycosylation: Cascade Biosynthesis of the *C*-Glucoside Nothofagin with Efficient Reuse of Enzymes. Adv Synth Catal 363, 2157–2169 (2021).

20. Elling, L. & Kula, M.-R. Characterization of sucrose synthase from rice grains for the enzymatic synthesis of UDP and TDP glucose. Enzyme and Microbial Technology 17, 929–934 (1995).

21. Sebkova, V., Unger, C., Hardegger, M. & Sturm, A. Biochemical, Physiological, and Molecular Characterization of Sucrose Synthase from *Daucus carota*. Plant Physiol. 108, 75–83 (1995).

22. Diricks, M. Unlocking nature’s glycosylation potential: characterization and engineering of novel sucrose/trehalose synthases. (Ghent University, 2017).

23. Diricks, M., De Bruyn, F., Van Daele, P., Walmagh, M. & Desmet, T. Identification of sucrose synthase in nonphotosynthetic bacteria and characterization of the recombinant enzymes. Appl Microbiol Biotechnol 99, 8465–8474 (2015).

24. Chen, K. et al. Identification of sucrose synthase from *Micractinium conductrix* to favor biocatalytic glycosylation. Front. Microbiol. 14, 1220208 (2023).

25. Zhang, L. et al. Mining of Sucrose Synthases from *Glycyrrhiza uralensis* and Their Application in the Construction of an Efficient UDP-Recycling System. J. Agric. Food Chem. 67, 11694–11702 (2019).

26. Nidetzky, B. Glycosyltransferase Cascades Made Fit For the Biocatalytic Production of Natural Product Glycosides. in Biocatalysis for Practitioners (eds De Gonzalo, G. & Lavandera, I.) 225–243 (Wiley, 2021). doi:10.1002/9783527824465.ch8.

27. Bungaruang, L., Gutmann, A. & Nidetzky, B. β-Cyclodextrin Improves Solubility and Enzymatic C-Glucosylation of the Flavonoid Phloretin. Advanced Synthesis & Catalysis 358, 486–493 (2016).

28. Zhang, S., Zhao, L., Li, T., Ding, Z. & Dong, W. Bioinspired interfacial assembly of amorphous ZIF nanoarmor for robust direct immobilization of crude sucrose synthase. Colloids and Surfaces B: Biointerfaces 265, 115704 (2026).

29. â-Cyclodextrin Improves Solubility and Enzymatic C-Glucosylation of the Flavonoid Phloretin. Advanced Synthesis and Catalysis 358, 486–493 (2016).

30. Schmölzer, K., Lemmerer, M. & Nidetzky, B. Glycosyltransferase cascades made fit for chemical production: Integrated biocatalytic process for the natural polyphenol C-glucoside nothofagin. Biotechnology and Bioengineering 115, 545–556 (2018).

31. Gharabli, H. et al. Enzymatic Glycosylation of Anthranilates for Enhanced Functionality. 2025.05.29.656899 Preprint at 10.1101/2025.05.29.656899 (2025).

32. US20210054349A1 - Engineered glycosyltransferases and steviol glycoside glucosylation methods - Google Patents. https://patents.google.com/patent/US20210054349A1/en?q=(Sucrose+synthase)&assignee=codexis&oq=Sucrose+synthase+codexis.

33. Zhao, L. et al. Engineering the Thermostability of Sucrose Synthase by Reshaping the Subunit Interaction Contributes to Efficient UDP-Glucose Production. J. Agric. Food Chem. 71, 3832–3841 (2023).

34. Zhao, L. et al. Highly Efficient Production of UDP-Glucose from Sucrose via the Semirational Engineering of Sucrose Synthase and a Cascade Route Design. J. Agric. Food Chem. 71, 12549–12557 (2023).

35. Zhao, L. et al. Synthesis of value-added uridine 5’-diphosphate-glucose from sucrose applying an engineered sucrose synthase counteracts the activity-stability trade-off. Food Chemistry 464, 141765 (2025).

36. Chen, K. et al. Identification and engineering of a sucrose synthase from *Stevia rebaudiana* for glycosylation applications. Journal of Biotechnology 405, 169–181 (2025).

37. Wu, R. et al. The Crystal Structure of *Nitrosomonas europaea* Sucrose Synthase Reveals Critical Conformational Changes and Insights into Sucrose Metabolism in Prokaryotes. Journal of Bacteriology 197, 2734–2746 (2015).

38. Zheng, Y., Anderson, S., Zhang, Y. & Garavito, R. M. The Structure of Sucrose Synthase-1 from *Arabidopsis thaliana* and Its Functional Implications. Journal of Biological Chemistry 286, 36108–36118 (2011).

39. Nakai, T. et al. An Increase in Apparent Affinity for Sucrose of Mung Bean Sucrose Synthase Is Caused by In Vitro Phosphorylation or Directed Mutagenesis of Ser11. Plant Cell Physiol 39, 1337–1341 (1998).

40. Duncan, K. A. & Huber, S. C. Sucrose Synthase Oligomerization and F-actin Association are Regulated by Sucrose Concentration and Phosphorylation. Plant and Cell Physiology 48, 1612–1623 (2007).

41. Sauerzapfe, B., Engels, L. & Elling, L. Broadening the biocatalytic properties of recombinant sucrose synthase 1 from potato (*Solanum tuberosum* L.) by expression in *Escherichia coli* and *Saccharomyces cerevisiae*. Enzyme and Microbial Technology 43, 289–296 (2008).

42. Chebotareva, N. A., Roman, S. G. & Kurganov, B. I. Dissociative mechanism for irreversible thermal denaturation of oligomeric proteins. Biophys Rev 8, 397–407 (2016).

43. Unsworth, L. D., van der Oost, J. & Koutsopoulos, S. Hyperthermophilic enzymes − stability, activity and implementation strategies for high temperature applications. The FEBS Journal 274, 4044–4056 (2007).

44. Gutmann, A., Lepak, A., Diricks, M., Desmet, T. & Nidetzky, B. Glycosyltransferase cascades for natural product glycosylation: Use of plant instead of bacterial sucrose synthases improves the UDP-glucose recycling from sucrose and UDP. Biotechnol. J. 12, 1600557 (2017).

45. Diricks, M. et al. Sequence determinants of nucleotide binding in Sucrose Synthase: improving the affinity of a bacterial Sucrose Synthase for UDP by introducing plant residues. *Protein Engineering*, Design and Selection proeng; gzw048v1 (2016) doi:10.1093/protein/gzw048.

46. Livada, J., Vargas, A. M., Martinez, C. A. & Lewis, R. D. Ancestral Sequence Reconstruction Enhances Gene Mining Efforts for Industrial Ene Reductases by Expanding Enzyme Panels with Thermostable Catalysts. ACS Catal. 13, 2576–2585 (2023).

47. Dauparas, J. et al. Robust deep learning–based protein sequence design using ProteinMPNN. Science 378, 49–56 (2022).

48. Nidetzky, B., Gutmann, A. & Zhong, C. Leloir Glycosyltransferases as Biocatalysts for Chemical Production. ACS Catal. 8, 6283–6300 (2018).

49. Cui, H. et al. How to Engineer Organic Solvent Resistant Enzymes: Insights from Combined Molecular Dynamics and Directed Evolution Study. ChemCatChem 12, 4073–4083 (2020).

50. Ortega-Quintanilla, G. & Millet, O. On the Molecular Basis of the Hypersaline Adaptation of Halophilic Proteins. Journal of Molecular Biology 438, 169439 (2026).

51. Casadevall, G., Casadevall, J., Duran, C. & Osuna, S. The shortest path method (SPM) webserver for computational enzyme design. Protein Eng Des Sel 37, gzae005 (2024).

52. Osuna, S. The challenge of predicting distal active site mutations in computational enzyme design. WIREs Computational Molecular Science 11, e1502 (2021).

53. Luo, Z. W., Cho, J. S. & Lee, S. Y. Microbial production of methyl anthranilate, a grape flavor compound. Proc. Natl. Acad. Sci. U.S.A. 116, 10749–10756 (2019).

54. Avery, M. L. et al. Methyl Anthranilate as a Rice Seed Treatment to Deter Birds. The Journal of Wildlife Management 59, 50–56 (1995).

55. Gumulya, Y. et al. Engineering highly functional thermostable proteins using ancestral sequence reconstruction. Nat Catal 1, 878–888 (2018).

56. Musil, M. et al. FireProt: web server for automated design of thermostable proteins. Nucleic Acids Res 45, W393–W399 (2017).

57. Musil, M. et al. FireProtASR: A Web Server for Fully Automated Ancestral Sequence Reconstruction. Briefings in Bioinformatics 22, bbaa337 (2021).

58. Hon, J. et al. SoluProt: prediction of soluble protein expression in *Escherichia coli*. Bioinformatics 37, 23–28 (2021).

59. Jurrus, E. et al. Improvements to the APBS biomolecular solvation software suite. Protein Science 27, 112–128 (2018).

60. Sumida, K. H. et al. Improving Protein Expression, Stability, and Function with ProteinMPNN. J. Am. Chem. Soc. 146, 2054–2061 (2024).

61. Sumbalova, L., Stourac, J., Martinek, T., Bednar, D. & Damborsky, J. HotSpot Wizard 3.0: web server for automated design of mutations and smart libraries based on sequence input information. Nucleic Acids Res 46, W356–W362 (2018).

62. Dürr, S. L. ProteinMPNN Gradio Webapp. Zenodo 10.5281/zenodo.7630417 (2023).

63. Jumper, J. et al. Highly accurate protein structure prediction with AlphaFold. Nature 596, 583–589 (2021).

64. Mirdita, M. et al. ColabFold: making protein folding accessible to all. Nat Methods 19, 679–682 (2022).

65. Van Durme, J. et al. A graphical interface for the FoldX forcefield. Bioinformatics 27, 1711–1712 (2011).

66. Krieger, E., Koraimann, G. & Vriend, G. Increasing the precision of comparative models with YASARA NOVA—a self-parameterizing force field. Proteins: Structure, Function, and Bioinformatics 47, 393–402 (2002).

67. Xue, L. C., Rodrigues, J. P., Kastritis, P. L., Bonvin, A. M. & Vangone, A. PRODIGY: a web server for predicting the binding affinity of protein–protein complexes. Bioinformatics 32, 3676–3678 (2016).

68. Vangone, A. & Bonvin, A. M. Contacts-based prediction of binding affinity in protein– protein complexes. eLife 4, e07454 (2015).

69. Honorato, R. V. et al. Structural Biology in the Clouds: The WeNMR-EOSC Ecosystem. Front. Mol. Biosci. 8, (2021).

70. Craig, D. B. & Dombkowski, A. A. Disulfide by Design 2.0: a web-based tool for disulfide engineering in proteins. BMC Bioinformatics 14, 346 (2013).

71. Bitinaite, J. et al. USER^TM^ friendly DNA engineering and cloning method by uracil excision. Nucleic Acids Res 35, 1992–2002 (2007).

72. Sorgenfrei, F. A. et al. Solvent concentration at 50% protein unfolding may reform enzyme stability ranking and process window identification. Nat Commun 15, 5420 (2024).

73. Team, P. et al. PXDesign: Fast, Modular, and Accurate De Novo Design of Protein Binders. 2025.08.15.670450 Preprint at 10.1101/2025.08.15.670450 (2025).

74. Team, B. A. A. et al. Protenix - Advancing Structure Prediction Through a Comprehensive AlphaFold3 Reproduction. 2025.01.08.631967 Preprint at 10.1101/2025.01.08.631967 (2025).

75. Gordon, J. C. et al. H++: a server for estimating pKas and adding missing hydrogens to macromolecules. Nucleic Acids Res 33, W368–371 (2005).

76. Søndergaard, C. R., Olsson, M. H. M., Rostkowski, M. & Jensen, J. H. Improved Treatment of Ligands and Coupling Effects in Empirical Calculation and Rationalization of pKa Values. J Chem Theory Comput 7, 2284–2295 (2011).

77. Olsson, M. H. M., Søndergaard, C. R., Rostkowski, M. & Jensen, J. H. PROPKA3: Consistent Treatment of Internal and Surface Residues in Empirical pKa Predictions. J. Chem. Theory Comput. 7, 525–537 (2011).

78. Izadi, S., Anandakrishnan, R. & Onufriev, A. V. Building Water Models: A Different Approach. J. Phys. Chem. Lett. 5, 3863–3871 (2014).

79. Tian, C. et al. ff19SB: Amino-Acid-Specific Protein Backbone Parameters Trained against Quantum Mechanics Energy Surfaces in Solution. J. Chem. Theory Comput. 16, 528–552 (2020).

80. Kirschner, K. N. et al. GLYCAM06: A generalizable biomolecular force field. Carbohydrates. Journal of Computational Chemistry 29, 622–655 (2008).

81. Wang, J., Wolf, R. M., Caldwell, J. W., Kollman, P. A. & Case, D. A. Development and testing of a general amber force field. Journal of Computational Chemistry 25, 1157–1174 (2004).

82. Case, D. A. et al. AmberTools. J. Chem. Inf. Model. 63, 6183–6191 (2023).

83. Salomon-Ferrer, R., Götz, A. W., Poole, D., Le Grand, S. & Walker, R. C. Routine Microsecond Molecular Dynamics Simulations with AMBER on GPUs. 2. Explicit Solvent Particle Mesh Ewald. J. Chem. Theory Comput. 9, 3878–3888 (2013).

84. Taleb, V. et al. Structural and mechanistic insights into the cleavage of clustered O-glycan patches-containing glycoproteins by mucinases of the human gut. Nat Commun 13, 4324 (2022).

85. Ryckaert, J.-P., Ciccotti, G. & Berendsen, H. J. C. Numerical integration of the cartesian equations of motion of a system with constraints: molecular dynamics of *n*-alkanes. Journal of Computational Physics 23, 327–341 (1977).

86. Darden, T., York, D. & Pedersen, L. Particle mesh Ewald: An N⋅log(N) method for Ewald sums in large systems. J. Chem. Phys. 98, 10089–10092 (1993).

87. Roe, D. R. & Cheatham, T. E. I. PTRAJ and CPPTRAJ: Software for Processing and Analysis of Molecular Dynamics Trajectory Data. J. Chem. Theory Comput. 9, 3084– 3095 (2013).

88. Torshin, I. Y., Weber, I. T. & Harrison, R. W. Geometric criteria of hydrogen bonds in proteins and identification of ‘bifurcated’ hydrogen bonds. *Protein Engineering*, Design and Selection 15, 359–363 (2002).

89. Tubiana, T., Carvaillo, J.-C., Boulard, Y. & Bressanelli, S. TTClust: A Versatile Molecular Simulation Trajectory Clustering Program with Graphical Summaries. J. Chem. Inf. Model. 58, 2178–2182 (2018).

90. Jurrus, E. et al. Improvements to the APBS biomolecular solvation software suite. Protein Science 27, 112–128 (2018).

91. Huijbregts, M. A. J. et al. ReCiPe2016: a harmonised life cycle impact assessment method at midpoint and endpoint level. Int J Life Cycle Assess 22, 138–147 (2017).

92. Farwell Cas#134-20-3 Methyl Anthranilate 98% Min. www.alibaba.com https://www.alibaba.com/product-detail/FARWELL-CAS-134-20-3-METHYL_60614625631.html.

93. Sugar - Agriculture and rural development - European Commission. https://agriculture.ec.europa.eu/data-and-analysis/markets/price-data/price-monitoring-sector/sugar_en (2025).

94. Water Treatment Chemicals Disodium Hydrogen Phosphate Na2hpo4 99% Industrial Dsp Sodium Acid Phosphate Kexing Brand 1kg/25kg. www.alibaba.com https://www.alibaba.com/product-detail/Water-Treatment-Chemicals-Disodium-Hydrogen-Phosphate_1601368211594.html.

95. Organic Intermediate Top Quality Dimethyl Sulfoxide Cas 67-68-5 Usp & Industrial Grade Dmso 99.99%. www.alibaba.com https://www.alibaba.com/product-detail/Organic-Intermediate-Top-Quality-Dimethyl-Sulfoxide_1601630635199.html.

96. R Core Team. R: A Language and Environment for Statistical Computing. (2025).

97. Posit team. RStudio: Integrated Development Environment for R. Posit Software, PBC (2025).

