## Supplementary Figures for "Efficient and low-impact enzymatic glycosylation with robust sucrose synthase variants"

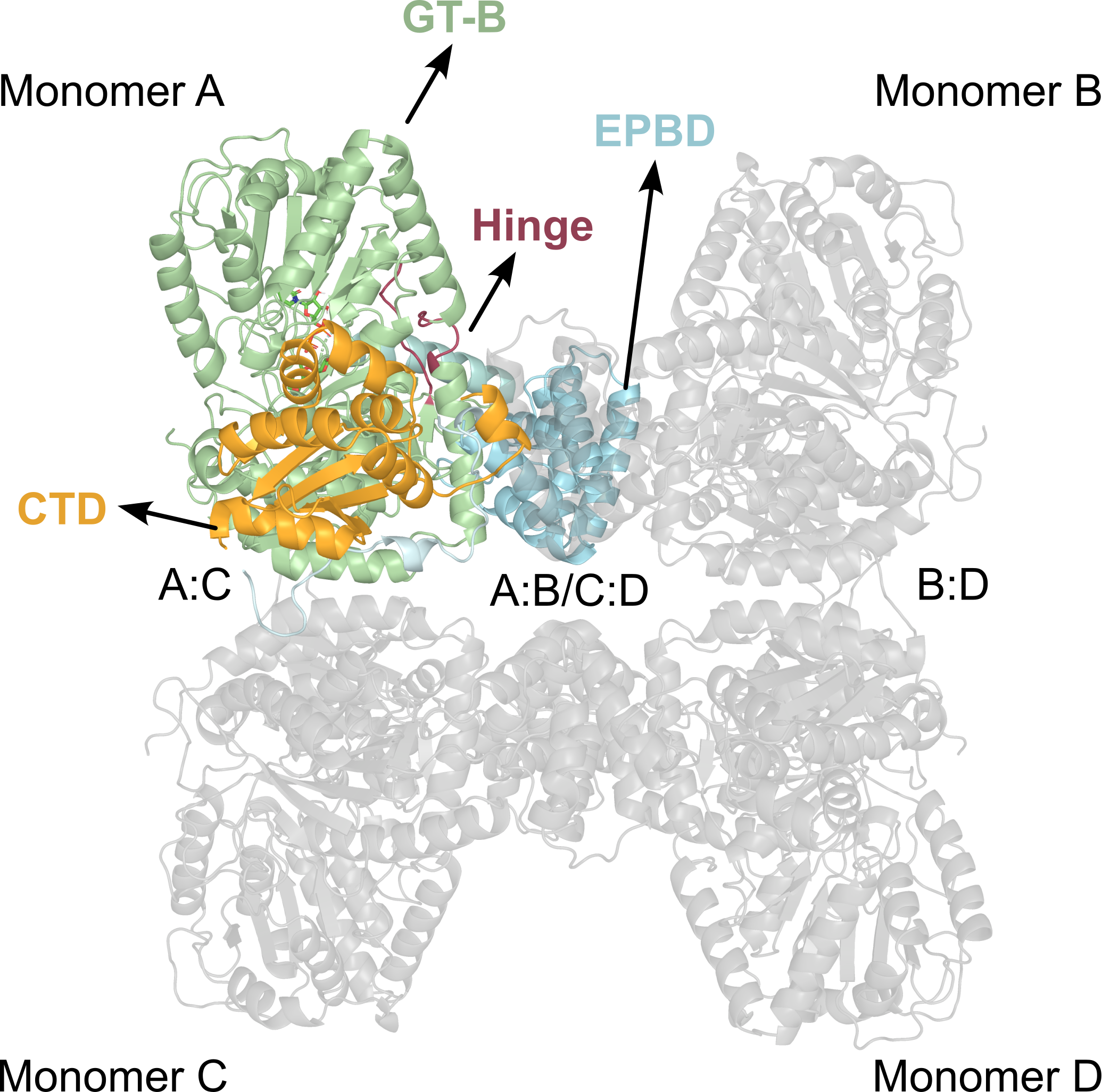

**Figure S1.** Domain organization of SuSy. In plant SuSys, the cellular targeting (CTD) and ENOD40 peptide-binding (EPBD) domains form the tetramer interfaces, whereas the two GT-B domains contain the active site and a hinge-latch system controlling catalytic activity^1,2^. Domains were assigned by comparison with the *At*SuSy crystal structure (PDB: 3S27)^1^. The cellular targeting domain (CTD) comprises residues 1-125, the ENOD40 peptide-binding domain (EPBD) comprises residues 155–274, and the GT-B fold region comprises residues 275–805. The CTD and EPBD are connected by a linker spanning residues 126-154. The hinge is centered around residues 525 and 752, with associated coils extending approximately across residues 521-532 and 750-756, respectively. The hinge connects the N- and C-terminal of the GT-B fold.

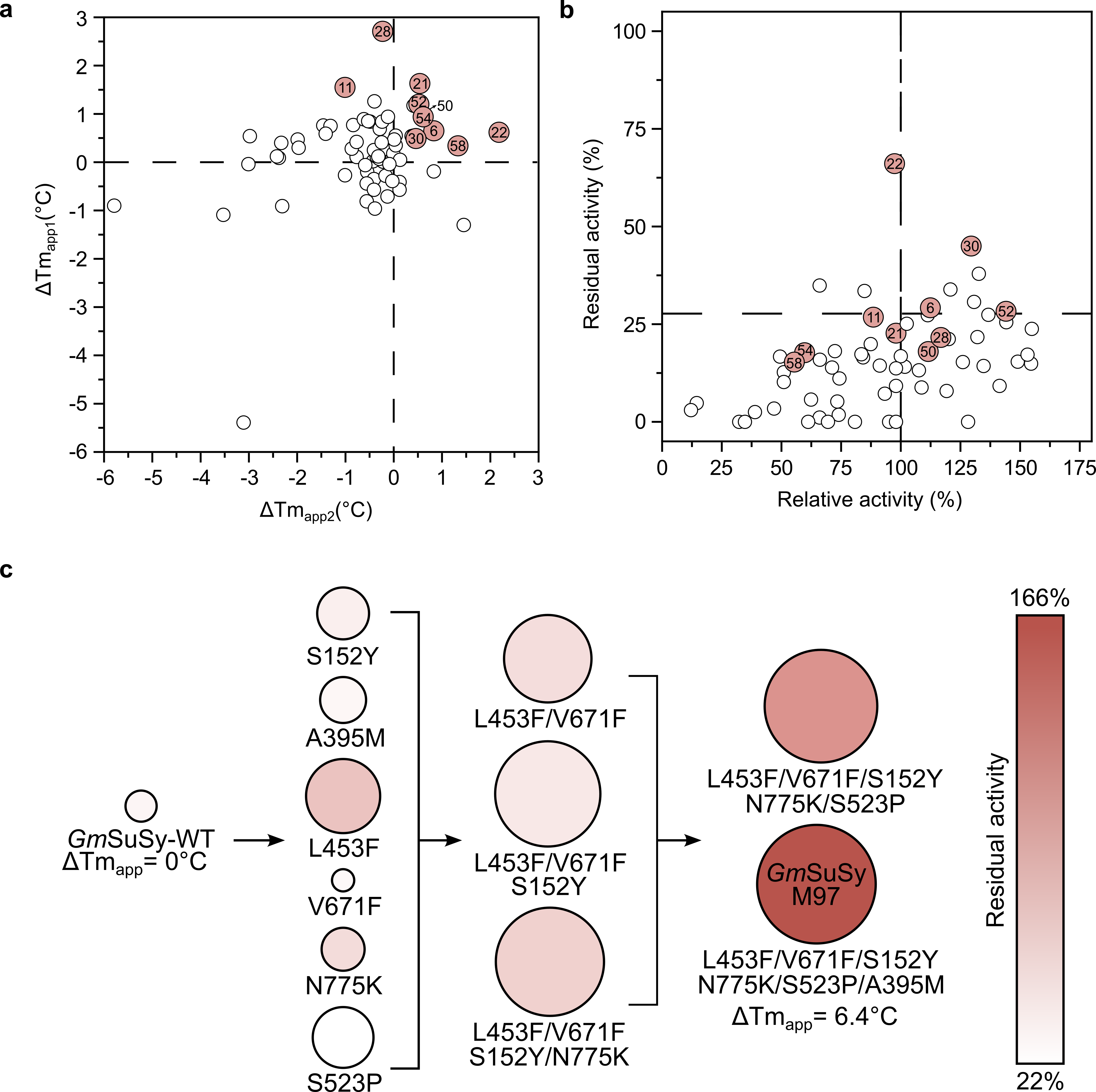

**Figure S2.** Single amino acid mutants were screened for **a**, thermostability and **b**, activity. Mutations selected for combinatorial analysis are shown in red and were selected because they improved in either thermostability or activity considering the change in melting temperature of the first (ΔTm_app1_) and second thermal transition (ΔTm_app2_), and both relative and residual activity. Numbers inside circles correspond to mutations as follows, 6,S152Y; 11, H534Y; 21, A395M; 22, L453F; 28, V671F; 30, N775K; 50, C660A; 52, N694H; 54, Q754K; and 58, S523P. **c**, Progressive combination of selected mutations from consensus-guided engineering. Melting temperatures from the second thermal transition (Tm_app_) are represented by the circle area while activity is represented by intensity in color as shown in the bar.

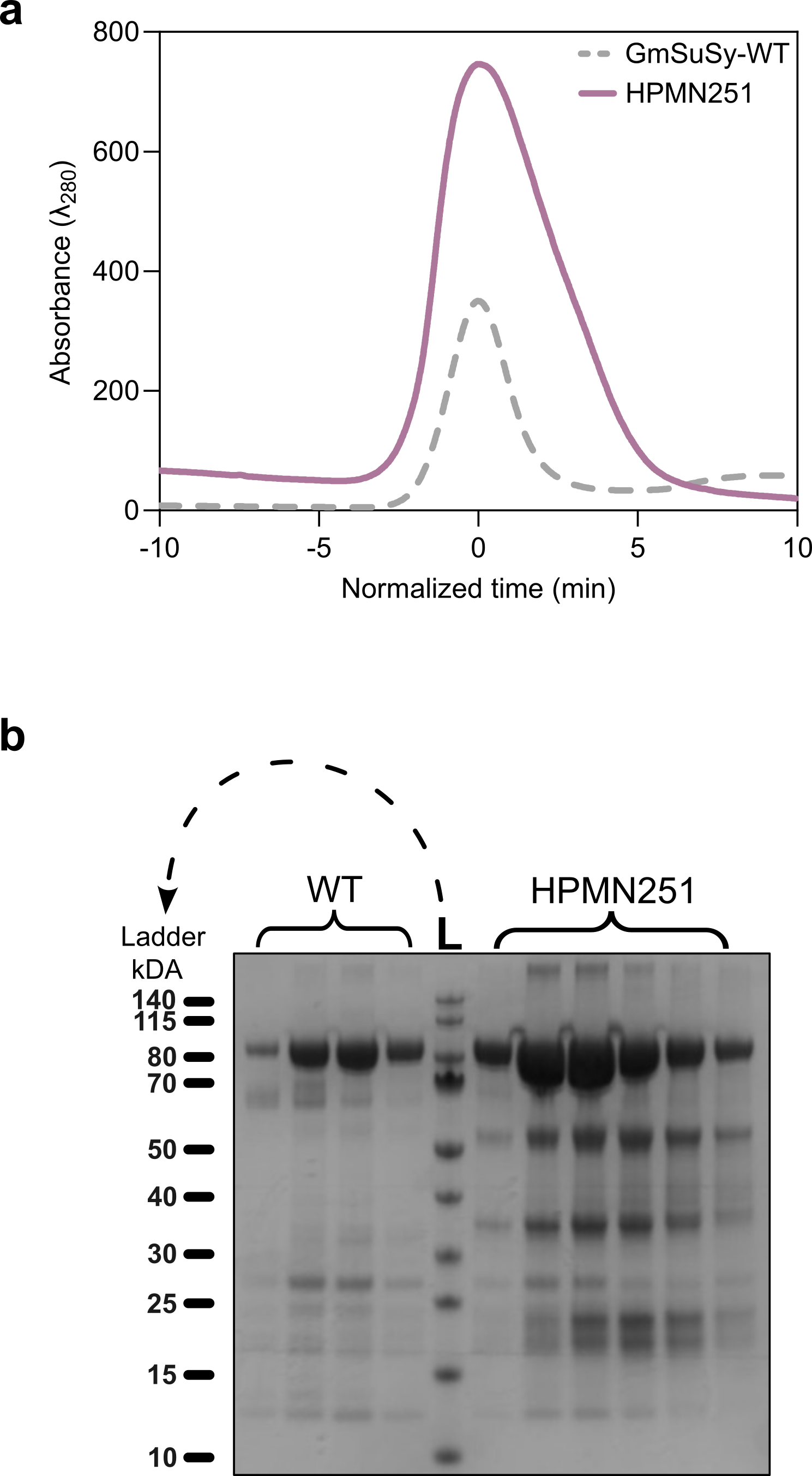

**Figure S3.** Increased soluble expression of HPMN251 compared with WT. **a,** Nickel-affinity chromatography elution profiles showing an increased elution peak for HPMN251, consistent with enhanced soluble expression. **b,** SDS-PAGE analysis of fractions collected after nickel-affinity chromatography from equivalent culture volumes, comparing soluble protein recovery for HPMN251 and WT.

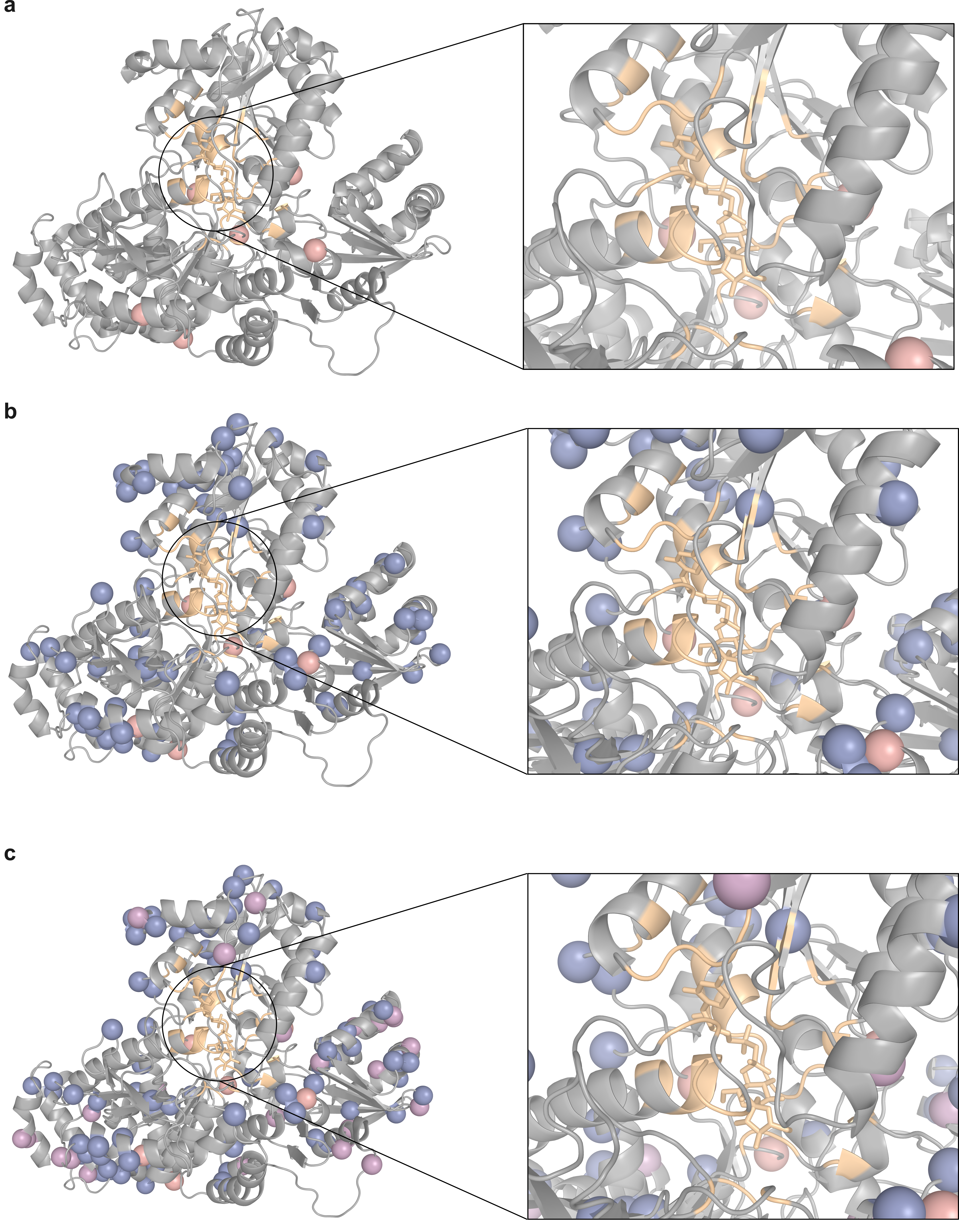
**Figure S4.** Cartoon representation of monomeric SuSy highlighting the locations of mutations as spheres and with the active site in a zoomed view. Sucrose, UDP, and residues within 5 Å are shown in yellow. Mutations are introduced progressively across variants with **a**, *Gm*SuSy-97 containing 6 mutations, shown as pink spheres; **b**, Anc165 containing 83 mutations, the original 6 mutations remain in pink, while the additional mutations are shown in blue; and **c**, HPMN251 containing 106 mutations; with remaining mutations in purple.

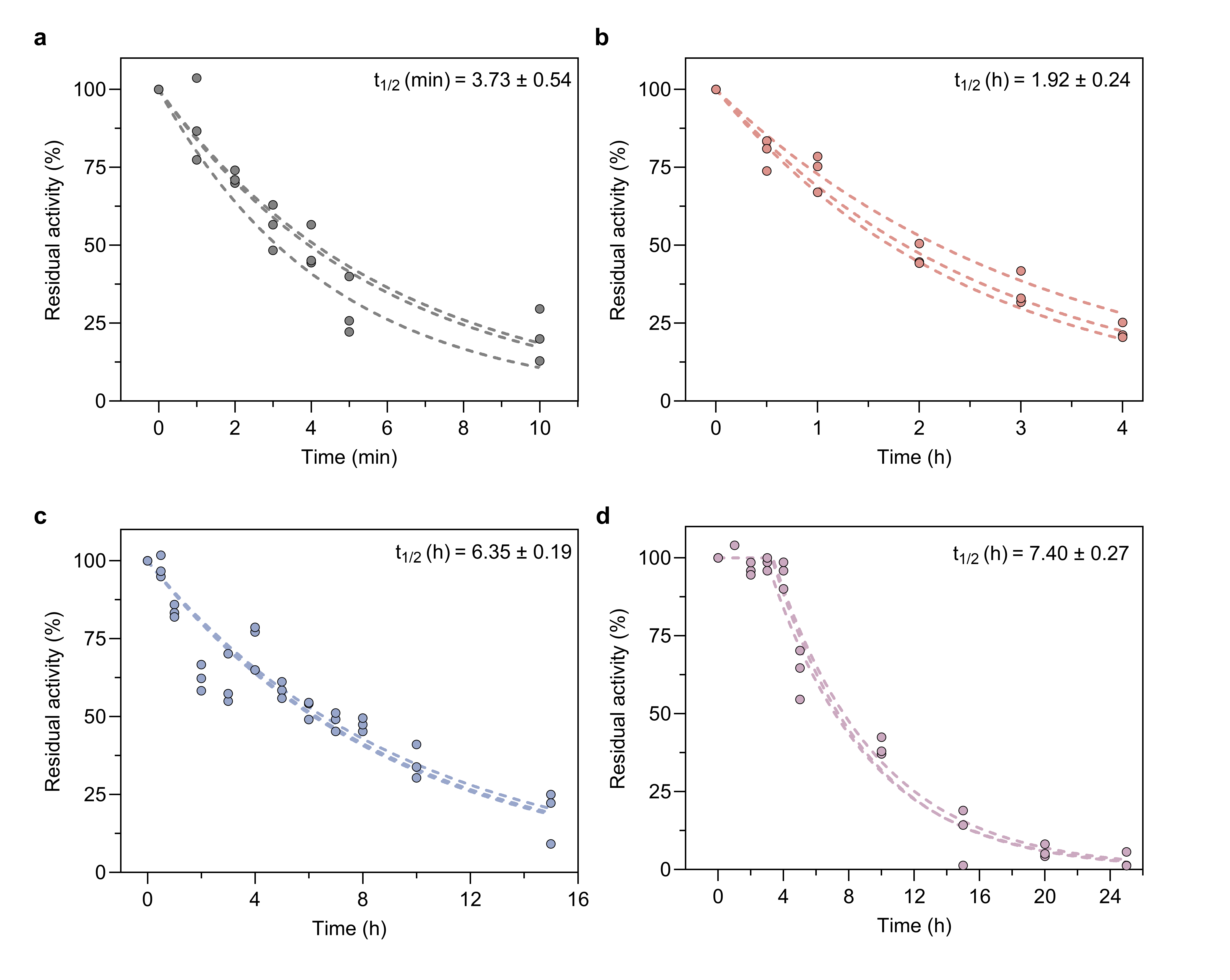
**Figure S5.** Kinetic stability profiles and predicted half-lives at 60 °C calculated based on a one-phase decay for **a**, WT; **b**, *Gm*SuSy-97; **c**, Anc165; and a plateau followed by one-phase decay for **d**, HPMN251*.* Circles correspond to the triplicates (*n*=3) at different time points, and dotted lines refer to the corresponding predicted model.

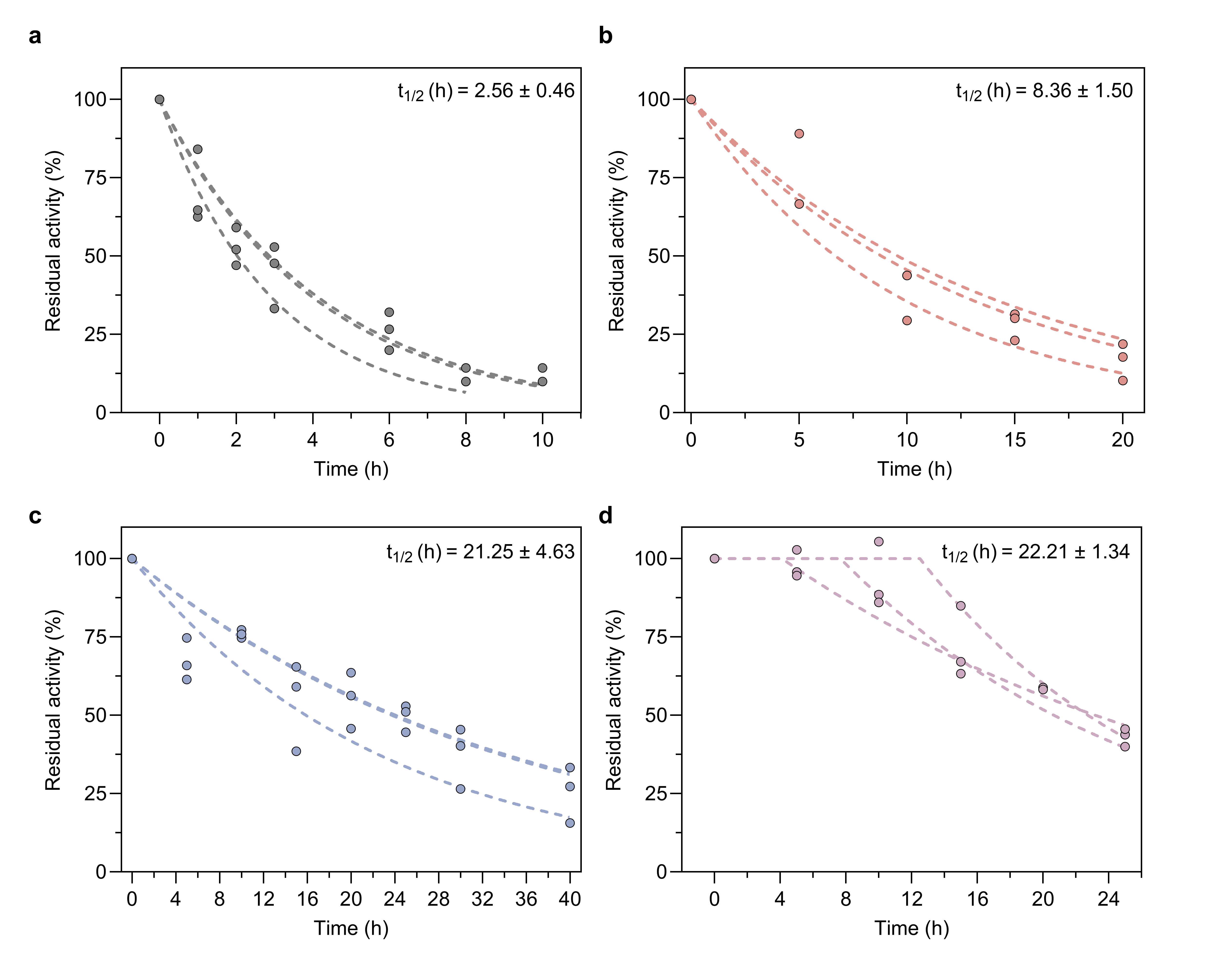
**Figure S6.** Kinetic stability profiles and predicted half-lives at 55 °C calculated based on a one-phase decay for **a**, WT; **b**, *Gm*SuSy-97; **c**, Anc165; and a plateau followed by one-phase decay for **d**, HPMN251*.* Circles represent individual replicate measurements (*n*=3) at different time points, and dotted lines refer to the corresponding predicted model.

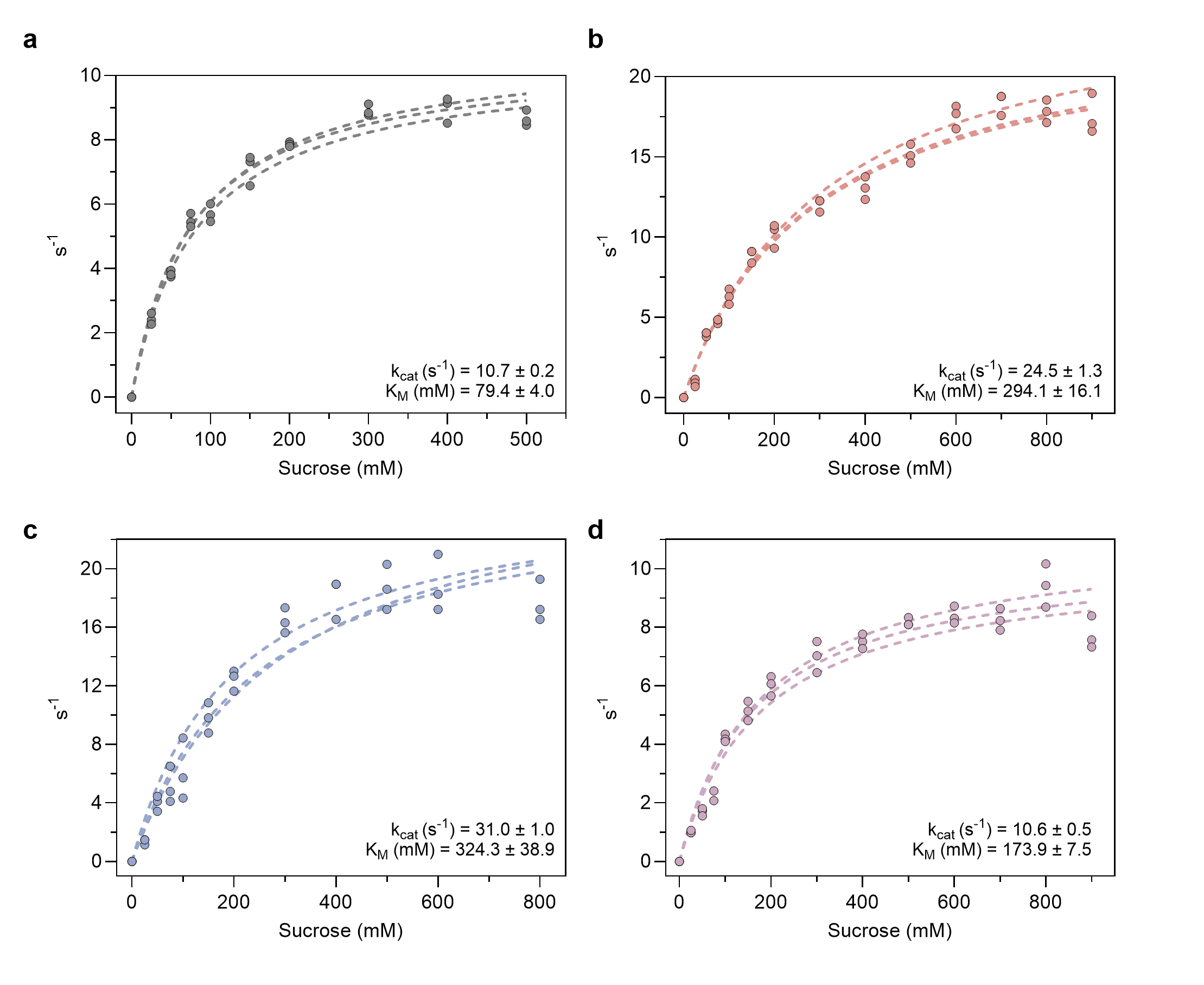
**Figure S7.** Michaelis-Menten models and predicted k_cat_ and K_M_ for engineered SuSy variants **a**, WT; **b**, *Gm*SuSy-97; **c**, Anc165; and **d**, HPMN251*.* Circles correspond to the triplicates (*n*=3) at different sucrose concentrations, and dotted lines refer to the corresponding predicted model.

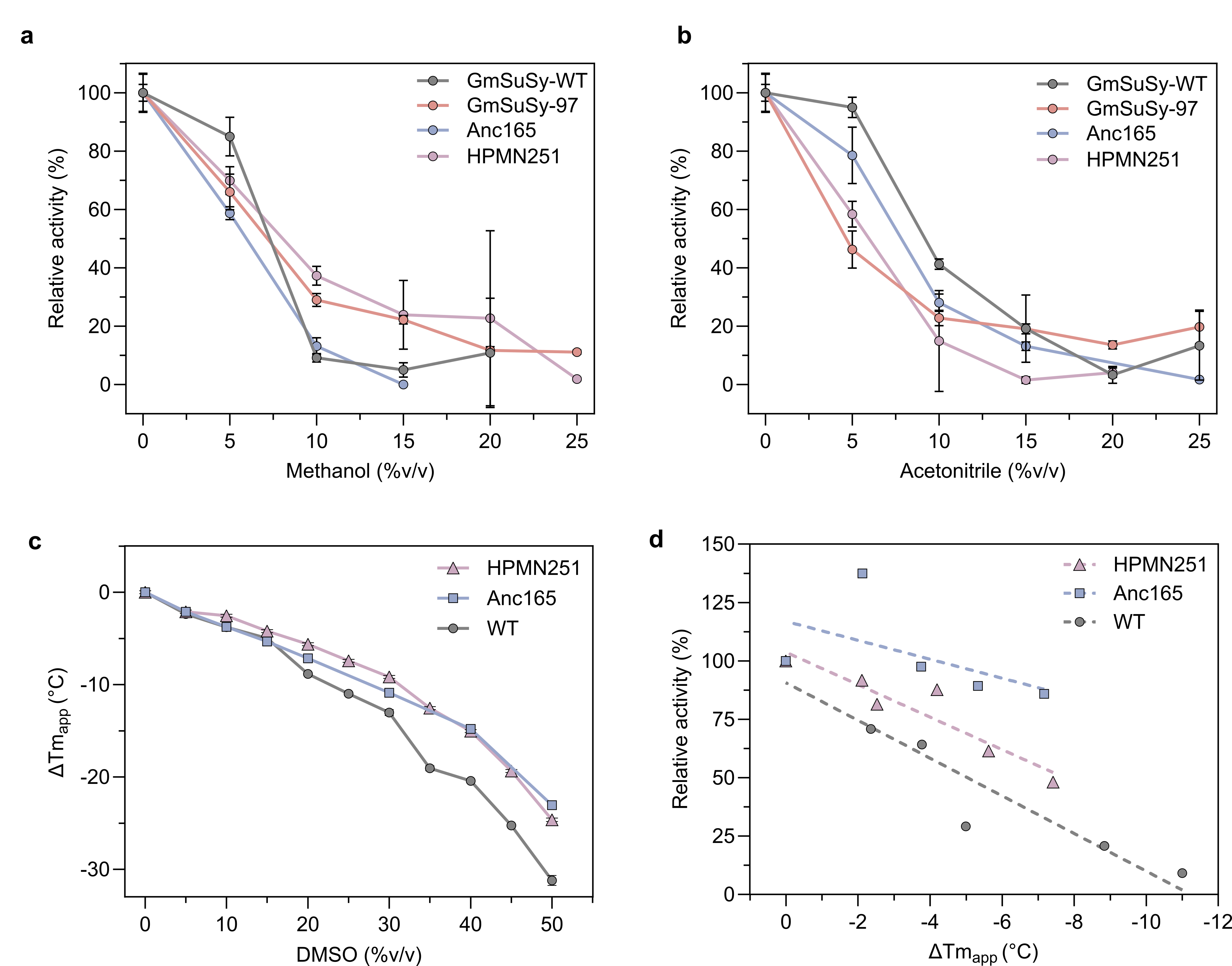

**Figure S8.** Solvent tolerance profiles of all variants to varying concentrations of **a**, methanol and **b**, acetonitrile. Error bars correspond to standard deviations from triplicates (*n*=3). **c,** Changes in apparent melting temperature (ΔTm_app_) at varying concentrations of DMSO, with error bars corresponding to standard deviations from duplicates (*n*=2) and including error propagation. **d**, Linear correlation between changes in Tm_app_ and relative activity at varying concentrations of DMSO. Dotted lines correspond to the predicted linear model.

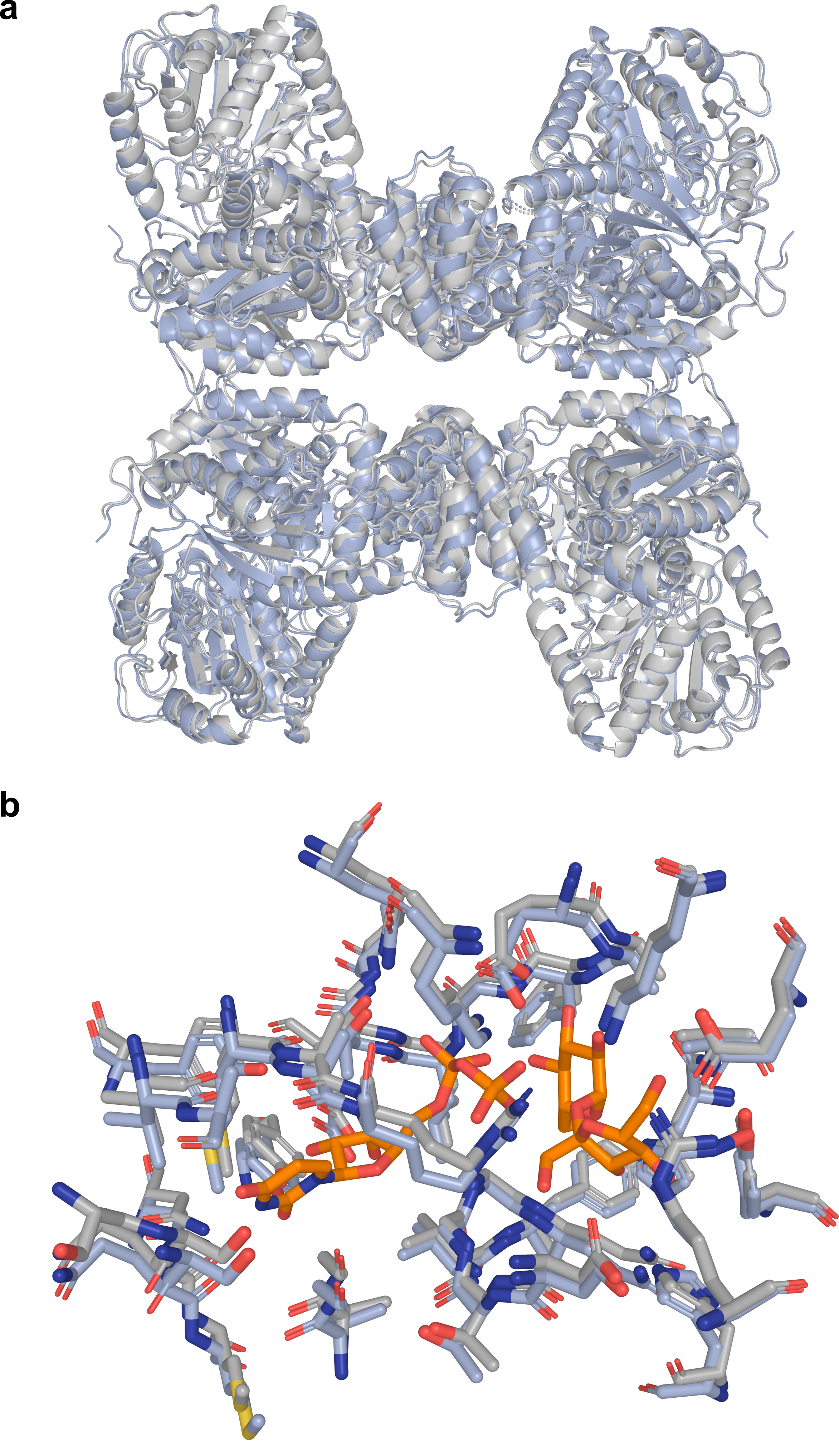

**Figure S9.** Structural superimposition of tetrameric WT and variants. **a**, Superimposition of the WT and Anc165 tetramers, with WT shown in gray and Anc165 shown in blue. The structures aligned with a C_α_ RMSD of 1.132 Å, with the main structural differences localized at the tetrameric interfaces. **b**, Anc165 mutations did not alter the active-site architecture, as shown by residues within 5 Å of UDP and sucrose, highlighted in orange. WT carbon atoms are shown in gray, and Anc165 carbon atoms are shown in blue. Superimposition of WT and *Gm*SuSy-97 revealed no relevant structural differences and is therefore not shown, as the two structures were essentially identical.

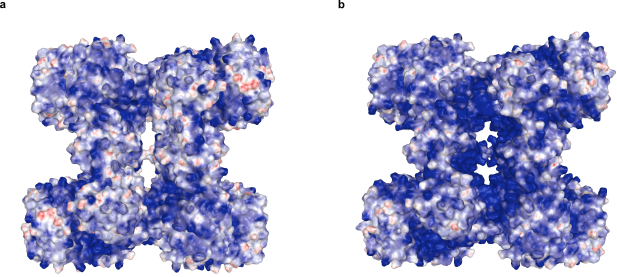

**Figure S10**. Electrostatic potential of the tetrameric SuSy surface at the optimal pH for **a**, WT and **b**, Anc165.

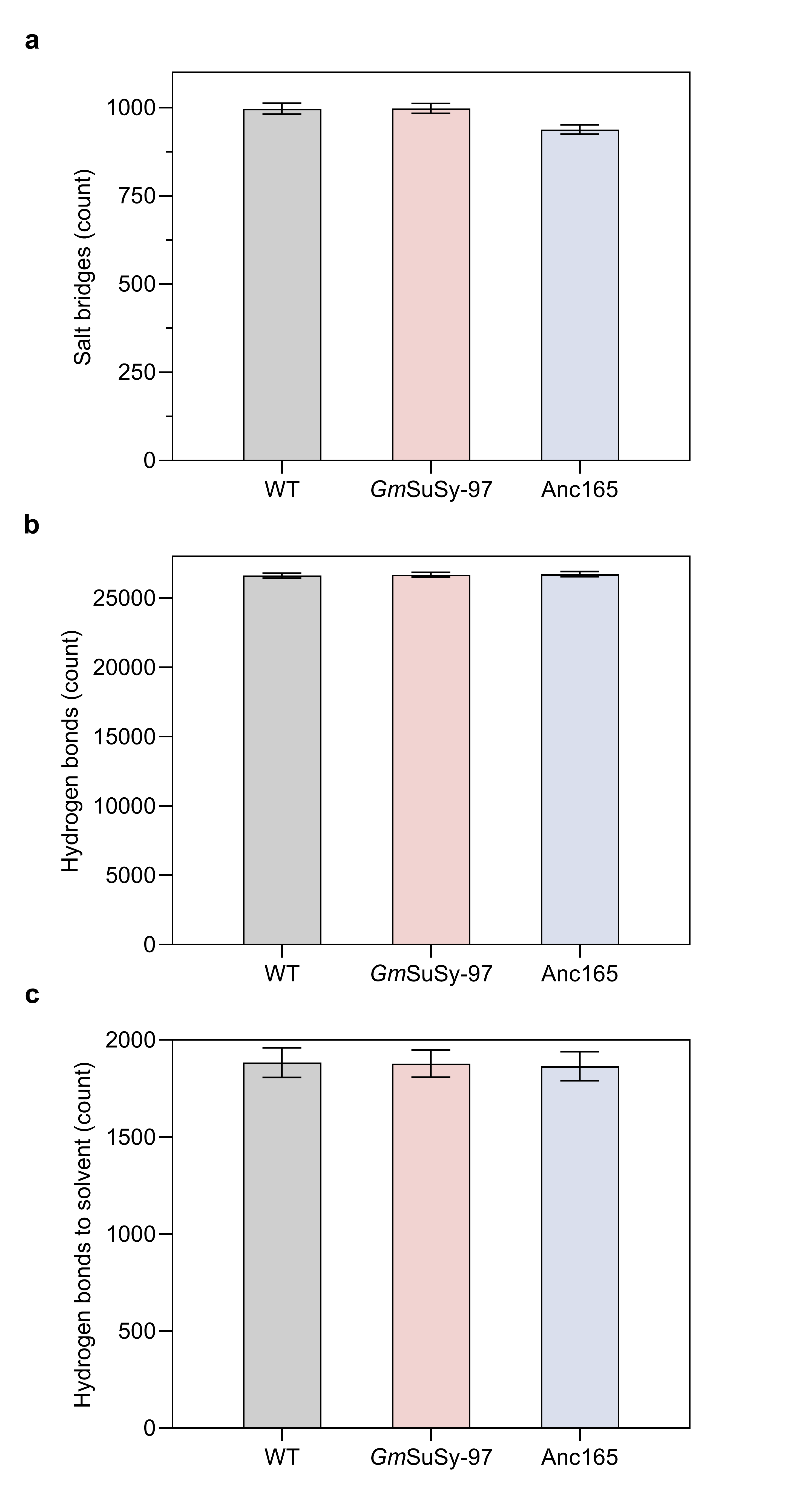

**Figure S11.** Interaction counts over molecular dynamics trajectory of **a**, salt bridges; **b**, hydrogen bonds within the protein; and **c**, hydrogen bonds between the protein and the solvent. Bar heights reflect means and error bars refer to standard deviations of the interaction counts (*n*=1800) from the 3.6 µs trajectories of classical molecular dynamics simulations.

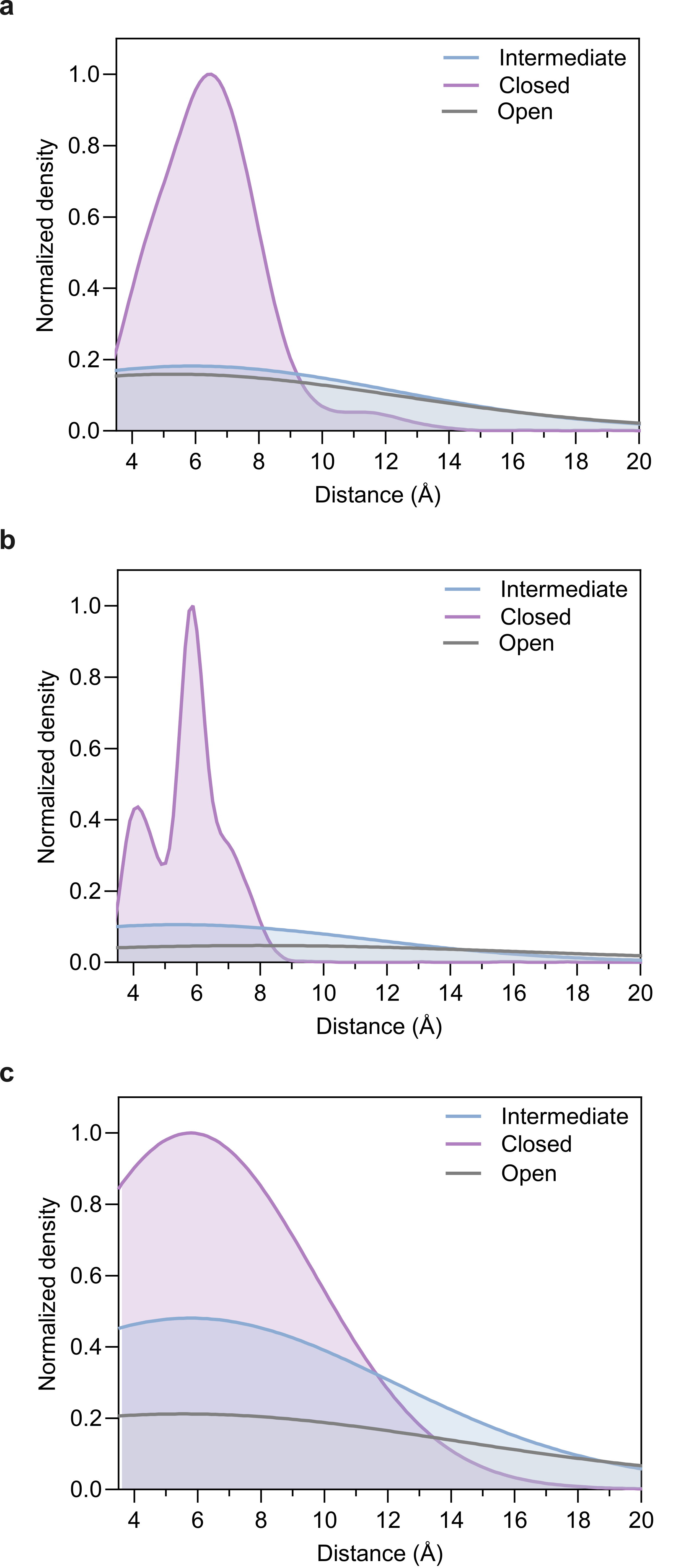

**Figure S12**. **a-c**, Distribution of catalytic ligand distances. Proton transfer from UDP to sucrose is assessed by measuring the distance between the β-phosphate center (Pβ) and the glycosidic oxygen (O). The distributions are shown for the closed, intermediate, and open states in **a**, WT; **b**, *Gm*SuSy-97; and **c**, Anc165. Only the closed cluster exhibited productive catalytic bonds.

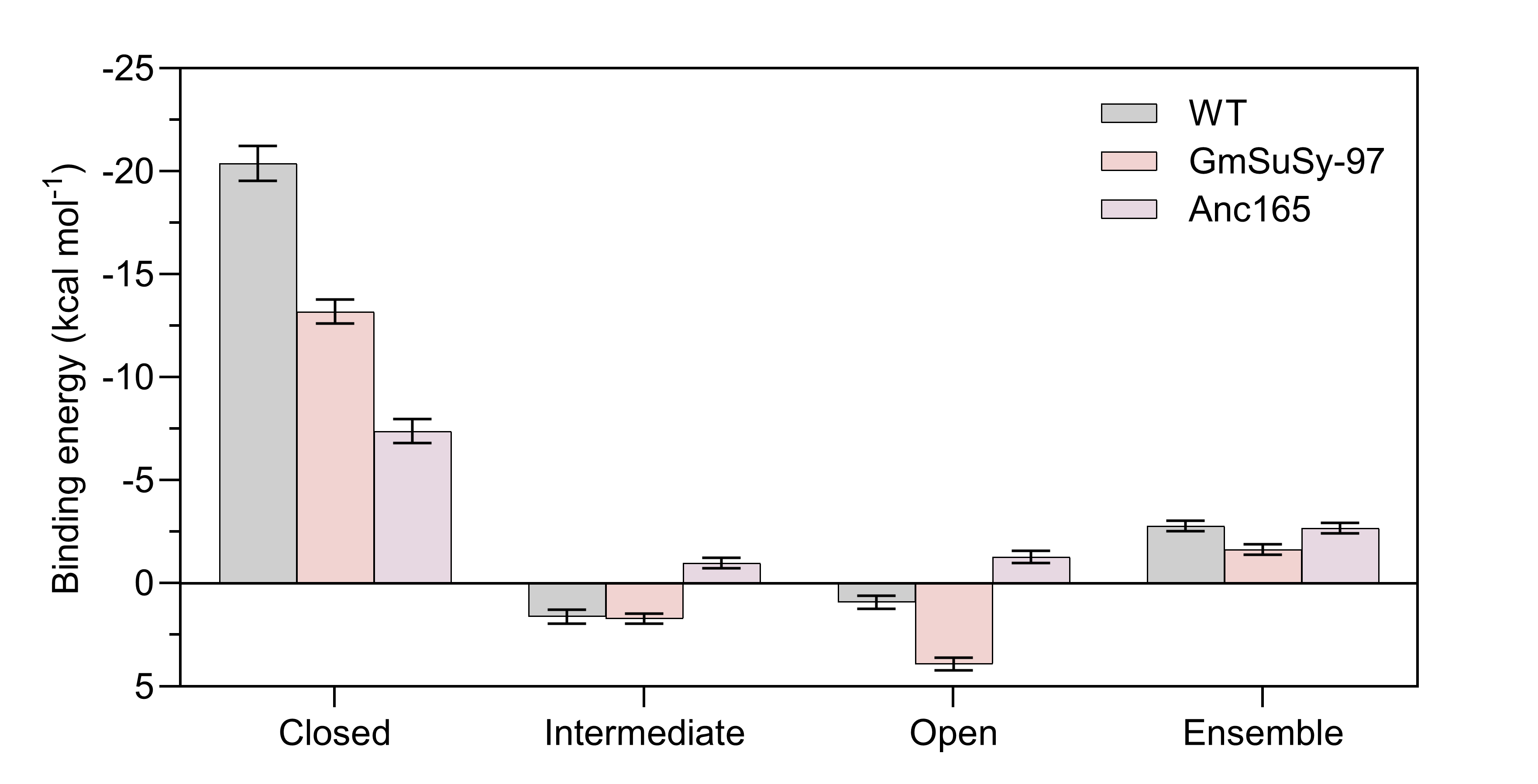
**Figure S13**. Ligand (sucrose and UDP) binding energies were calculated for the closed, intermediate, and open conformational states, as well as for the full ensemble, across the three variants: WT, *Gm*SuSy-97, and Anc165. Bar heights reflect means and error bars refer to standard deviations of the simulation steps (*n*=1800) from the 3.6 µs trajectories of classical molecular dynamics simulations. Only the closed cluster exhibited productive binding free energies. In contrast, substrate binding seems prohibitively unfavorable (> −1 kcal mol⁻¹) in the remaining two clusters (“intermediate” and “open”).

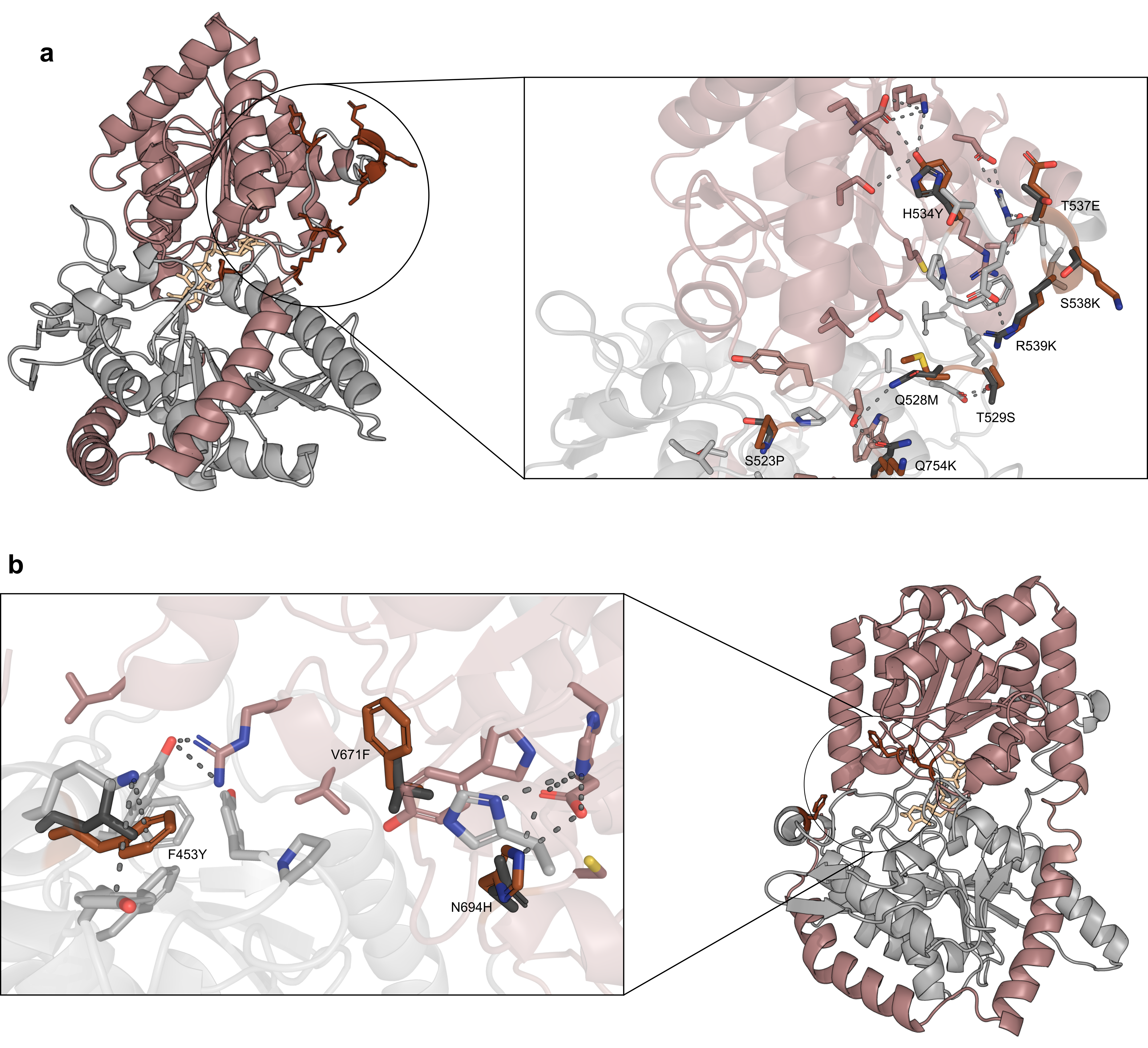
**Figure S14.** Mutations of Anc165 which are located in the **a**, hinge and **b**, latch of the GT-B fold. Possible modified molecular interactions are shown as dotted lines. WT residues are colored black while mutations in Anc165 are colored grey. The overall structure shows the GT-B fold with the C-terminal colored rose-beige and the N-terminal in grey.

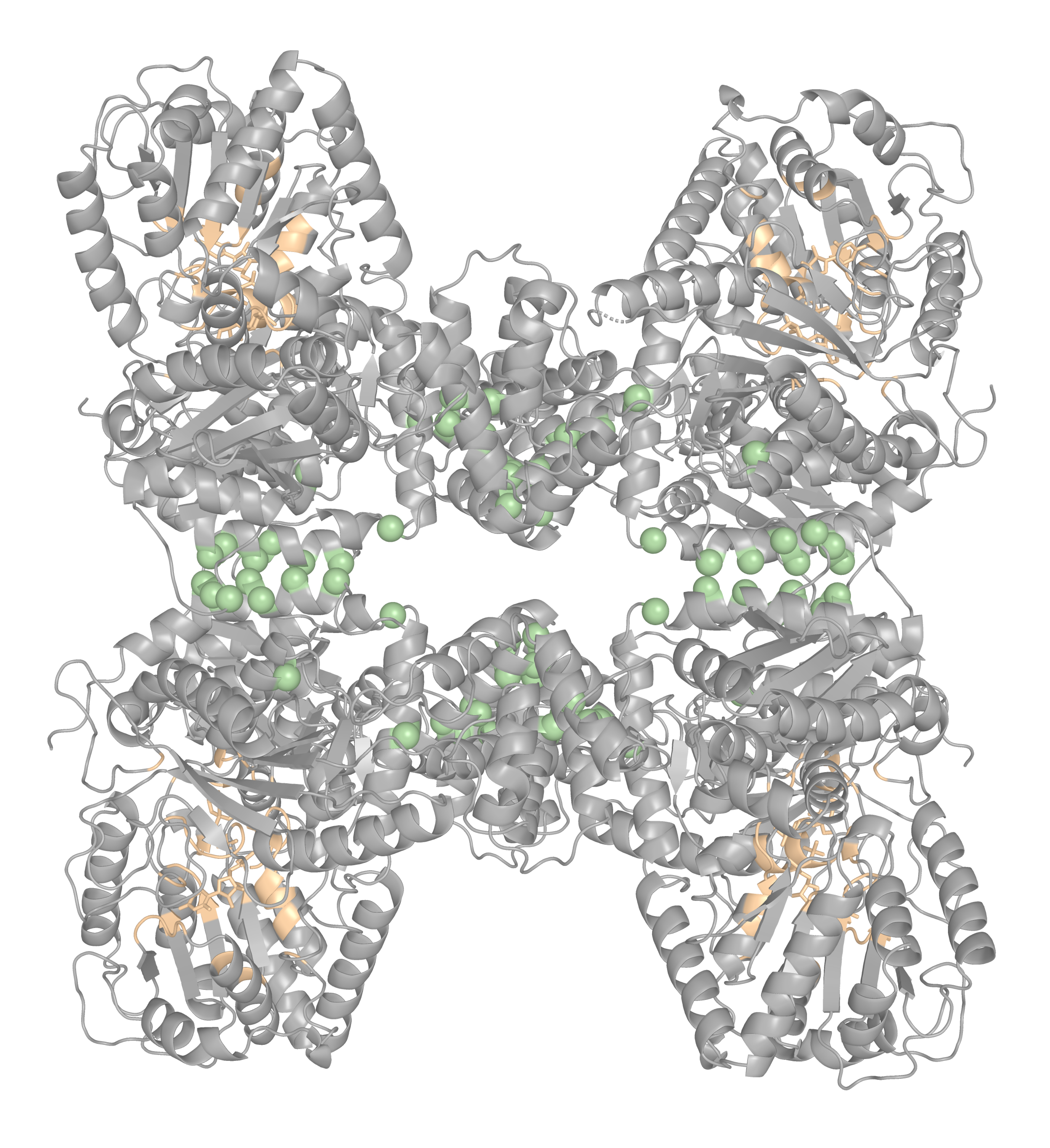
**Figure S15.** Cartoon representation of tetrameric WT with the locations of mutations evaluated in oligomeric interfaces, shown as green spheres. Sucrose, UDP, and residues within 5 Å are shown in yellow.

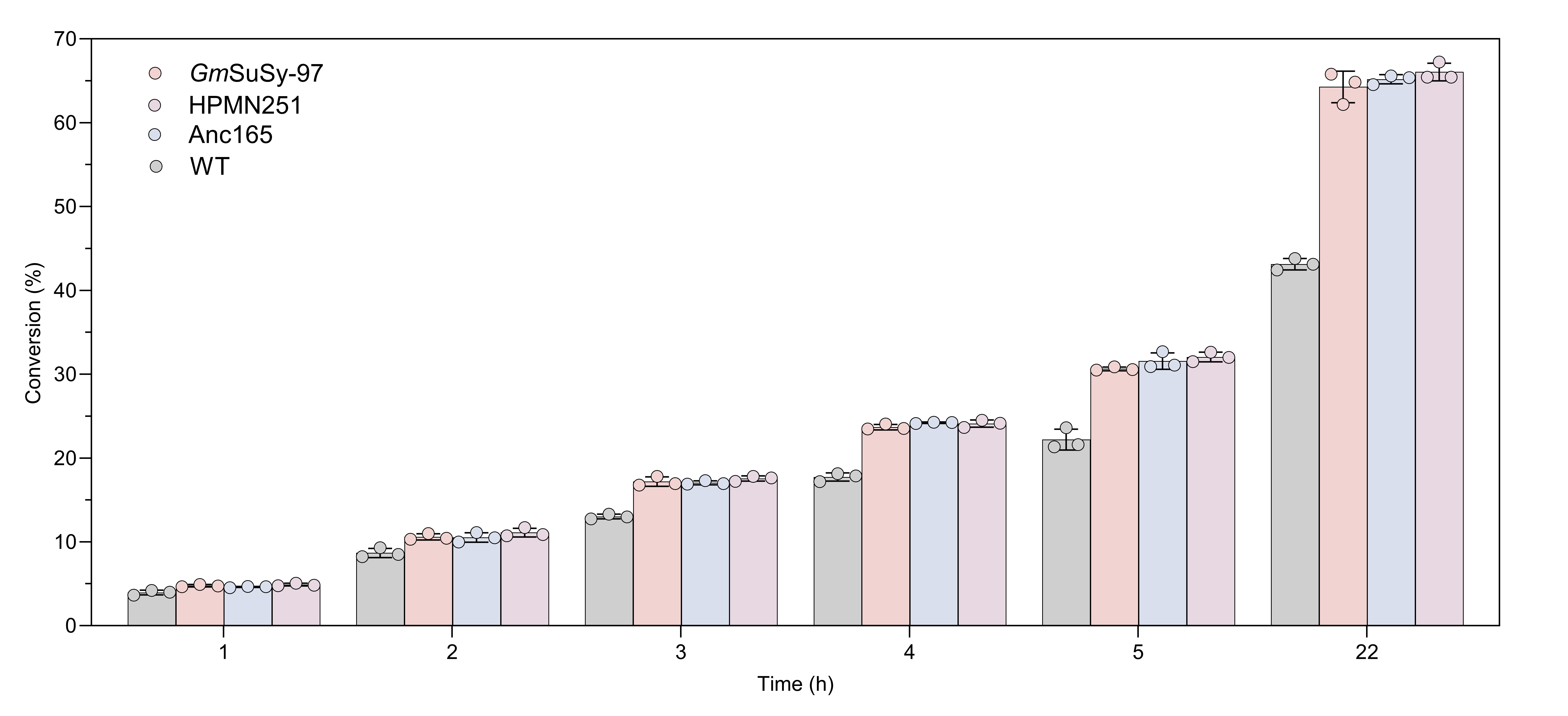
**Figure S16.** Screening of SuSy variants for MANT glycosylation over time and the corresponding conversion percentages. Circles represent individual data points (*n*=3), bar heights reflect means and the error bars their standard deviations.

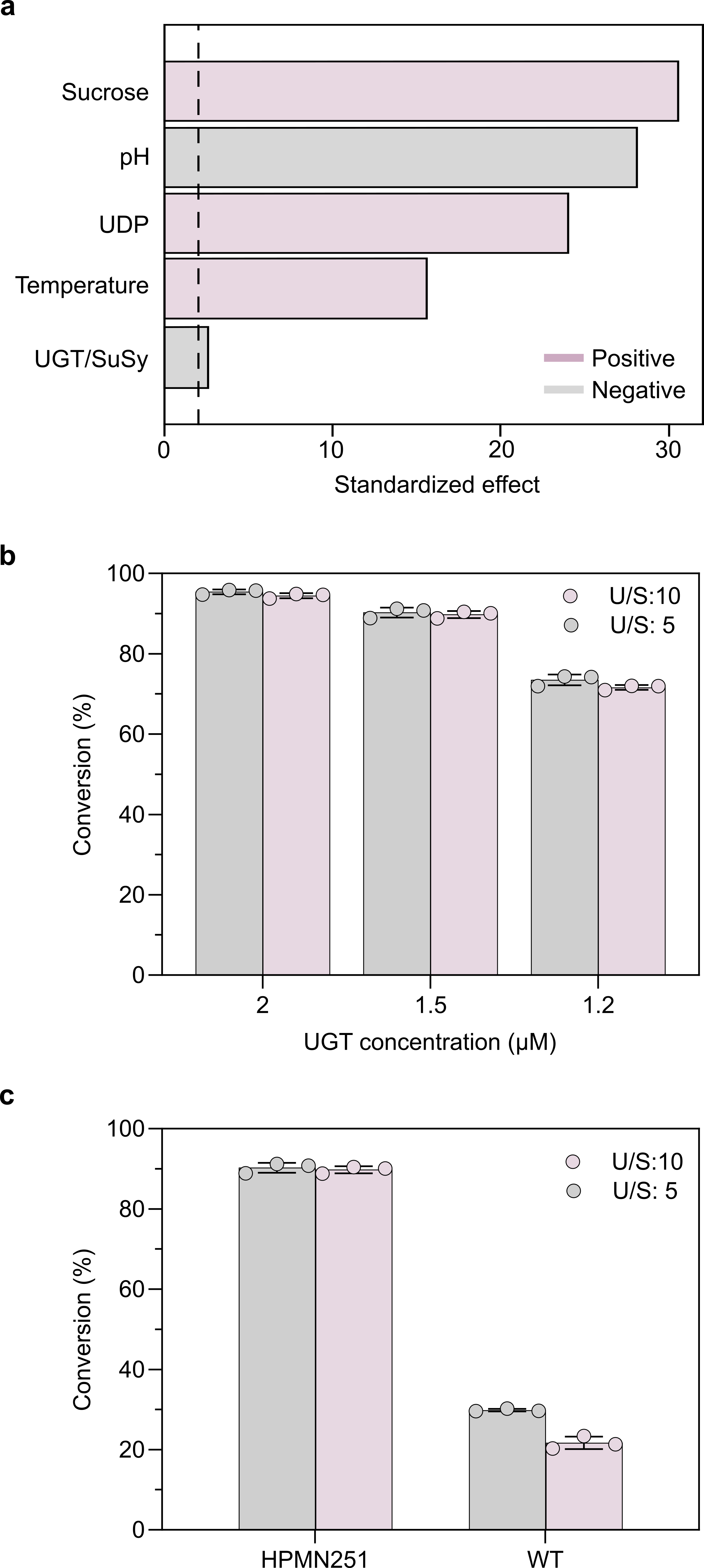

**Figure S17.** Optimization of MANT glycosylation cascade. **a**, Pareto chart illustrates the effects of reaction parameters on MANT conversion, where positive effects indicate that increasing the variable enhances conversion (purple) and negative effects indicate that decreasing the variable enhances conversion (gray bars). The dashed line denotes statistical significance at the 95% confidence level. Sucrose and UDP refer to concentrations of each compound. **b**, Evaluation of UGT concentration on glycosylation of 20 mM MANT with HPMN251 under optimized conditions and at different UGT/SuSy (U/S) ratios. **c**, Comparison of UGT/SuSy ratios between WT and HPMN251. Circles represent the triplicates (*n*=3) and, for panels a and b, bar heights reflect means and error bars indicate the corresponding standard deviations.

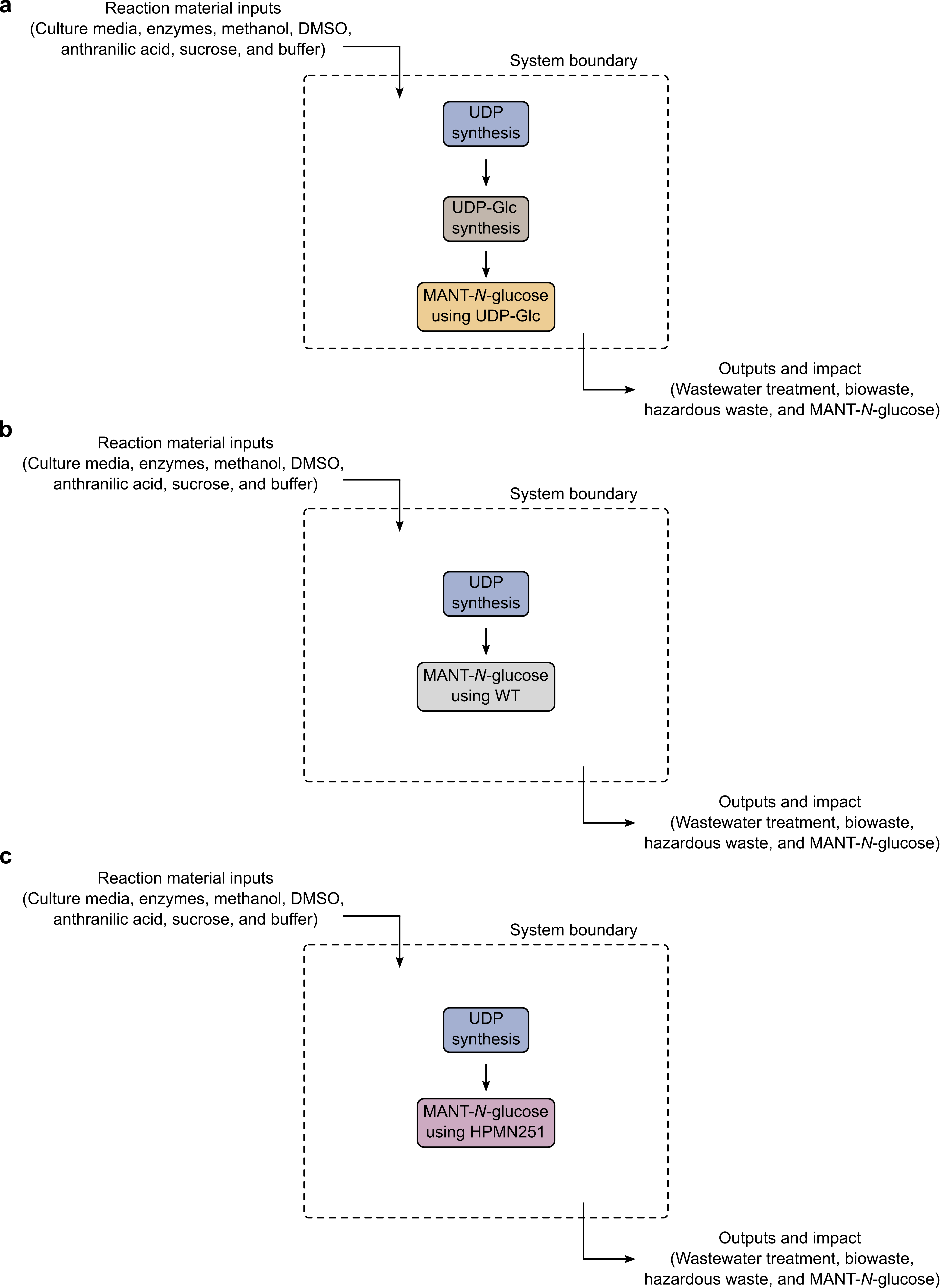
**Figure S18.** System boundaries defined for the preliminary life cycle assessment (LCA) of production of 1 kg MANT-*N*-glucose with three systems: **a**, UDP-Glc from literature^3^; **b**, using SuSy system with WT; and **c**, using SuSy system with HPMN251.

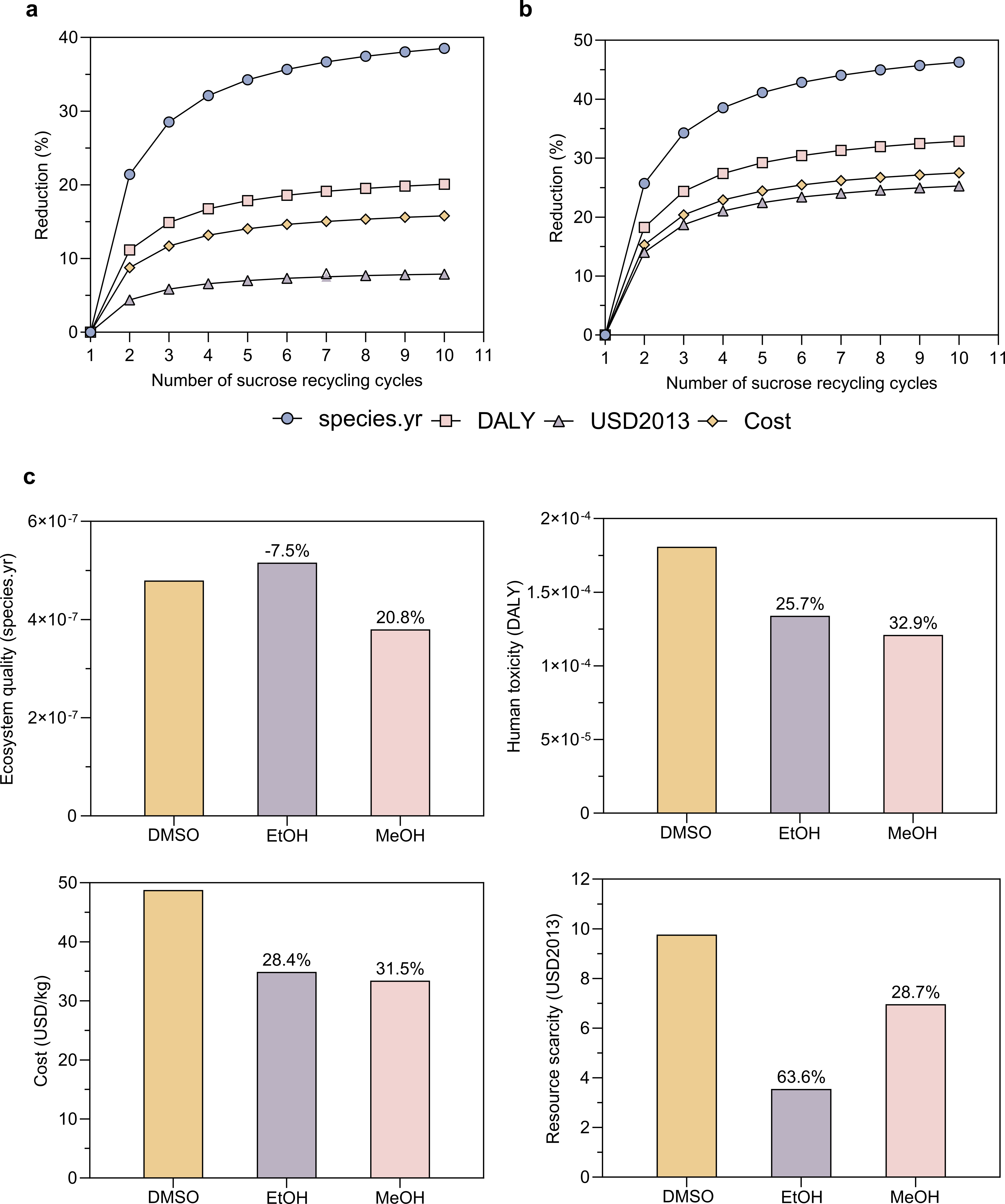

**Figure S19.** Solvent and substrate-recycling sensitivity analysis for UGT-SuSy cascade performance. **a,b,** Predicted relative reductions in endpoint categories and reaction costs as a function of sucrose recycling cycles for MANT and indoxyl glycosylation, respectively. **c**, Comparative sensitivity analysis of solvent substitution relative to DMSO. These projections identify solvent choice and sucrose recycling as major levers for improving process sustainability, although their quantitative benefits will require experimental validation of UGT-SuSy cascade performance in alternative solvents, including purification efficiency, product stability, and potential enzyme inhibition at the elevated product concentrations expected during repeated batch reuse.

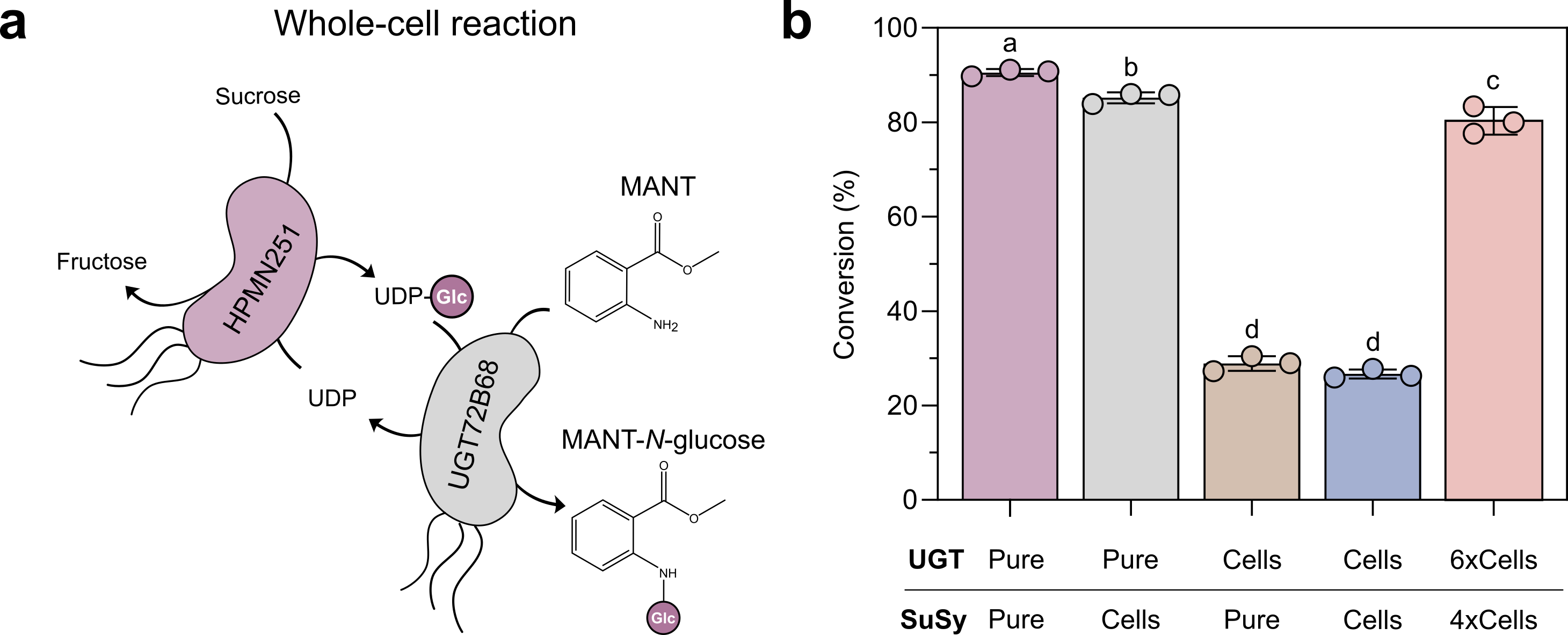

**Figure S20.** Evaluation of the whole cell reaction on MANT glycosylation. **a**, Combined biomass from two separate bacterial cultures overexpressing UGT and SuSy, respectively. **b**, MANT conversion using purified enzymes and biomass showed successful MANT glycosylation. UGT activity in biomass was limiting, whereas combining whole-cell HPMN251 with purified UGT resulted in only a 5.4% decrease in conversion compared to the pure enzyme system. Further increase of biomass led to approximately 80% conversion, demonstrating that SuSy can be effectively implemented as a whole-cell biocatalyst. Circles represent the triplicates (*n*=3), bar heights reflect means, and the error bars their subsequent standard deviations. Bars with different superscripts differ statistically significantly between them according to analysis of variance (ANOVA) with Tukey’s post hoc test (p < 0.05).

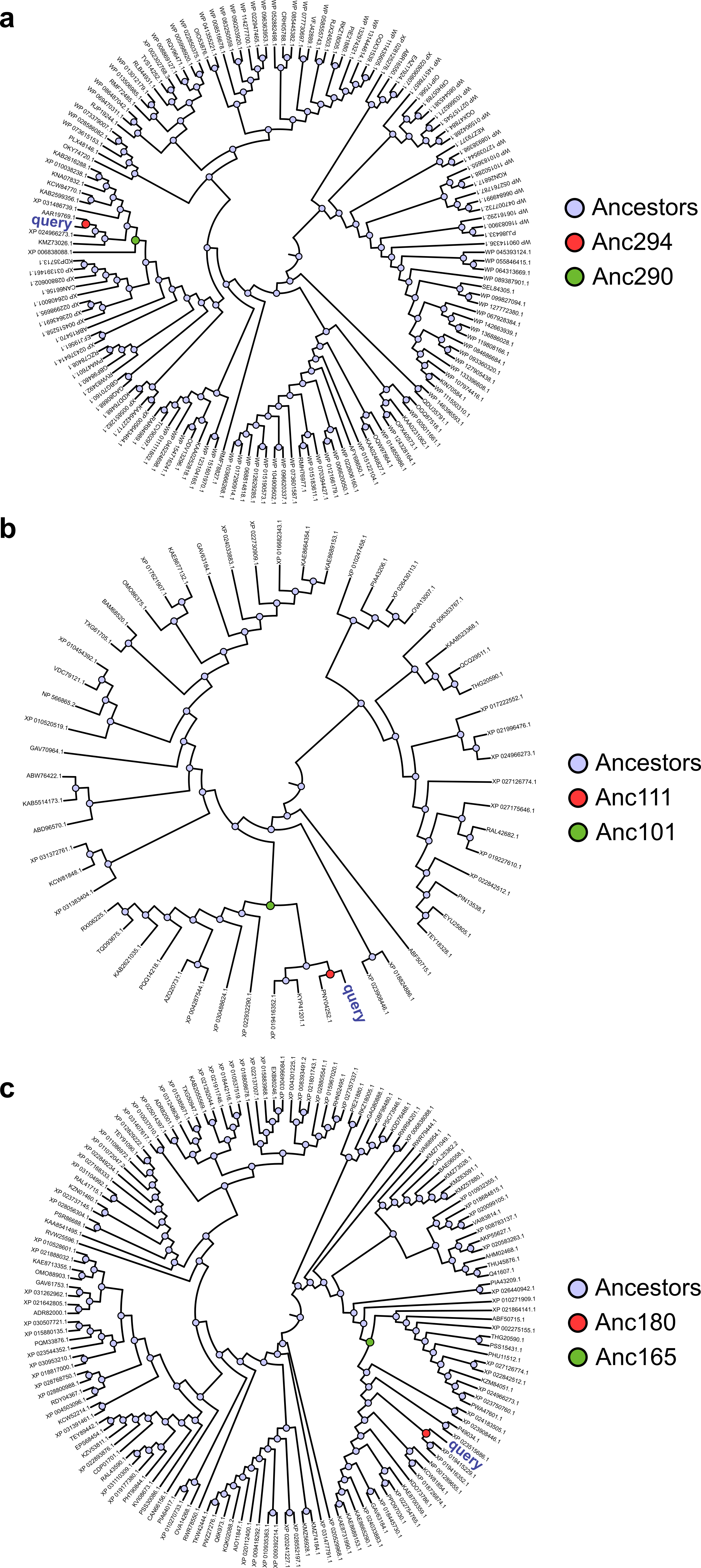

**Figure S21.** Phylogenetic trees and reconstructed ancestral sequences according to different levels of residue restriction. **a**, No residue restrictions with selected ancestors 294 (Anc294) and 290 (Anc290). **b**, Restriction of entire UDP binding site, selecting ancestors 111 (Anc111), and 101 (Anc101). **c**, Restriction of Q646, R649, R651, and N652, resulting in ancestors 180 (Anc180) and 165 (Anc165). Sequence name indicates GenBank protein accession number.

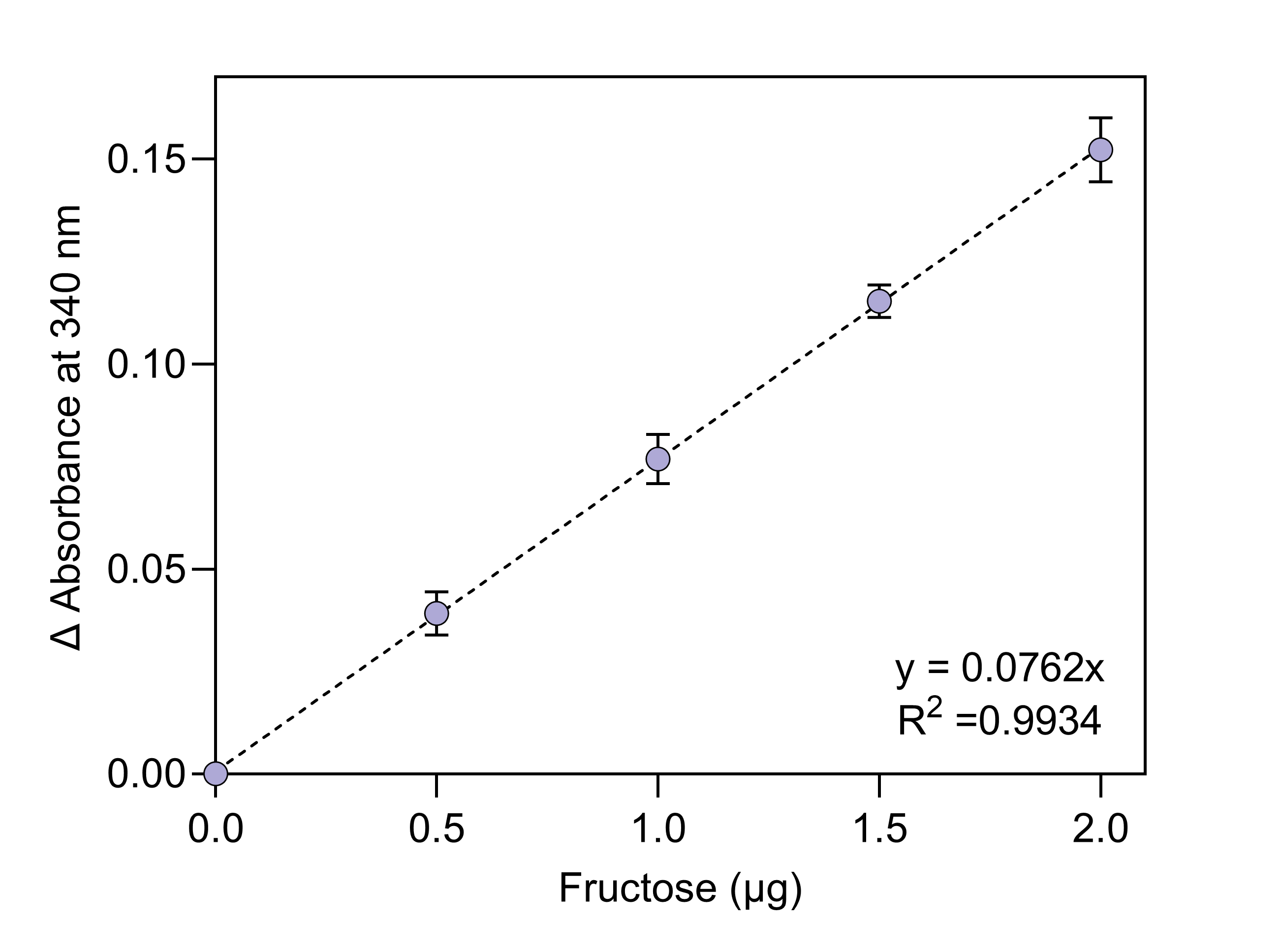
**Figure S22.** The fructose kit standard curve is defined with ΔAbsorbance (y) representing the change in absorbance before and after sample addition, and fructose amount in µg (x) as the independent variable. Circles represent the mean of triplicates (*n*=3) and error bars indicate the corresponding standard deviations.

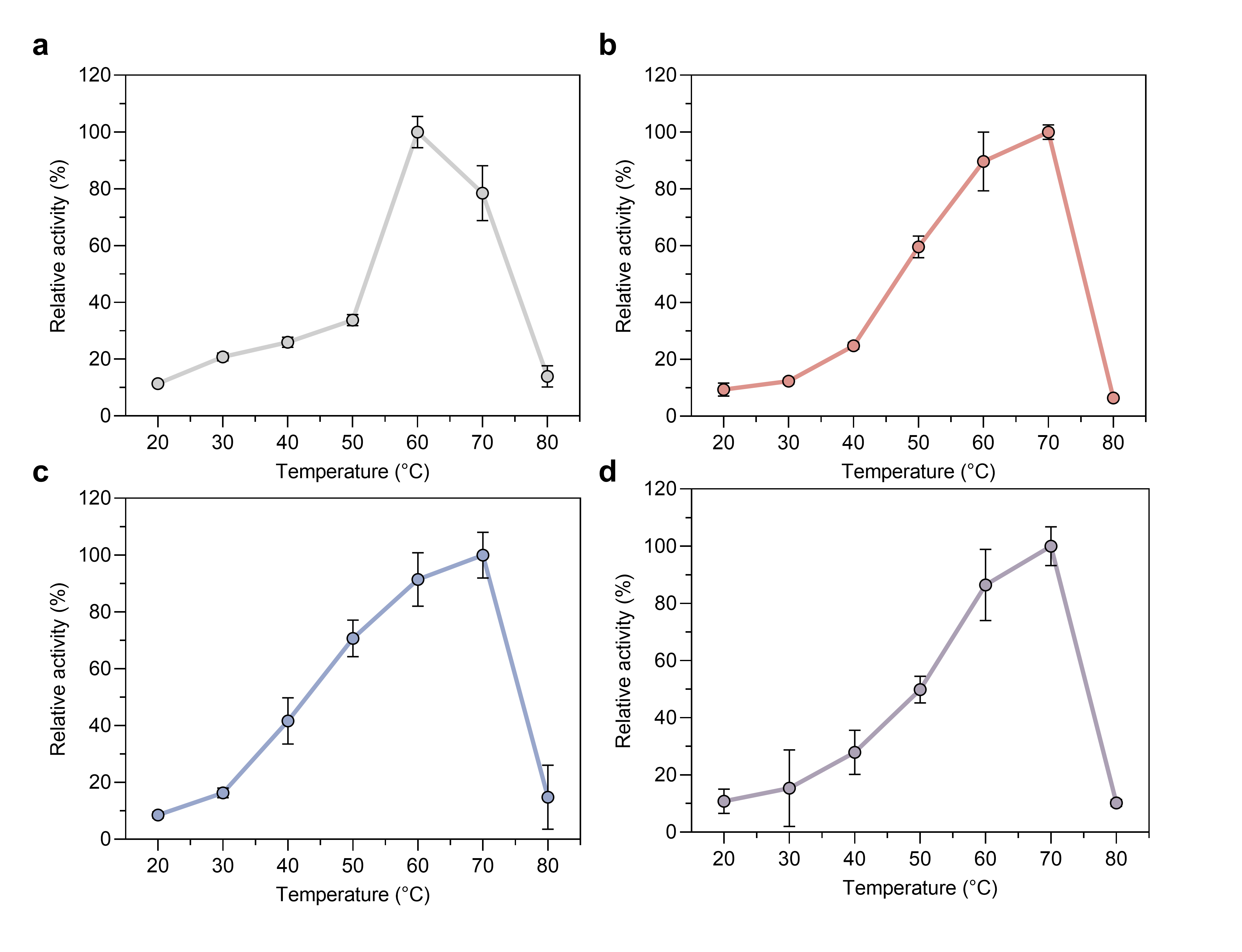
**Figure S23.** Effect of temperature on the catalytic activity of SuSy variants expressed as relative activity, with the highest activity for each variant set to 100%. **a**, WT; **b**, *Gm*SuSy-97; **c**, Anc165; **d**, HPMN251. Error bars correspond to standard deviations calculated from triplicates and include error propagation. Circles represent the mean of triplicates (*n*=3) and error bars indicate the corresponding standard deviations including error propagation.

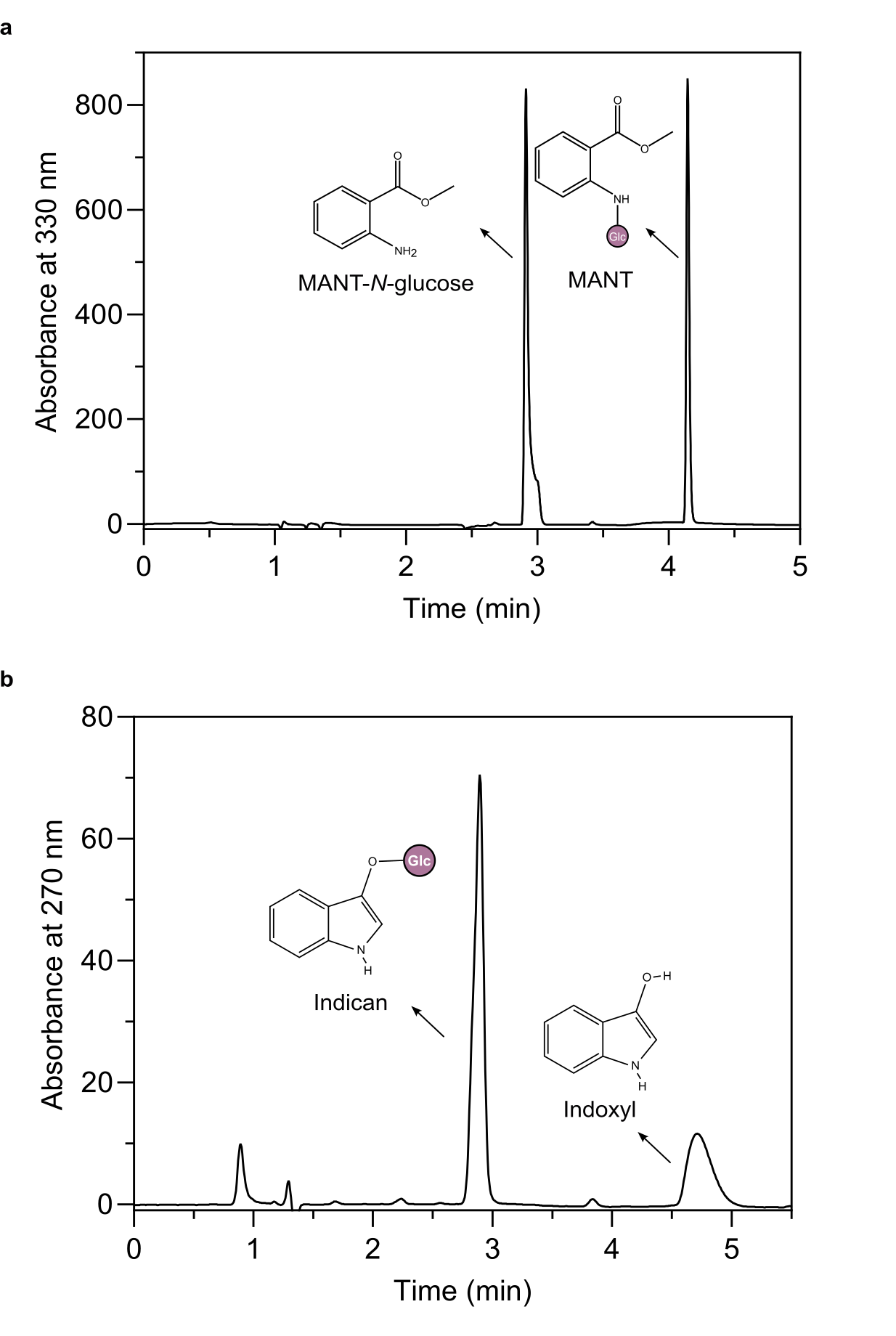
**Figure S24.** Example chromatograms for **a**, MANT (4.14 min) and MANT-*N*-glucose (2.91 min) detection. **b**, Indoxyl (4.70 min) and indican (2.89 min) detection.

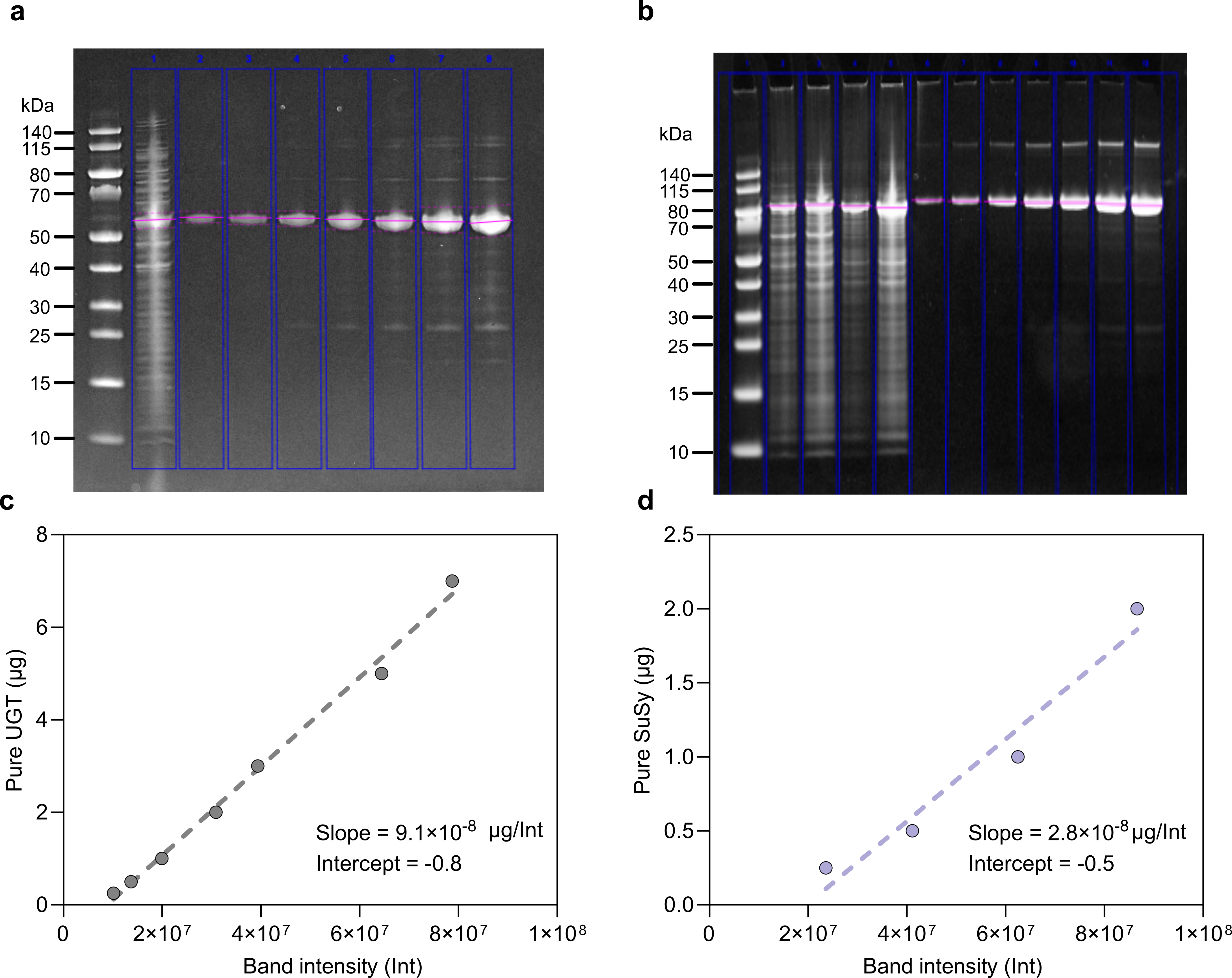

**Figure S25.** Quantification of enzyme in whole cell pellets from SDS-PAGE band intensity using a purified protein standard curve. **a**, SDS gel for UGT72B68 where lane 1 corresponds to 25 µg of wet cells and lanes 2 to 8 correspond to 0.25 µg to 7 µg of purified UGT72B68. **b**, SDS gel for SuSy where lanes 2-3 correspond to 37.5 µg and 75 µg of wet cell pellets of HPMN251 and lanes 6 to 12 correspond to 0.25 µg to 7 µg of purified *Gm*SuSy-WT. **c**, Linear correlation between band intensity and µg of pure protein for UGT72B68 and for **d**, HPMN251.

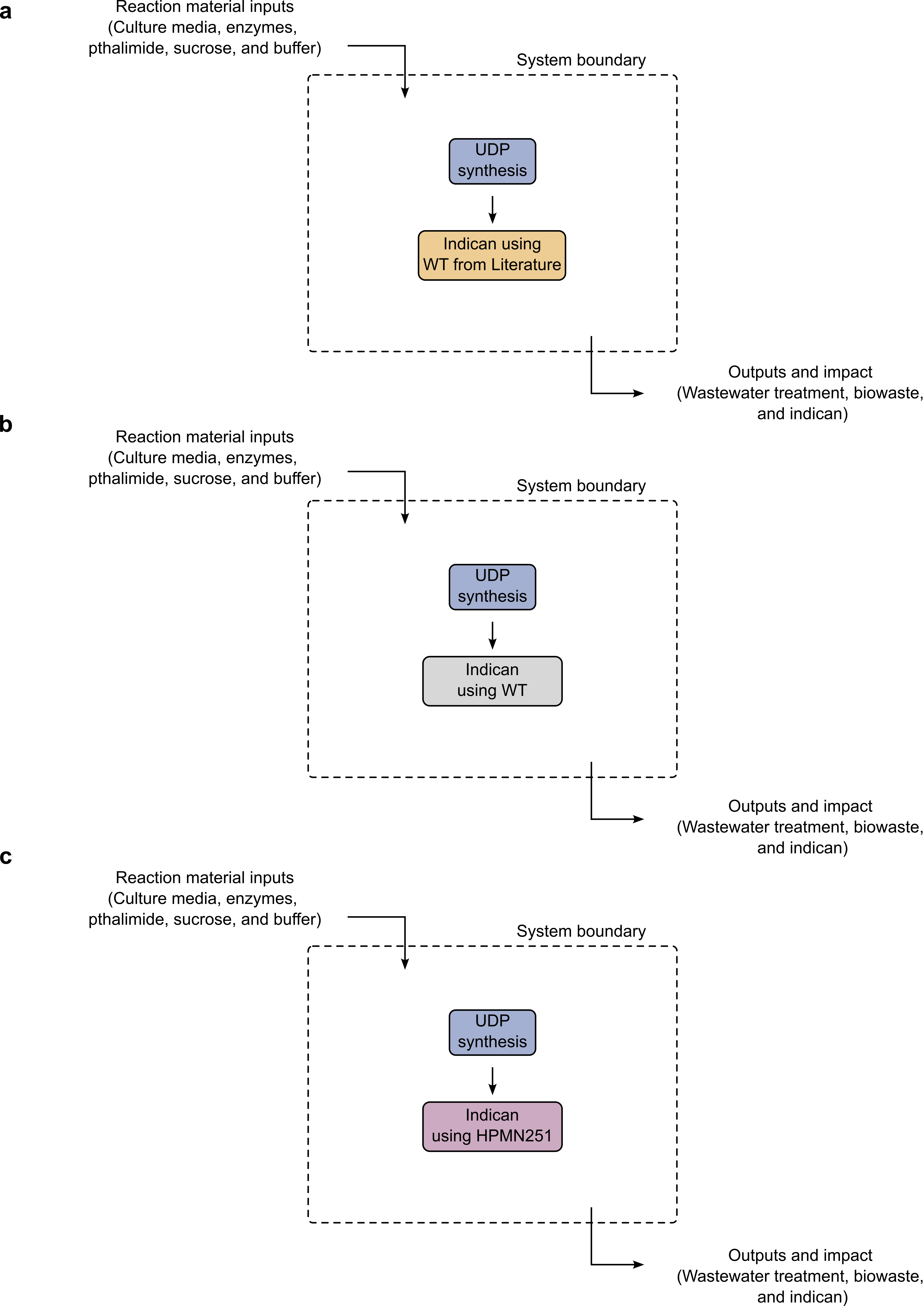

**Figure S26.** System boundaries defined for the preliminary life cycle assessment (LCA) of production of 1 kg indican with three systems: **a**, WT from literature^4^; **b**, WT from this work; and **c**, HPMN251.

**References**

1.	Zheng, Y., Anderson, S., Zhang, Y. & Garavito, R. M. The Structure of Sucrose Synthase-1 from Arabidopsis thaliana and Its Functional Implications *. *Journal of Biological Chemistry* **286**, 36108–36118 (2011).

2.	Wu, R. *et al.* The Crystal Structure of Nitrosomonas europaea Sucrose Synthase Reveals Critical Conformational Changes and Insights into Sucrose Metabolism in Prokaryotes. *J Bacteriol* **197**, 2734–2746 (2015).

3.	Gharabli, H. *et al.* Enzymatic Glycosylation of Anthranilates for Enhanced Functionality. 2025.05.29.656899 Preprint at https://doi.org/10.1101/2025.05.29.656899 (2025).

4.	Bidart, G. N. *et al.* Chemoenzymatic indican for light-driven denim dyeing. *Nat Commun* **15**, 1489 (2024).
