## Supplementary Tables for "Efficient and low-impact enzymatic glycosylation with robust sucrose synthase variants"

**Table S1.** Apparent melting temperatures and catalytic activity for single amino acid mutants from consensus-guided engineering.

| **Mutation** | **Thermal transition 1** | | | | **Thermal transition 2** | | | | **Activity** | | | |
| --- | --- | --- | --- | --- | --- | --- | --- | --- | --- | --- | --- | --- |
|  | **Tm_app_ (°C)** | **SD (°C)** | **ΔTm_app_ (°C)** | **SD (°C)** | **Tm_app2_ (°C)** | **SD (°C)** | **ΔTm_app_ (°C)** | **SD (°C)** | **Relative (%)** | **SD (%)** | **Residual (%)** | **SD (%)** |
| L453F | 46.28 | 0.56 | 0.62 | 0.57 | 63.76 | 0.03 | 2.18 | 0.03 | 97.4 | 12.8 | 66.1 | 9.6 |
| V658Y | 43.91 | 0.15 | -1.3 | 0.21 | 63.58 | 0.84 | 1.45 | 0.84 | 12.1 | 5.6 | 3 | 4.3 |
| S523P | 45.55 | 0.13 | 0.34 | 0.2 | 63.45 | 0.12 | 1.33 | 0.14 | 55.4 | 16 | 15.3 | 7.6 |
| S152Y | 45.78 | 0.08 | 0.65 | 0.13 | 63.55 | 0.06 | 0.83 | 0.07 | 112.4 | 39.5 | 29.2 | 3.6 |
| H319L | 45.02 | 0.52 | -0.19 | 0.54 | 62.95 | 1.43 | 0.83 | 1.43 | 14.5 | 5.9 | 4.8 | 6.9 |
| Q754K | 46.15 | 0.11 | 0.94 | 0.19 | 62.73 | 0.03 | 0.61 | 0.08 | 59.7 | 14 | 17.7 | 9.6 |
| A395M | 46.56 | 0.17 | 1.63 | 0.17 | 62.33 | 0.05 | 0.54 | 0.16 | 98 | 28.7 | 22.7 | 12.7 |
| C660A | 46.37 | 0.03 | 1.16 | 0.15 | 62.65 | 0.31 | 0.52 | 0.32 | 111.5 | 37.9 | 18 | 10 |
| N694H | 46.41 | 0.39 | 1.2 | 0.42 | 62.64 | 0.12 | 0.52 | 0.14 | 144.1 | 38.1 | 28.3 | 15 |
| V671F | 48.38 | 0.15 | 2.71 | 0.16 | 61.35 | 0.08 | -0.23 | 0.08 | 116.8 | 44.9 | 21.6 | 2.9 |
| H534Y | 46.69 | 0.09 | 1.55 | 0.14 | 61.72 | 0.06 | -1.01 | 0.07 | 88.5 | 9.7 | 26.8 | 14.1 |
| N775K | 46.15 | 0.17 | 0.49 | 0.18 | 62.04 | 0.06 | 0.46 | 0.06 | 129.5 | 53.6 | 45 | 8 |
| N694Y | 46.93 | 0.06 | 1.26 | 0.07 | 61.18 | 0.03 | -0.4 | 0.04 | 132.1 | 44.3 | 21.7 | 11.8 |
| T529S | 45.98 | 0.04 | 1.17 | 0.11 | 62.53 | 0.02 | 0.42 | 0.15 | 120.8 | 34.9 | 33.9 | 9.3 |
| Y390F | 45.48 | 0.18 | 0.55 | 0.19 | 61.83 | 0.12 | 0.04 | 0.19 | 132.7 | 39.1 | 37.9 | 20.9 |
| L86F | 45.97 | 0.07 | 0.84 | 0.13 | 62.23 | 0.01 | -0.49 | 0.03 | 66 | 11.2 | 34.9 | 18.8 |
| V410I | 46.02 | 0.05 | 0.89 | 0.12 | 62.11 | 0.08 | -0.62 | 0.08 | 84.8 | 16.3 | 33.5 | 18 |
| Q111P | 45.6 | 0.07 | 0.47 | 0.13 | 60.74 | 0.03 | -1.99 | 0.04 | 130.7 | 88.7 | 30.7 | 17.3 |
| V650L | 45.18 | 0.21 | 0.25 | 0.22 | 61.38 | 0.08 | -0.41 | 0.17 | 136.7 | 27.2 | 27.4 | 18.2 |
| T21S | 45.22 | 0.11 | 0.41 | 0.15 | 61.85 | 0.02 | -0.25 | 0.15 | 111.3 | 25.4 | 27.3 | 6 |
| V47I | 45.89 | 0.08 | 0.75 | 0.13 | 61.42 | 0.05 | -1.31 | 0.06 | 144.2 | 45.3 | 25.5 | 14.7 |
| Q758E | 44.23 | 0.1 | -0.57 | 0.15 | 61.7 | 0.03 | -0.41 | 0.15 | 102.5 | 22.9 | 25.1 | 14.4 |
| T222L | 44 | 0.31 | -0.81 | 0.33 | 61.54 | 0.15 | -0.56 | 0.17 | 154.9 | 75.1 | 23.8 | 12.8 |
| A104E | 45.85 | 0.1 | 0.18 | 0.1 | 61.57 | 0.06 | -0.01 | 0.07 | 97.3 | 31.4 | 22.2 | 11.8 |
| G348N | 44.82 | 0.2 | 0.01 | 0.23 | 61.88 | 0.05 | -0.22 | 0.08 | 120.1 | 56.7 | 21.2 | 11.4 |
| V218L | 45.41 | 0.06 | 0.28 | 0.12 | 61.86 | 0.03 | -0.87 | 0.04 | 87.4 | 31.8 | 19.9 | 11 |
| N23Y | 45.91 | 0.12 | 0.77 | 0.16 | 61.89 | 0.01 | -0.84 | 0.03 | 72.4 | 9.3 | 18.1 | 10 |
| Q528R | 45.98 | 0.06 | 0.85 | 0.12 | 62.2 | 0.03 | -0.53 | 0.05 | 83.5 | 10 | 17.3 | 9.4 |
| M317L | 44.48 | 0.12 | -0.44 | 0.13 | 61.23 | 0.17 | -0.56 | 0.23 | 153.1 | 31.1 | 17.2 | 10.5 |
| R456K | 45.4 | 0.12 | 0.47 | 0.14 | 61.8 | 0.11 | 0.01 | 0.19 | 100 | 20.9 | 16.8 | 9.2 |
| Q528M | 46.15 | 0.18 | 0.94 | 0.23 | 62.01 | 0.17 | -0.12 | 0.18 | 49.4 | 19.9 | 16.7 | 4.3 |
| H229R | 45.89 | 0.05 | 0.76 | 0.12 | 61.27 | 0.05 | -1.46 | 0.06 | 84.2 | 24.6 | 16.5 | 10.5 |
| E35V | 45.17 | 0.17 | -0.04 | 0.23 | 62.04 | 0.12 | -0.09 | 0.14 | 66.1 | 21.1 | 15.9 | 4.7 |
| A143S | 45.61 | 0.16 | 0.69 | 0.17 | 61.5 | 0.14 | -0.29 | 0.21 | 149 | 35.2 | 15.4 | 10.9 |
| N315M | 45.71 | 0.23 | 0.05 | 0.23 | 61.71 | 0.08 | 0.13 | 0.08 | 126 | 35.1 | 15.3 | 9 |
| L618R | 45.25 | 0.23 | -0.41 | 0.23 | 61.7 | 0.05 | 0.12 | 0.06 | 154.6 | 43.3 | 14.9 | 9 |
| G216L | 46.05 | 0.38 | 0.84 | 0.4 | 61.88 | 0.27 | -0.24 | 0.28 | 91.2 | 26.2 | 14.4 | 4.6 |
| V329T | 44.84 | 0.14 | 0.02 | 0.18 | 61.66 | 0.13 | -0.43 | 0.15 | 134.7 | 65.1 | 14.3 | 8.9 |
| G629L | 45.39 | 0.12 | -0.28 | 0.13 | 61.41 | 0.06 | -0.17 | 0.07 | 101.7 | 35.7 | 14.1 | 13 |
| V374I | 44.74 | 0.32 | -0.06 | 0.34 | 61.52 | 0.13 | -0.59 | 0.2 | 71.2 | 19.6 | 13.9 | 5.4 |
| R663K | 43.97 | 0.42 | -0.96 | 0.42 | 61.4 | 0.06 | -0.39 | 0.16 | 98 | 25.2 | 13.7 | 8.1 |
| G216R | 46.09 | 0.1 | 0.42 | 0.11 | 60.81 | 0.06 | -0.77 | 0.06 | 107.6 | 36.9 | 13.2 | 8.9 |
| S73Y | 45.46 | 0.08 | 0.54 | 0.1 | 62.16 | 0.15 | 0.37 | 0.21 | 51 | 13.4 | 12.7 | 8.4 |
| A66P | 45.25 | 0.09 | 0.11 | 0.14 | 61.95 | 0.03 | -0.77 | 0.04 | 74.3 | 28.5 | 11.1 | 7.8 |
| T537K | 44.5 | 0.05 | -0.71 | 0.16 | 61.99 | 0.16 | -0.13 | 0.18 | 51 | 13.4 | 10.2 | 4 |
| S250M | 45.27 | 0.05 | 0.35 | 0.07 | 61.83 | 0.02 | 0.04 | 0.15 | 98 | 19.3 | 9.2 | 6.5 |
| I755Y | 45.43 | 0.04 | 0.3 | 0.11 | 60.76 | 0.03 | -1.97 | 0.04 | 141.4 | 39.1 | 9.2 | 6.8 |
| F481Y | 45.31 | 0.07 | -0.35 | 0.09 | 61.16 | 0.11 | -0.41 | 0.12 | 108.7 | 31.9 | 8.8 | 6.2 |
| H206S | 44.93 | 0.15 | 0.12 | 0.19 | 61.78 | 0.23 | -0.32 | 0.28 | 119.2 | 38.3 | 7.9 | 8.1 |
| E180D | 45.11 | 0.23 | -0.2 | 0.24 | 61.78 | 0.03 | -0.55 | 0.03 | 93.3 | 12.3 | 7.2 | 9.3 |
| E18D | 44.64 | 0.12 | -0.57 | 0.19 | 62.24 | 0.1 | 0.12 | 0.13 | 62.4 | 15.3 | 5.7 | 8.8 |
| V132E | 44.42 | 0.28 | -0.9 | 0.28 | 56.54 | 0.17 | -5.79 | 0.17 | 73.4 | 11.2 | 5.2 | 4.6 |
| A622M | 45.47 | 0.21 | 0.55 | 0.22 | 62.2 | 0.04 | 0.41 | 0.16 | 46.9 | 9.5 | 3.4 | 4.5 |
| T734S | 44.82 | 0.13 | -0.39 | 0.2 | 62.09 | 0.19 | -0.03 | 0.21 | 38.9 | 10.7 | 2.5 | 4 |
| L796C | 45.54 | 0.03 | 0.4 | 0.11 | 60.39 | 0.04 | -2.33 | 0.05 | 74 | 23.3 | 1.8 | 5 |
| C563L | 39.92 | 0.13 | -5.39 | 0.14 | 59.22 | 0.16 | -3.11 | 0.16 | 66 | 6.9 | 1.1 | 3.8 |
| A395V | 45.03 | 0.21 | 0.11 | 0.22 | 61.44 | 0.12 | -0.35 | 0.19 | NR | NR | NR | NR |
| C563G | 44.54 | 0.33 | -0.27 | 0.35 | 61.08 | 0.05 | -1.01 | 0.08 | NR | NR | NR | NR |
| V374L | 45.52 | 0.06 | 0.59 | 0.08 | 60.38 | 0.15 | -1.41 | 0.21 | 98 | 20.3 | ND | ND |
| L687C | 45.67 | 0.12 | 0.54 | 0.16 | 59.74 | 0.03 | -2.98 | 0.04 | 80.8 | 14 | ND | ND |
| T490W | 45.04 | 0.24 | 0.12 | 0.25 | 59.37 | 0.25 | -2.42 | 0.29 | 34.7 | 7.4 | ND | ND |
| V132M | 45.23 | 0.06 | 0.09 | 0.12 | 60.35 | 0.03 | -2.38 | 0.04 | 95.1 | 34.4 | ND | ND |
| M624R | 44.99 | 0.19 | 0.07 | 0.2 | 61.49 | 0.07 | -0.3 | 0.17 | 61.2 | 13 | ND | ND |
| V514P | 45.62 | 0.03 | -0.04 | 0.05 | 58.57 | 0.02 | -3.01 | 0.03 | 128.2 | 37.4 | ND | ND |
| V132K | 44.41 | 0.34 | -0.91 | 0.34 | 60.02 | 1.68 | -2.31 | 1.68 | 69.5 | 10.8 | ND | ND |
| A465V | 44.58 | 0.13 | -1.09 | 0.14 | 58.05 | 0.17 | -3.53 | 0.17 | 32.3 | 4.5 | ND | ND |

SD refers to standard deviation of triplicates (*n*=3). Shaded rows indicate the selected variants for combinatorial mutagenesis. Seven variants with ΔTm_app_ of the second thermal transition > 0.5°C were selected, after excluding V658Y and H319L which exhibited loss of activity. Two additional variants were chosen based on the ΔTm_app_ of the first thermal transition > 1.5°C. N775K was also included due to its high residual activity relative to WT.

**Table S2.** Apparent melting temperatures for combined mutants from consensus-guided engineering.

| **Mutation** | **Thermal transition 1** | | | | **Thermal transition 2** | | | |
| --- | --- | --- | --- | --- | --- | --- | --- | --- |
|  | **Tm_app_ (°C)** | **SD (°C)** | **ΔTm_app_ (°C)** | **SD (°C)** | **Tm_app_ (°C)** | **SD (°C)** | **ΔTm_app_ (°C)** | **SD (°C)** |
| C660A;N694H | 47.34 | 0.32 | 1.76 | 0.37 | 62.88 | 0.10 | 0.29 | 0.14 |
| Q754K;C660A | 46.75 | 0.46 | 1.18 | 0.49 | 62.74 | 0.10 | 0.15 | 0.14 |
| C660A;S523P | 46.35 | 0.87 | 0.77 | 0.89 | 63.47 | 0.08 | 0.88 | 0.13 |
| N694H;Q754K | 46.69 | 0.06 | 1.11 | 0.19 | 63.05 | 0.28 | 0.46 | 0.30 |
| N694H;S523P | 46.55 | 0.01 | 0.97 | 0.18 | 63.74 | 0.24 | 1.15 | 0.26 |
| Q754K;S523P | 47.01 | 0.36 | 1.43 | 0.40 | 63.84 | 0.16 | 1.25 | 0.19 |
| H534Y;S523P | 47.63 | 0.44 | 2.05 | 0.48 | 63.69 | 0.02 | 1.10 | 0.10 |
| A395M;S523P | 47.81 | 1.70 | 2.24 | 1.71 | 64.19 | 0.08 | 1.60 | 0.13 |
| S152Y;N775K | 46.54 | 0.56 | 0.96 | 0.59 | 64.00 | 0.09 | 1.41 | 0.14 |
| L453F;V671F | 48.92 | 0.35 | 3.14 | 0.36 | 65.07 | 0.10 | 3.27 | 0.10 |
| S152Y;L453F | 45.83 | 0.28 | 0.05 | 0.29 | 65.11 | 0.10 | 3.32 | 0.11 |
| L453F;N775K | ND | ND | ND | ND | 64.25 | 0.04 | 2.46 | 0.04 |
| S152Y; V671F | 49.31 | 0.53 | 3.54 | 0.53 | 63.96 | 0.07 | 2.16 | 0.07 |
| V671F;N775K | 48.99 | 0.13 | 3.21 | 0.14 | 63.75 | 0.04 | 1.96 | 0.04 |
| N694H;Q754K;S523P | 48.98 | 1.02 | 3.07 | 1.04 | 63.62 | 0.06 | 1.29 | 0.08 |
| L453F;V671F;N775K | 48.66 | 0.67 | 3.85 | 0.68 | 66.09 | 0.15 | 4.44 | 0.28 |
| S152Y;L453F;N775K | 47.62 | 1.45 | 2.81 | 1.45 | 65.25 | 0.10 | 3.60 | 0.26 |
| S152Y;L453F;V671F | 49.52 | 0.16 | 4.71 | 0.20 | 66.66 | 0.09 | 5.02 | 0.25 |
| A395M;N694H;Q754K;S523P | ND | ND | ND | ND | 64.77 | 0.05 | 2.21 | 0.14 |
| A395M;L453F;V671F;N775K | 48.64 | 0.39 | 3.07 | 0.43 | 66.62 | 0.10 | 4.03 | 0.14 |
| S152Y;L453F;N775K;S523P | 45.87 | 1.87 | -0.05 | 1.88 | 66.91 | 0.05 | 4.58 | 0.08 |
| L453F;V671F;N775K;S523P | 50.09 | 0.85 | 4.52 | 0.87 | 67.84 | 0.07 | 5.25 | 0.12 |
| S152Y;L453F;V671F;N775K | 49.31 | 0.18 | 4.50 | 0.22 | 66.95 | 0.11 | 5.30 | 0.26 |
| S152Y;L453F;V671F;N775K; N694H | 49.09 | 0.51 | 3.52 | 0.54 | 68.13 | 0.30 | 5.54 | 0.32 |
| S152Y;L453F;V671F;N775K; C660A | 49.10 | 0.51 | 3.53 | 0.54 | 66.62 | 0.07 | 4.03 | 0.12 |
| S152Y;L453F;V671F;N775K; S523P | 48.38 | 2.64 | 2.47 | 2.65 | 68.06 | 0.09 | 5.73 | 0.11 |
| S152Y;A395M;L453F;V671F; N775K;S523P | ND | ND | ND | ND | 68.97 | 0.06 | 6.41 | 0.14 |

SD refers to standard deviation of triplicates (*n*=3). ND stands for not detected. Shaded row refers to *Gm*SuSy-97, which yielded the best combined mutant in Tm_app_.

**Table S3.** Relative and residual activities for combined mutants from consensus-guided engineering.

| **Mutation** | **Activity** | | | |
| --- | --- | --- | --- | --- |
|  | **Relative (%)** | **SD (%)** | **Residual (%)** | **SD (%)** |
| C660A;N694H | 142.8 | 24.3 | 39.1 | 13.8 |
| Q754K;C660A | 132.4 | 22.2 | 34.1 | 14.1 |
| C660A;S523P | 168.4 | 26.2 | 78.4 | 22.8 |
| N694H;Q754K | 134.4 | 24.0 | 31.9 | 12.0 |
| N694H;S523P | 192.4 | 40.8 | 94.6 | 28.6 |
| Q754K;S523P | 152.6 | 25.8 | 76.6 | 21.0 |
| H534Y;S523P | 179.4 | 32.2 | 69.1 | 17.8 |
| A395M;S523P | 155.7 | 23.8 | 74.4 | 17.3 |
| S152Y;N775K | 109.4 | 12.9 | 67.3 | 15.4 |
| L453F;V671F | 94.8 | 11.9 | 43.8 | 6.4 |
| S152Y;N775K | 60.7 | 7.8 | 24.0 | 6.1 |
| L453F;N775K | 133.4 | 14.7 | 112.5 | 26.6 |
| S152Y; V671F | 63.4 | 10.3 | 15.9 | 4.2 |
| V671F;N775K | 72.9 | 9.6 | 21.5 | 7.1 |
| N694H;Q754K;S523P | 204.0 | 51.1 | 100.5 | 11.2 |
| L453F;V671F;N775K | 83.6 | 8.4 | 55.6 | 7.7 |
| S152Y;L453F;N775K | 81.4 | 5.1 | 64.1 | 8.1 |
| S152Y;L453F;V671F | 59.1 | 12.5 | 35.0 | 7.1 |
| A395M;N694H;Q754K;S523P | 207.3 | 43.3 | 146.3 | 23.5 |
| A395M;L453F;V671F;N775K | 128.5 | 22.5 | 80.4 | 22.3 |
| S152Y;L453F;N775K;S523P | 178.0 | 24.4 | 138.0 | 14.3 |
| L453F;V671F;N775K;S523P | 141.2 | 22.4 | 122.3 | 26.5 |
| S152Y;L453F;V671F;N775K | 73.4 | 7.2 | 54.5 | 6.5 |
| S152Y;L453F;V671F;N775K;N694H | 104.1 | 16.4 | 81.3 | 19.6 |
| S152Y;L453F;V671F;N775K;C660A | 95.7 | 21.6 | 152.5 | 33.1 |
| S152Y;L453F;V671F;N775K;S523P | 141.0 | 10.7 | 106.0 | 7.8 |
| S152Y;A395M;L453F;V671F;N775K;S523P | 178.0 | 40.1 | 165.9 | 31.3 |

SD refers to standard deviation of triplicates (*n*=3). Shaded row refers to *Gm*SuSy-97, which yielded the best combined mutant in both relative and residual activity.

**Table S4.** Amino acid sequences of SuSy wildtype and variants designed in this study.

| **SuSy variant** | **Amino acid sequence** |
| --- | --- |
| WT | MATDRLTRVHSLRERLDETLTANRNEILALLSRIEAKGKGILQHHQVIAEFEEIPEENRQKLTDGAFGEVLRSTQEAIVLPPWVALAVRPRPGVWEYLRVNVHALVVEELQPAEYLHFKEELVDGSSNGNFVLELDFEPFNAAFPRPTLNKSIGNGVQFLNRHLSAKLFHDKESLHPLLEFLRLHSVKGKTLMLNDRIQNPDALQHVLRKAEEYLGTVPPETPYSEFEHKFQEIGLERGWGDNAERVLESIQLLLDLLEAPDPCTLETFLGRIPMVFNVVILSPHGYFAQDNVLGYPDTGGQVVYILDQVRALENEMLHRIKQQGLDIVPRILIITRLLPDAVGTTCGQRLEKVFGTEHSHILRVPFRTEKGIVRKWISRFEVWPYLETYTEDVAHELAKELQGKPDLIVGNYSDGNIVASLLAHKLGVTQCTIAHALEKTKYPESDIYWKKLEERYHFSCQFTADLFAMNHTDFIITSTFQEIAGSKDTVGQYESHTAFTLPGLYRVVHGIDVFDPKFNIVSPGADQTIYFPHTETSRRLTSFHPEIEELLYSSVENEEHICVLKDRSKPIIFTMARLDRVKNITGLVEWYGKNAKLRELVNLVVVAGDRRKESKDLEEKAEMKKMYGLIETYKLNGQFRWISSQMNRVRNGELYRVICDTRGAFVQPAVYEAFGLTVVEAMTCGLPTFATCNGGPAEIIVHGKSGFHIDPYHGDRAADLLVDFFEKCKLDPTHWDKISKAGLQRIEEKYTWQIYSQRLLTLTGVYGFWKHVSNLDRRESRRYLEMFYALKYRKLAESVPLAAE |
| *Gm*SuSy-97 | MATDRLTRVHSLRERLDETLTANRNEILALLSRIEAKGKGILQHHQVIAEFEEIPEENRQKLTDGAFGEVLRSTQEAIVLPPWVALAVRPRPGVWEYLRVNVHALVVEELQPAEYLHFKEELVDGSSNGNFVLELDFEPFNAAFPRPTLNKYIGNGVQFLNRHLSAKLFHDKESLHPLLEFLRLHSVKGKTLMLNDRIQNPDALQHVLRKAEEYLGTVPPETPYSEFEHKFQEIGLERGWGDNAERVLESIQLLLDLLEAPDPCTLETFLGRIPMVFNVVILSPHGYFAQDNVLGYPDTGGQVVYILDQVRALENEMLHRIKQQGLDIVPRILIITRLLPDAVGTTCGQRLEKVFGTEHSHILRVPFRTEKGIVRKWISRFEVWPYLETYTEDVMHELAKELQGKPDLIVGNYSDGNIVASLLAHKLGVTQCTIAHALEKTKYPESDIYWKKFEERYHFSCQFTADLFAMNHTDFIITSTFQEIAGSKDTVGQYESHTAFTLPGLYRVVHGIDVFDPKFNIVPPGADQTIYFPHTETSRRLTSFHPEIEELLYSSVENEEHICVLKDRSKPIIFTMARLDRVKNITGLVEWYGKNAKLRELVNLVVVAGDRRKESKDLEEKAEMKKMYGLIETYKLNGQFRWISSQMNRVRNGELYRVICDTRGAFVQPAFYEAFGLTVVEAMTCGLPTFATCNGGPAEIIVHGKSGFHIDPYHGDRAADLLVDFFEKCKLDPTHWDKISKAGLQRIEEKYTWQIYSQRLLTLTGVYGFWKHVSKLDRRESRRYLEMFYALKYRKLAESVPLAAE |
| Anc165 | MAERALTRVHSLRERLDETLSAHRNEILALLSRIESHGKGILQPHQLLAEFEAIPEENRKKLTDGAFGEVLRSTQEAIVLPPWVALAVRPRPGVWEYIRVNVNALAVEELTVAEYLHFKEELVDGSSNGNFVLELDFEPFNASFPRPTLSKYIGNGVEFLNRHLSAKLFHDKDSMHPLLDFLRVHHYKGKTMMLNDRIQNLNALQSVLRKAEEYLSTLPPETPYSEFEHKFQEIGLERGWGDTAERVLEMIQLLLDLLEAPDPCTLEKFLGRIPMVFNVVILSPHGYFAQDNVLGYPDTGGQVVYILDQVRALENEMLHRIKQQGLDITPRILIVTRLLPDAVGTTCGQRLEKVYGTEHSHILRVPFRTEKGIVRKWISRFEVWPYLETYTEDVMNEIAGELQAKPDLIIGNYSDGNIVASLLAHKLGVTQCTIAHALEKTKYPDSDIYWKKFEDKYHFSCQFTADLIAMNHTDFIITSTFQEIAGSKDTVGQYESHTAFTLPGLYRVVHGIDVFDPKFNIVPPGADMSIYFPYTEEKKRLTSLHPEIEELLYSDVENEEHLCVLKDRNKPIIFSMARLDRVKNITGLVEWYGKNARLRELVNLVVVAGDRRKESKDLEEKAEMKKMYELIETYKLNGQFRWISSQMNRVRNGELYRYIADTKGAFVQPAFYEAFGLTVVEAMTCGLPTFATCHGGPAEIIVHGKSGFHIDPYHGDQAAELLVDFFEKCKADPSHWDKISQGGLKRIEEKYTWKIYSERLLTLAGVYGFWKYVSKLDRRETRRYLEMFYALKYRKLAESVPLAVEE |
| HPMN251 | MAERALTRVHSLRERLDEELSAHRNEILLLLSRIESHGKGILQPHQLLAEFEALPEELRKRLRDGAFGEVLRHTQEAIVLPPWVALAVRPRPGVWEYIRVNVNTLAVEALTVAEYLAFKEELVDGPSNGNFVLELDFEPFNKSFPRPTLSKYIGNGVEFLNRHLSAKLFHDKDSMHPLLDFLRVHHYKGEKMMLNDRIQNLNQLQSVLRKAEEYLSTLPPETPYSEFEEYFQEIGLERGWGDTAERVLEMIQLLLDLLEAPDPQTLEKFLGRIPMVFNVVILSPHGYFAQDNVLGYPDTGGQVVYILDQVRALENEMLHRIKQQGLDITPRILIVTRLLPDAVGTTCGQRLEKVYGTEHSHILRVPFRTEKGIVRKWISRFEVWPYLETYTEDVMNEIAGELQAKPDLIIGNYSDGNIVASLLAHKLGVTQCTIAHALEKTKYPDSDIYWKKFEDKYHFSCQFTADLIAMNHTDFIITSTFQEIAGSKDTVGQYESHHAFTLPGLYRVVHGIDVFDPKFNIVPPGADMSIYFPYTEEKKRLTELHPEIEELLYSDVENKEHLCTLKDRNKPIIFSMARLDSVKNITGLVEWYGKNARLRELVNLVVVAGDRRKESKDLEEKAEMKKMYELIETYKLNGQFRWISSQMNRVRNGELYRYIADTKGAFVQPAFYEAFGLTVVEAMTCGLPTFATCHGGPAEIIVHGKSGFHIDPYHGDQAAELLVDFFEKCKADPSHWDKISQGGLKRIEEKYTWKIYSERLLTLAGVYGFWKYVSKLDRRETRRYLEMFYALKYRKLAESVPLAVEE |
| HPMN | MATDRLTRVHSLRERLDEELEANRNEILLLLSRIEARGKGILQHHQVIAEFEALPEELRKRLRDGAFGEVLRHTQEAIVLPPWVALAVRPRPGVWEYLRVNVHTLKVEALQPAEYLAFKEELVDGPSNGNFVLELDFEPFNKNFPRPTLNKYIGNGVQFLNRHLSAKLFHDKESLHPLLEFLRSHSVKGEKLMLNDRIQNPDQLQEVLRKAEEYLKTVPPETPYSEFEEYFQEIGLERGWGDNAERVLESIQLLLDLLEAPDPQTLETFLGRIPMVFNVVILSPHGYFAQDNVLGYPDTGGQVVYILDQVRALENEMLHRIKQQGLDIVPRILIITRLLPDAVGTTCGQRLEKVSGTEHSHILRVPFRTEKGIVRKWISRFEVWPYLETYTEDVMEELAKELQGKPDLIIGNYSDGNIVASLLAHKLGVTQCTIAHALEKTKYPESDIYWKKFEERYHFSCQFTADLFAMNHTDFIITSTFQEIAGSKDTVGQYESHHAFTLPGLYRVVHGIDVFDPKFNIVPPGADQSIYFPHTETSRRLTEFHPEIEELLYSEVENKEHICTLKDRSKPIIFTMARLDSVKNITGLVEWYGKNAKLRELVNLVVVAGDRRKESKDLEEKAEMKKMYELIETYKLNGQFRWISSQMNRVRNGELYRVIADTRGAFVQPAFYEAFGLTVVEAMTCGLPTFATCNGGPAEIIVHGKSGFHIDPYHGDRAADLLVDFFEKCKKDPTHWDKISKAGLQRIEEKYTWKIYSQRLLTLTGVYGFWKHVSKLDRRESRRYLEMFYALKYRKLAESVPLAAE |
| Anc294 | MATSRLTRVLSLRERLDETLTAHRNEILALLSRIEQGNKNGGVEFLNRHLSAKMFHDSMHPLLDFLRMHHYKGKTMMLNDRIQNLDSLQSVLRKAEEYLTTIPPDTPYSEFEHKFQEIGLERGWGDMIQLLLDLLEAPDPCTLEKFLGRIFNVVILSPHGYFAQANVALGYDPDTGGQVVYILDQVRALENEMLHRIKQQGLDIIVTRLLPDAVGTTCGQRLEKVFGHSHILRVPFRTEKGIVRKWISRFEVWPYLETYTEDVMNEIAAEQAKPDLIIGNYSDGNIVASLLAHKLGVTQCTIAHALEKTKYPDSDIYWKKLEEKYHFSCQFTADLIAMNHTDFIITSTFQEIAGSKDTVGQYESHTAFPKFNIVPPGADMSIYFPHTEKKRRLTALLYSSVENVEEHICVLKDRSKPIIFSMARLDRVKNITGLVEWYGKNAKLRELVNLVVVAGDRRKESKDLEEKAEMKKMYSLIEEYKLNGQFRWISSQMNRVRNGELYRVIADTRGAFVQPAFYEAFGLTVVEAMTCGLPTFATCHGGPAEIIVHGKSGFHIDPYHGDKAADLLVDFFEKCKKDPSHWDKISKGGLQRIEEKYTWKIYSDRLLTLAGVYGFWKYVSKLDRRETRRYLEMFYALKYRKLAESVPLAIDD |
| Anc290 | MATSRLTRVLSMRERVEDTLSAHRNELVSLLSRYEQGDRNDGVQFLNRHLSSKLFHDSMQPLLDFLRTHKYKGQTMMLNDRIQSLSGLQSALVKAEEYLSKLPPDTPYSEFEHKFQEMGLEKGWGDMIHLLLDILQAPDPSTLEKFLGRIFNVVILSPHGYFGQANVALGLDPDTGGQVVYILDQVRALENEMLHRIKQQGLDIVVTRLIPDAKGTTCNQRLEKISGHTHILRVPFRTEKGIVRQWISRFDVWPYLETFTEDVMNEIAAEQGKPDLIIGNYSDGNLVASLLAHKLGVTQCTIAHALEKTKYPDSDIYWKKLDEKYHFSCQFTADLIAMNHADFIITSTYQEIAGSKDTVGQYESHTAFPKFNIVPPGADMSIYFPYTEKQKRLTALLYSPEQNVDEHICVLNDRKKPIIFSMARLDRVKNITGLVEWYGKNAKLRELVNLVVVAGYIDVKKSKDREEIAEIEKMHNLIKEYNLNGQFRWICSQTNRVRNGELYRYIADTRGAFVQPAFYEAFGLTVVEAMTCGLPTFATCHGGPAEIIVHGVSGFHIDPYHGDQAAELMVDFFEKCKKDPSHWDKISEGGLQRIYERYTWKIYSERLMTLAGVYGFWKYVSKLERRETRRYLEMFYILKFRDLAKSVPLAVDDPA |
| Anc180 | MATRRLTRVHSLRERLDETLTAHRNEILALLSRIEAKGKGILQHHQLIAEFEEIPEENRQKLTDGAFGEVLRSTQEAIVLPPWVALAVRPRPGVWEYLRVNVHALVVEELQAAEYLHFKEELVDGSSNGNFVLELDFEPFNASFPRPTLNKYIGNGVEFLNRHLSAKLFHDKESMHPLLEFLRLHSYKGKNLMLNDRIQNLNALQHVLRKAEEYLSTLAPETPYSEFEHKFQEIGLERGWGDTAERVLEMIQLLLDLLEAPDPCTLETFLGRIPMVFNVVILSPHGYFAQDNVLGYPDTGGQVVYILDQVRALENEMLHRIKQQGLDITPRILIITRLLPDAVGTTCGQRLEKVYGTEHSHILRVPFRTEKGIVRKWISRFEVWPYLETYTEDVMHELAKELQGKPDLIIGNYSDGNIVASLLAHKLGVTQCTIAHALEKTKYPESDIYWKKFEDKYHFSCQFTADLFAMNHTDFIITSTFQEIAGSKDTVGQYESHTAFTLPGLYRVVHGIDVFDPKFNIVPPGADMSIYFPYTETKRRLTSFHPEIEELLYSSVENEEHICVLKDRNKPIIFTMARLDRVKNITGLVEWYGKNARLRELVNLVVVAGDRRKESKDLEEKAEMKKMYGLIETYKLNGQFRWISSQMNRVRNGELYRVIADTKGAFVQPAFYEAFGLTVVEAMTCGLPTFATCNGGPAEIIVHGKSGFHIDPYHGDQAADLLVDFFEKCKADPSHWDKISQGGLQRIEEKYTWKIYSERLLTLTGVYGFWKHVSKLDRRESRRYLEMFYALKYRKLAESVPLAVE |
| Anc111 | MATDALTRVHSLRERLDETLTANRNEILALLSRIEAKGKGILQHHQVIAEFEEIPEENRQKLTDGAFGEVLRSTQEAIVLPPWVALAVRPRPGVWEYLRVNVHALVVEELQPAEYLHFKEELVDGSSNGNFVLELDFEPFNASFPRPTLNKYIGNGVQFLNRHLSAKLFHDKESLHPLLLEFLRLHSYKGKTLMLNDRIQNPDALQHVLRKAEEYLGTVAPETPYSEFEHKFQEIGLERGWGDTAERVLESIQLLLDLLEAPDPCTLETFLGRIPMVFNVVILSPHGYFAQDNVLGYPDTGGQVVYILDQVRALENEMLHRIKQQGLDIVPRILIITRLLPDAVGTTCGQRLEKVYGTEHCHILRVPFRTEKGIVRKWISRFEVWPYLETYTEDVMHELAKELQGKPDLIIGNYSDGNIVASLLAHKLGVTQCTIAHALEKTKYPESDIYWKKFEEKYHFSCQFTADLFAMNHTDFIITSTFQEIAGSKDTVGQYESHTAFTLPGLYRVVHGIDVFDPKFNIVPPGADQSIYFPYTETSRRLTSFHPEIEELLYSSVENEEHICVLKDRNKPIIFTMARLDRVKNITGLVEWYGKNAKLRELVNLVVVAGDRRKESKDLEEKAEMKKMYGLIETYKLNGQFRWISSQMNRVRNGELYRVIADTKGAFVQPAFYEAFGLTVVEAMTCGLPTFATCNGGPAEIIVHGKSGFHIDPYHGDRAADLLVDFFEKCKADPSHWDKISQGGLQRIEEKYTWKIYSQRLLTLTGVYGFWKHVSKLDRRSRRYLEMFYALKYRKLAESVPLAVEE |
| Anc101 | MAERALTRVHSLRERLDETLSAHRNEILALLSRIEAKGKGILQHHQLIAEFEAIPEENRQKLLDGAFGEVLRSTQEAIVLPPWVALAVRPRPGVWEYIRVNVHALVVEELQVAEYLHFKEELVDGSSNGNFVLELDFEPFNASFPRPTLSKYIGNGVEFLNRHLSAKLFHDKESMHPLLLEFLRVHCYKGKTMMLNDRIQNLNALQHVLRKAEEYLSTLAPETPYSEFEHKFQEIGLERGWGDTAERVLEMIQLLLDLLEAPDPCTLEKFLGRIPMVFNVVILSPHGYFAQDNVLGYPDTGGQVVYILDQVRALENEMLHRIKQQGLDITPRILIITRLLPDAVGTTCGQRLEKVFGTEHSHILRVPFRTEKGIVRKWISRFEVWPYLETYTEDVMHELAKELQGKPDLIIGNYSDGNIVASLLAHKLGVTQCTIAHALEKTKYPDSDIYWKKFDEKYHFSCQFTADLIAMNHTDFIITSTFQEIAGSKDTVGQYESHTAFTLPGLYRVVHGIDVFDPKFNIVPPGADMSIYFPYTETEKRLTSFHPEIEELLYSSVENEEHLCVLKDRNKPIIFTMARLDRVKNITGLVEWYGKNAKLRELVNLVVVAGDRRKESKDLEEKAEMKKMYGLIETYKLNGQFRWISSQMNRVRNGELYRYIADTKGAFVQPAFYEAFGLTVVEAMTCGLPTFATCNGGPAEIIVHGKSGFHIDPYHGDQAAEILVDFFEKCKADPSHWDKISQGGLQRIHEKYTWKIYSERLLTLTGVYGFWKHVSKLDRLSRRYLEMFYALKYRKLAESVPLAVEE |
| T-1 | MATDRLTRVHSLRERLDEDLERNREEIALLLRRIAAQGKGILQHHEVMAIFESLPEELRERLRNGAFGEVLESTVEAVVLPPYVALLVRPRPGVWEYLRIHVETLEVEELTPAEYLAFKEELVLGPSNGSPVLEIDFSPFRKNEPRPTDVKYIGNGTEFLNRHLSEELFKDKESLHPLLDFLKSHEVNGEKLMLNDKIKNPDELEAALREALERLATLPEDTPYEEFLEWFQSIGLERGWGDNAGRVRESIQLLLDLLEAPDPETLEEFLSRIPMVFNVVIISPHGYFAQENVLGYPDTGGQVVYILDQVRALEKEMERRIKEQGLDITPRILIVTRLLPDAEGTTCGQRLEKVKGTKHSYILRVPFRNEKGIVRKWISRFEVWPYLERYTEDVLEELKKELNGKPDLIVGNYSDGNIVASLLAKKLNVTQCTIAHALEKTKYPYSDIYWKELEARYHFSCQFTADLFAMNSTDFIITSTFQEIAGSETTVGQYESHRTFTLPGLYRVVNGIDVFDPKFNIVPPGADESIYFPHTLKERRRDEYHPEIHELLYSTVESDESICTLKDKSKPIIFTMARLDSVKNITGLVEWYGKNEELRKLVNLVVVAGDRRVESTDEEEREEMEKMYELIKKYNLDGQFRWISSQMNRVRNGELYREICDTRGAFVQPALYEAFGLTVVEAMRCGLPTFATNNGGPAEIIEHGVSGFHIDPYHPDAAAQLLVDFFKKCEEDPSHWDEISDAGLARIRANYTWENYSRRLLRLANVYGFWKHYSKDSRVELDRYLEMFYNLLYRPLAAAVPLAAE |
| T-2 | MATDRLTRVHSLRERLDEDLARNREAIARLLAALAAQGKGILQHHQVLALFEALPPEEQERLRRGAFGEALEHTQEAVVLPPYIALLIRPRPGVWEYVRLHVETLRLEALSPAEYLAFKEELVLGPSNGEEVLEIDFRPFRKNEPRPTDVKYIGNGTEFLNRHLSEELFKDKESLHPLLDFLKSHEVNGEKLMLNDKIQNPDQLEAVLRKAAEFLDTLPADTPYEEFLEYFQSIGLERGWGANAARVRESIRLLLDLLEAPDPETLEEFLSRIPTVFNVVIISPHGYFAQENVLGYPDTGGQVVYILDQVRALEEEMVRRIEEQGLDITPRILIVTRLLPDAEGTTCGQRLEKVAGTEHSFILRVPFRNENGIVRKWISRFEVWPYLERYTEDVLEELAKELNGKPDLIVGNYSDGNIVASLLAKKLNVTQCTIAHALEKTKYPYSDIYWKELEERYHFSCQFTADLFAMNYTDFIITSTFQEIAGSETTVGQYESHHTFTLPGLYRVVHGIDVFDPKFNIVPPGADSSIYFPHTLEERRRTEYHPEIDELLYSEVESDEHVCRLKDRSKPIIFTMARLDSVKNITGLVEWYGKNEKLRELVNLVVVAGDRRRESTDPEERAEMEKMYELIKKYNLDGQFRWISSQMNRVRNGELYREICDTRGAFVQPALYEAFGLTVVEAMRCGLPTFATNNGGPAEIIEHGVSGFHIDPYHPDEAAEKLVEFFEKCEEDPSHWDRISDAGLKRIEENYTWENYSRRLLSLANVYGFWKHYSKDARKELERYLEMFYNLLYRPLAASVPLAAE |
| T-3 | MATDRLTRVHSLRERLDEELERNRDAIKLLLERMAARGKGILQHHEVLAVFESLPPEDRERLSSGAFGEALASTQEAVILPPYIALLIRPRPGVWEYLRIHVETLKVEALTPAEYLRFKEELVLGPSNGTPVLEIDFAPFRKNIPRPTDVKYIGNGIEFLNRHLSEELFANKESLHPLLDFLRSHEVNGEKLMLNDKIQNPDELEKALKEALQKLATLPADTPYTKFLDYFQSIGLERGWGNNAARVKESIELLLDLLEAPDPETLEEFLSRIPIVFNVVIISPHGYFAQENVLGYPDTGGQVVYILDQVRALEREMERRIADQGLDITPRILIVTRLLPDAQGTTCGQRLEKVPGTQHSYILRVPFRNEKGIVRKWISRFEVWPYLERYTEDVLEELEKELDGKPDLIVGNYSDGNIVASLLAKKLGVTQCTIAHALEKTKYPHSDIYWKDLEARYHFSCQFTADLFAMNYTDFIITSTFQEIAGSATTIGQYESHHSFTLPGLYRVVNGIDVFDPKFNIVPPGADSSIYFPHTDRARRRDEYHPEIDNLLYSSVESEEQKCTLKDRTKPIIFTMARLDSVKNITGLVEWYGKNEELRKLVNLVVVAGDLRRESEDEEERKEMEKMYELIEKYNLDGQFRWISSQMNRVRNGELYRHICDHRGAFVQPALYEAFGLTVVEAMRCGLPTFATNNGGPAEIIDHGVSGFHIDPYHPDEAAEKLVEFFTKCREDPSHWDEISAAGLARINAHYTWEIYSERLLKLASVYGFWKHYSKNERTELERYLEMFYELLYRPLAASVPLAAE |
| *Ac*SuSy (L637M-T640V) | MIEALRQQLLDDPRSWYAFLRHLVASQRDSWLYTDLQRACADFREQLPEGYAEGIGPLEDFVAHTQEVIFRDPWMVFAWRPRPGRWIYVRIHREQLALEELSTDAYLQAKEGIVGLGAEGEAVLTVDFRDFRPVSRRLRDESTIGDGLTHLNRRLAGRIFSDLAAGRSQILEFLSLHRLDGQNLMLSNGNTDFDSLRQTVQYLGTLPRETPWAEIREDMRRRGFAPGWGNTAGRVRETMRLLMDLLDSPSPAALESFLDRIPMISRILIVSIHGWFAQDKVLGRPDTGGQVVYILDQARALEREMRNRLRQQGVDVEPRILIATRLIPESDGTTCDQRLEPVVGAENVQILRVPFRYPDGRIHPHWISRFKIWPWLERYAQDLEREVLAELGSRPDLIIGNYSDGNLVATLLSERLGVTQCNIAHALEKSKYLYSDLHWRDHEQDHHFACQFTADLIAMNAADIIVTSTYQEIAGNDREIGQYEGHQDYTLPGLYRVENGIDVFDSKFNIVSPGADPRFYFSYARTEERPSFLEPEIESLLFGREPGADRRGVLEDRQKPLLLSMARMDRIKNLSGLAELYGRSSRLRGLANLVIIGGHVDVGNSRDAEEREEIRRMHEIMDHYQLDGQLRWVGALMDKVVAGELYRVVADGRGVFVQPALFEAFGLTVIEAMSSGLPVFATRFGGPLEIIEDGVSGFHIDPNDHEATAERLADFLEAARERPKYWLEISDAALARVAERYTWERYAERLMTIARIFGFWRFVLDRESQVMERYLQMFRHLQWRPLAHAVPME |

**Table S5.** Initial characterization of selected SuSy ancestors

| **SuSy variant** | **Thermostability** | | | | **Activity** | | | |
| --- | --- | --- | --- | --- | --- | --- | --- | --- |
|  | **Tm_app_ (°C)** | **SD (°C)** | **ΔTm_app_ (°C)** | **SD (°C)** | **Relative (%)** | **SD (%)** | **Residual (%)** | **SD (%)** |
| WT | 62.62 | 0.09 | 0 | - | 100.0 | 15.6 | 17.1 | 3.6 |
| Ancestor 180 | 69.85 | 0.17 | 7.23 | 0.16 | 126.9 | 31.7 | 104.9 | 29.4 |
| Ancestor 165 | 74.37 | 0.06 | 11.75 | 0.09 | 165.7 | 31.8 | 158.3 | 31.1 |
| Ancestor 111 | 73.29 | 0.06 | 10.67 | 0.09 | 127.4 | 25.9 | 122.1 | 27.2 |
| Ancestor 101 | 72.20 | 0.03 | 9.58 | 0.08 | 127.4 | 20.8 | 119.0 | 20.9 |

Anc290 and Anc294 were not expressed in soluble form and therefore were discarded. SD refers to standard deviation of triplicates (*n*=3). Ancestor 165 indicates Anc165 as described in the main manuscript.

**Table S6.** Protein yield after purification process expressed as mg of protein per liter of culture.

| **Variant** | **Yield (mg L^-1^)** |
| --- | --- |
| WT | 6.9 |
| *Gm*SuSy-97 | 5.8 |
| Ancestor 165 | 5.5 |
| HPMN | 32.0 |
| HPMN251 | 14.5 |
| AcSuSy_L637M/T640V_ | 0.1 |

**Table S7.** Experimental characterization of ProteinMPNN variants.

| **SuSy variant** | **Thermostability** | | | | **Activity** | | | |
| --- | --- | --- | --- | --- | --- | --- | --- | --- |
|  | **Tm_app_ (°C)** | **SD (°C)** | **ΔTm_app_ (°C)** | **SD (°C)** | **Relative (%)** | **SD (%)** | **Residual (%)** | **SD (%)** |
| WT | 62.62 | 0.09 | 0 | - | 100.0 | 15.6 | 17.1 | 3.6 |
| HPMN | 70.69 | 0.05 | 8.07 | 0.11 | 87.0 | 20.8 | 91.5 | 20.9 |
| HPMN251 | 75.92 | 0.10 | 13.30 | 0.13 | 160.3 | 20.0 | 133 | 20.9 |

SD refers to standard deviation of triplicates (*n*=3).

**Table S8.** Analysis of mutations on Anc165 and their possible effect on molecular interactions.

| **Domain** | **Mutation** | **Predicted effect on molecular interactions** |
| --- | --- | --- |
| CTD | T3E | Neutral/No evident effect |
|  | D4R | Neutral/No evident effect |
|  | R5A | Neutral/No evident effect |
|  | T21S | Neutral/No evident effect. Located at the protein surface |
|  | N23H | Possible salt bridge formation |
|  | A36S | Hydrogen bond formation |
|  | K37H | Improvement in hydrophobic interactions |
|  | H44P | Loss of hydrogen bond. Gain in rigidity |
|  | V47L | Neutral/No evident effect |
|  | I48L | Neutral/No evident effect |
|  | E53A | Loss of salt bridge |
|  | Q60K | Possible salt bridge formation |
|  | L98I | Neutral/No evident effect |
|  | H103N | Loss of salt bridge. Hydrogen bond formation |
|  | V106A | Neutral/No evident effect. Located at the protein surface |
|  | Q111T | Neutral/No evident effect |
|  | P112V | Loss of rigidity |
|  | A143S | Hydrogen bond formation |
|  | N150S | Neutral/No evident effect |
| Linker CTD - EPDB | S152Y | Improvement in hydrophobic interactions. At the tetramer interface |
| EPDB | Q158E | Neutral/No evident effect |
|  | E173D | Stronger salt bridge |
|  | L175M | Improvement in hydrophobic interactions |
|  | E180D | Neutral/No evident effect |
|  | L184V | Loss of hydrophobic interactions |
|  | S186H | Hydrogen bond formation and improvement in hydrophobic interactions |
|  | V187Y | Hydrogen bond formation and improvement in hydrophobic interactions |
|  | L192M | Neutral/No evident effect |
|  | P201L | Improvement in hydrophobic interactions |
|  | D202N | Loss of salt bridge |
|  | H206S | No evident effect |
|  | G216S | Hydrogen bond formation |
|  | V218L | Neutral/No evident effect |
|  | N243T | Loss of hydrogen bond |
|  | S250M | Improvement in hydrophobic interactions |
|  | T268K | Hydrogen bond formation |
| GT-B fold N-terminal | V329T | Neutral/No evident effect |
|  | I335V | Neutral/No evident effect |
|  | F355Y | Hydrogen bond formation |
|  | A395M | Improvement in hydrophobic interactions |
|  | H396N | Loss of salt bridge |
|  | L398I | Neutral/No evident effect |
|  | K400G | Gain in flexibility |
|  | G404A | Loss of flexibility |
|  | V410I | Neutral/No evident effect |
|  | E445D | Stronger hydrogen bond |
|  | L453F | Improvement in hydrophobic interactions |
|  | E455D | Neutral/No evident effect |
|  | R456K | Neutral/No evident effect |
|  | F468I | Weaker hydrophobic interactions |
| GT-B fold - Hinge | S523P | Gain in rigidity |
|  | Q528M | Loss of hydrogen bond |
|  | T529S | Neutral/No evident effect. Facing the solvent |
|  | H534Y | Stronger hydrophobic interactions and hydrogen bonding |
|  | T537E | Neutral/No evident effect. Facing the solvent |
|  | S538K | Neutral/No evident effect. Facing the solvent |
|  | R539K | Weaker salt bridge |
|  | Q754K | Neutral/No evident effect. Located at the protein surface |
| GT-B fold C-terminal | F544L | Neutral/No evident effect |
|  | S555D | Neutral/No evident effect. Located at the protein surface |
|  | I562L | Neutral/No evident effect |
|  | S569N | Weaker hydrogen bond |
|  | T575S | Improvement in hydrophobic interactions |
|  | K597R | Stronger salt bridge |
|  | G629E | Salt bridge formation |
|  | V658Y | Improvement in hydrophobic interactions, hydrogen bonding |
|  | C660A | Neutral/No evident effect |
|  | R663K | Loss of salt bridge |
|  | V671F | Improvement in hydrophobic interactions |
|  | N694H | Improvement in hydrophobic interactions, salt bridge formation |
|  | R717Q | Neutral/No evident effect |
|  | D720E | Salt bridge formation |
|  | L731A | Neutral/No evident effect |
|  | T734S | Neutral/No evident effect |
|  | K741Q | Neutral/No evident effect. Located at the protein surface |
|  | A742G | Neutral/No evident effect. Located at the protein surface |
|  | Q745K | Neutral/No evident effect. Located at the protein surface |
|  | Q758E | Neutral/No evident effect |
|  | T764A | Improvement in hydrophobic interactions |
|  | H772Y | Loss of salt bridge |
|  | N775K | Neutral/No evident effect. Located at the protein surface |
|  | S781T | Neutral/No evident effect |
|  | A804V | Neutral/No evident effect |
|  | Insertion 806 | Longer C-terminal interacting to close the latch |

*Gm*SuSy comprises four main structural regions: a cellular targeting domain (CTD), an ENOD40 peptide-binding domain (EPDB), and two GT-B fold domains (GT-B C-terminal and N-terminal)^1,2^. Domains were assigned by comparison with the AtSuSy crystal structure (PDB: 3S27)^1^. CTD comprises residues 1-125, EPBD comprises residues 155–274, and the GT-B fold region comprises residues 275–805. The CTD and EPBD are connected by a linker spanning residues 126-154. The hinge is centered around residues 525 and 752, with associated coils extending approximately across residues 521-532 and 750-756, respectively. This hinge connects the N- and C-terminal of the GT-B fold.

**Table S9.** Residues involved in the SPM of tetrameric SuSy variants.

| **WT – Total residues: 254** | |
| --- | --- |
| **Domain** | **Residues** |
| **CTD and EPDB domains** | 39 (D), 67 (A), 70 (A), 73 (A), 79 (D), 84 (D), 86 (C), 88 (B,C,D), 89 (B,C,D), 91 (B,C,D), 92 (A,B,C,D), 98 (A,B,C,D), 99 (A,B,C,D), 100 (B), 101 (B), 110 (A,B,C,D), 111 (A,B,C,D), 113 (A,B), 114 (A,B,C,D), 117 (A,B,C,D), 120 (A,B,C,D), 135 (A,B,C,D), 136 (A,B,C,D), 137 (A,B,C,D), 138 (A,C,D), 139 (A,B,C,D), 140 (A,C,D), 142 (A), 143 (A), 144 (A), 145 (A), 146 (A), 147 (A), 148 (A), 149 (A), 152 (A), 154 (B,C,D), 155 (A,B,C,D), 156 (A,B,C,D), 157 (B,C,D), 158 (A), 159 (A,B,C,D), 160 (B,C,D), 161 (A), 162 (A,B,C,D), 163 (A,B,C,D), 165 (A,B,C,D), 166 (A,B,C,D), 167 (B), 168 (B,C), 169 (A), 174 (B,C), 177 (B,C), 240 (B), 241 (A,B,C), 247 (A,B,C), 248 (B), 249 (C), 250 (A,B,C), 251 (B), 252 (C), 253 (A,B,C), 254 (B), 255 (C), 256 (A,B,C), 257 (B), 258 (C), 259 (A,B), 260 (A,B,C), 261 (A,B,C), 262 (A,B,C), 263 (A,B,C,D), 266 (A,B,C,D), 267 (C), 269 (A,B,C,D), 270 (A,B,C,D), 271 (A), 272 (A,C), 273 (A,C,D), 274 (A,C,D) |
| **GT-B domain** | 275 (A,C,D), 276 (A,C,D), 280 (B), 281 (B,D), 282 (B,C,D), 312 (B,C), 313 (D), 315 (B,C), 318 (B,C), 321 (B,C), 324 (A,B,C), 326 (A,B,C,D), 328 (A,B,C,D), 329 (A,B,C,D), 330 (A,B,C,D), 331 (A,B,C,D), 332 (A,B,C,D), 333 (A,B,D), 334 (B), 338 (B), 339 (B), 362 (A), 363 (A,B), 364 (A,B), 365 (A), 366 (A), 367 (A), 374 (A), 375 (B), 376 (A,B), 377 (A,B), 378 (A,B), 379 (A,B), 382 (A,B), 384 (A,B,D), 385 (C), 386 (C), 387 (A,B,D), 390 (A,B,C,D), 391 (A,B,C,D), 407 (C), 408 (C), 409 (B), 416 (A,B,C,D), 419 (A,B,C,D), 431 (B), 448 (C), 450 (C), 454 (C), 458 (C), 461 (A,B,C,D), 464 (A,B,C,D), 465 (C,D), 467 (B), 468 (B,C,D), 469 (B,C,D), 471 (B), 472 (B,D), 473 (B,C,D), 474 (B), 475 (C), 479 (A,B), 480 (A,B,C,D), 481 (A,B,C,D), 482 (A,B,C,D), 484 (A,B), 485 (A,B,C,D), 486 (A), 487 (A), 488 (A), 494 (A), 495 (A,B,C,D), 497 (A), 498 (A,B,C,D), 501 (A,B,C,D), 502 (A,B,C,D), 503 (A,B,C,D), 508 (A,B,C,D), 509 (A,B), 510 (A,B,C,D), 511 (B,D), 512 (A,B,D), 513 (A,B,D), 516 (A,B,D), 517 (A,B,C,D), 518 (B,C), 520 (A,B,C,D), 521 (A,B,C,D), 522 (A,B,D), 523 (A,B,D), 524 (A), 525 (A,B,C,D), 526 (A,B,C,D), 573 (A,B,C,D), 594 (B), 595 (B), 596 (B), 598 (B,D), 600 (B), 602 (B), 603 (B), 604 (A,B,C,D), 640 (B), 665 (A,B,C,D), 681 (A,B,C,D), 682 (A,B,C,D), 687 (A,B,C,D), 688 (A,B,D), 689 (A), 690 (A,C,D), 691 (A,C,D), 692 (A), 693 (B), 694 (A,B), 695 (A,B), 696 (A,C,D), 697 (A,C,D), 723 (B,D), 724 (B,D), 725 (B,D), 727 (B,D), 728 (B,D), 729 (B,D), 735 (B,D), 736 (B,D), 737 (A), 738 (B,D), 739 (B,D), 740 (A,B,D), 741 (A,B,D), 742 (B), 743 (B,D), 744 (A,B,D), 746 (B,D), 747 (A,B,D), 751 (A,B,D), 752 (A,B,C,D), 753 (A,B,C,D), 754 (B), 755 (A,B,D), 756 (A,B,C,D), 757 (D), 758 (A,B,D), 759 (A,B,C,D), 760 (D), 761 (A,B,C,D), 762 (A,B,C,D), 763 (D), 764 (A,B,C,D), 765 (A,B,C,D), 766 (D), 767 (B,C,D), 768 (A,C,D), 770 (B,C,D), 771 (A,B,C,D), 773 (B,C), 774 (B,C,D), 775 (B,C), 776 (A,B,C,D), 778 (B,C), 779 (A,B,C,D), 780 (A,B,C,D), 781 (A,B,C,D), 782 (B), 783 (A,B,C,D), 784 (A,B,C,D), 786 (A,B,C,D), 787 (A,B,D), 789 (A,B,C,D), 790 (A,B,D), 793 (A,B,C,D), 794 (A,B,C,D), 796 (A,B,C,D), 797 (A,B,C,D) |
| ***Gm*SuSy-97 – Total residues: 250** | |
| **Domain** | **Residues** |
| **CTD and EPDB domains** | 6 (D), 7 (D), 8 (D), 9 (D), 10 (D), 11 (D), 14 (D), 17 (D), 18 (D), 21 (D), 24 (D), 28 (D), 31 (D), 87 (A), 88 (C,D), 89 (C,D), 91 (C,D), 92 (A,B,C,D), 97 (A), 98 (A,B,D), 99 (B), 110 (A,B,C,D), 111 (A,B,C,D), 112 (B), 113 (A,B,C,D), 114 (A,B,C,D), 117 (A,B,C,D), 120 (A,B,C,D), 135 (A,B,C,D), 136 (A,B,C,D), 137 (A,B,C,D), 138 (A,C,D), 139 (A,B,C,D), 140 (A,B,C,D), 141 (D), 142 (D), 144 (D), 145 (D), 146 (D), 147 (D), 149 (D), 151 (D), 152 (D), 153 (A,B,C), 154 (A,B,C,D), 155 (A,B,C,D), 156 (A,C,D), 157 (A,B,C,D), 158 (C,D), 159 (A,C,D), 160 (A,B,C,D), 161 (D), 162 (A,C,D), 163 (A,B,C,D), 164 (D), 165 (A,B,C,D), 166 (A,B,C,D), 167 (B), 168 (B,C,D), 171 (B), 174 (B,C,D), 176 (C), 177 (B,C,D), 210 (C), 211 (C), 214 (C), 215 (C), 218 (C), 219 (C), 222 (C), 241 (A,C), 243 (C), 247 (A,C,D), 248 (A,C), 250 (C,D), 251 (A,C), 253 (C,D), 254 (A,C), 256 (C,D), 257 (A,C), 259 (D), 260 (B,C,D), 261 (A,B,C,D), 263 (A,B,C,D), 266 (A,B,C,D), 267 (C), 269 (A,B,C,D), 270 (A,B,C,D), 271 (B,C,D), 272 (B,D), 273 (B,D), 274 (B,D) |
| **GT-B domain** | 275 (B,D), 276 (B,D), 278 (A), 279 (A), 280 (A), 281 (A,B,D), 282 (A,B,D), 313 (A,C), 316 (A,C), 318 (A,C), 321 (A,C), 324 (A,C), 326 (A,B,C), 328 (A,B,C), 329 (A,B,C), 330 (A,B,C), 331 (A,B,C), 332 (A,B,C), 333 (A,B), 334 (A,B), 384 (A,B,D), 385 (D), 386 (D), 387 (B,D), 390 (B,D), 391 (B,D), 410 (D), 414 (A), 415 (A), 416 (A,B,D), 417 (B), 419 (B,D), 432 (C,D), 447 (A,B), 450 (A,B), 451 (A,B), 454 (A,B), 458 (A,B), 459 (A), 461 (A,B,C,D), 462 (A), 463 (C), 464 (A,B,C,D), 465 (A,B), 466 (A,C), 467 (D), 468 (B,D), 469 (A,B,C), 471 (D), 472 (B), 473 (A,B,D), 475 (B), 476 (B,C,D), 479 (A,C), 480 (A,B,C,D), 481 (A,B,C,D), 482 (A,B,C,D), 484 (A,C,D), 485 (A,B,C,D), 494 (A,D), 495 (A,B,C,D), 497 (A,D), 498 (A,B,C,D), 501 (A,B,C,D), 502 (A,B,C,D), 503 (A,B,C,D), 508 (A,B,C,D), 509 (A,B,C,D), 510 (A,B,C,D), 511 (A,B,C,D), 512 (A,B,C), 513 (A,B,C), 514 (C), 515 (A,B,C), 516 (A,D), 517 (A,D), 519 (B,C,D), 520 (B,D), 521 (B,C,D), 522 (B,D), 523 (A,B,D), 524 (A,B,D), 525 (A,B,C,D), 526 (A,B,C,D), 573 (A,B,C,D), 604 (A,B,C,D), 640 (A), 665 (A,B,C,D), 666 (B), 667 (B), 671 (B), 672 (B), 675 (C), 681 (A,B,C,D), 682 (A,B,C,D), 687 (A,B,C,D), 688 (A,B,C), 689 (C,D), 690 (D), 691 (B,D), 695 (B), 696 (B,D), 697 (B,D), 706 (D), 707 (D), 723 (A), 724 (A), 727 (A), 728 (A), 729 (A,B,C), 735 (A,B,C), 738 (A,B,C), 740 (A,B,C), 741 (A,B,C), 743 (A,B,C,D), 744 (A,B,C,D), 746 (A,B,C,D), 747 (A,B,C,D), 751 (A,B,C,D), 752 (A,B,C,D), 753 (A,B,C,D), 755 (B,C), 756 (A,B,C,D), 757 (A,B,C), 758 (B,C), 759 (A,C,D), 760 (A,B,C,D), 761 (A,B,C), 762 (A,B,C,D), 763 (B,C,D), 764 (A,B,C), 765 (A,B,C,D), 766 (B,C,D), 767 (B,C,D), 768 (A,B,C,D), 770 (B,C,D), 771 (A,B,C,D), 773 (B,C,D), 774 (B,C,D), 775 (A), 776 (B,C,D), 778 (A,D), 779 (A,B,C,D), 780 (A,B,C,D), 781 (A,B,C,D), 782 (C), 783 (B,C,D), 784 (A,B,C,D), 785 (C), 786 (A,B,C,D), 787 (B,D), 788 (C), 789 (A,B,C,D), 790 (B), 793 (A,B,C,D), 794 (A,B,D), 796 (A,B,C,D), 797 (A,B,D) |
| **Anc165 – Total residues: 393** | |
| **Domain** | **Residues** |
| **CTD and EPDB domains** | 5 (A), 6 (A), 7 (A), 8 (A,C), 9 (A,C), 10 (A,C), 11 (A,C), 12 (C), 13 (C), 14 (A), 16 (C), 17 (A), 19 (C), 20 (A), 22 (C), 24 (A), 25 (A), 26 (C), 27 (C), 28 (A), 30 (C), 31 (A), 33 (C), 88 (A), 89 (A,B), 90 (B), 91 (A,D), 92 (A,B,D), 93 (B), 94 (D), 95 (D), 98 (A,B), 99 (A,B,C), 100 (B,C), 101 (B,C,D), 102 (A,C), 103 (C), 104 (C), 110 (A), 111 (A,B), 112 (B,C), 113 (A,C,D), 114 (A,B,D), 115 (B), 116 (C,D), 117 (A,D), 118 (B), 119 (C), 120 (A,D), 121 (B), 122 (C), 123 (D), 135 (A), 136 (A,B), 137 (A,B,C), 138 (A,B,C,D), 139 (A,B,C,D), 140 (A,B,C,D), 141 (B,C,D), 142 (C,D), 143 (C), 145 (C), 146 (C), 147 (C), 148 (C), 149 (C), 151 (C), 152 (C), 153 (A,C), 154 (B,C), 155 (A,B,C), 156 (A,B,C,D), 157 (A,C,D), 158 (B,D), 159 (A,B,C), 160 (A,C,D), 161 (B,D), 162 (A,B,C), 163 (A,D), 164 (B,D), 165 (A,B,C), 166 (A,B,D), 167 (B,C,D), 168 (C,D), 169 (B,D), 171 (D), 176 (B), 177 (B), 178 (B,D), 179 (C,D), 180 (D), 211 (B), 212 (B), 213 (D), 214 (D), 215 (B), 216 (B), 217 (D), 218 (D), 219 (B), 220 (B), 221 (D), 222 (D), 223 (B), 225 (D), 242 (B), 243 (B,C), 244 (B,D), 245 (D), 246 (D), 247 (A), 248 (B), 249 (B,C), 250 (A,D), 251 (B), 252 (B,C), 253 (A,D), 254 (B), 255 (B,C), 256 (A,D), 257 (B), 258 (B,C), 259 (A,D), 260 (A,B), 261 (A,B,C), 262 (B,C,D), 263 (A,C,D), 264 (B,D), 265 (C), 266 (A,D), 267 (B), 268 (C), 269 (A,D), 270 (A,B), 271 (A,B,C), 272 (A,C,D), 273 (A,B,D), 274 (A,B,C) |
| **GT-B domain** | 275 (A,B,C,D), 276 (A,B,C,D), 277 (B,C,D), 278 (C,D), 279 (B,C,D), 280 (A,B,C), 281 (A,B,C), 282 (A,B,C), 283 (B,C,D), 284 (C,D), 285 (C,D), 286 (C), 287 (C), 288 (C), 296 (C), 301 (C), 311 (A), 313 (A), 314 (C,D), 316 (D), 317 (C,D), 320 (C,D), 323 (C,D), 325 (B), 326 (B,C,D), 327 (B), 328 (C,D), 329 (B,D), 330 (B,C), 331 (B,C,D), 332 (B,C,D), 333 (B,C,D), 334 (A,C,D), 335 (A,C,D), 336 (C), 337 (C), 338 (C,D), 354 (A), 357 (A,D), 360 (D), 361 (A), 362 (A), 364 (D), 365 (D), 366 (D), 382 (C), 384 (A), 385 (A,B,C), 386 (A,C), 387 (A,D), 388 (B,D), 389 (C,D), 390 (A,D), 391 (A,B), 392 (B,C), 393 (C,D), 394 (D), 410 (A,D), 411 (A,D), 412 (D), 413 (C,D), 414 (D), 415 (B,C), 416 (A,B), 417 (B), 418 (C), 419 (A,D), 420 (A,B), 421 (C), 422 (C,D), 423 (D), 432 (A), 433 (A,B), 435 (C), 437 (C), 438 (C), 449 (C), 452 (C), 453 (C), 456 (C), 460 (C), 461 (A), 462 (B), 463 (B,C), 464 (A,D), 465 (B), 466 (B,C), 467 (A,B,C,D), 468 (A,D), 469 (B), 470 (B,C), 471 (A,C,D), 472 (A,B), 473 (A,B), 474 (B,C,D), 475 (A,C,D), 476 (A,B,D), 477 (A,B,C), 478 (A,B,C,D), 479 (D), 480 (A,D), 481 (A,B,D), 482 (A,B,C,D), 483 (B,C,D), 484 (A,C,D), 485 (A,D), 486 (B), 488 (D), 492 (C), 493 (B,C), 494 (C), 495 (A), 496 (B), 497 (A,C), 498 (A,D), 499 (A,B), 500 (C), 501 (A,C,D), 502 (A,B,C,D), 503 (A,B,C), 504 (B,C,D), 505 (C), 506 (D), 508 (A), 509 (A,B), 510 (A,B,C), 511 (A,B,D), 512 (A,B,C,D), 513 (A,B,C,D), 514 (B,C,D), 515 (A,C,D), 516 (B,D), 517 (C,D), 518 (D), 519 (A), 520 (A,B), 521 (A,B,C), 522 (A,B,C,D), 523 (A,B,C,D), 524 (A,B,C,D), 525 (A,B,C,D), 526 (A,B,C,D), 527 (B,C,D), 528 (C,D), 529 (D), 573 (A,C), 574 (B,C), 596 (B), 598 (D), 599 (B,D), 601 (D), 604 (A), 605 (B,C), 647 (C), 648 (C), 649 (C), 662 (C), 664 (A), 665 (A), 666 (B,C), 667 (A,C), 668 (B,D), 672 (C), 674 (C), 675 (C), 676 (C), 681 (A), 682 (A,B), 683 (B,C), 684 (C,D), 687 (A), 688 (A,B), 689 (B,C), 690 (C), 691 (A,D), 692 (A,B), 693 (C), 694 (A,C), 695 (A), 696 (A,B,C), 697 (A,B,C), 698 (B), 701 (A), 724 (A,B), 725 (A,B), 726 (B,D), 727 (D), 728 (A,D), 729 (A,B), 730 (B,D), 731 (D), 732 (D), 736 (A), 737 (A,B), 738 (B), 739 (D), 740 (A,B,D), 741 (A,B,D), 742 (B,C,D), 743 (C,D), 744 (A,D), 745 (B), 746 (C), 747 (A,D), 748 (B), 749 (C), 750 (D), 751 (A), 752 (B), 753 (A,C), 754 (B,D), 755 (C,D), 756 (A,D), 757 (A,B), 758 (C,D), 759 (A,D), 760 (B,D), 761 (C,D), 762 (A,D), 763 (B), 764 (C,D), 765 (A,D), 766 (A,B), 767 (A,C,D), 768 (A,D), 770 (A,D), 771 (A), 773 (D), 774 (B), 775 (B,C), 776 (A,B,D), 777 (A,B,C,D), 778 (A,B,C,D), 779 (A,B,C), 780 (A,B,D), 781 (A,B,C,D), 782 (B,C), 783 (A,D), 784 (A,B,D), 785 (B,C), 786 (A,C,D), 787 (B,D), 788 (C), 789 (A,C,D), 790 (B), 791 (C), 792 (C,D), 793 (A), 794 (A,B), 795 (B,C), 796 (A,C,D), 797 (A,B,D), 798 (B,C), 799 (C,D), 800 (D) |

Letters inside the parenthesis refer to the respective monomer of SuSy.

**Table S10.** Apparent melting temperatures for single amino acid oligomerization mutants on WT.

| **Mutation** | **Thermal transition 1** | | | | **Thermal transition 2** | | | |
| --- | --- | --- | --- | --- | --- | --- | --- | --- |
|  | **ΔTm_app_ (°C)** | **SD (°C)** | **ΔTm_app_ (°C)** | **SD (°C)** | **ΔTm_app_ (°C)** | **SD (°C)** | **ΔTm_app_ (°C)** | **SD (°C)** |
| F131G | 44.90 | 0.17 | -0.14 | 0.17 | 57.29 | 0.97 | -4.44 | 0.97 |
| E780G | 45.12 | 0.12 | 0.08 | 0.12 | 65.22 | 0.06 | 3.49 | 0.06 |
| R783G | ND | ND | ND | ND | 44.86 | 0.07 | -16.87 | 0.17 |
| E786G | 44.52 | 0.19 | -0.52 | 0.19 | 56.56 | 0.27 | -5.17 | 0.27 |
| M787G | ND | ND | ND | ND | 45.54 | 0.32 | -16.19 | 0.36 |
| F159G | ND | ND | ND | ND | 44.84 | 0.23 | -16.89 | 0.28 |
| R162G | ND | ND | ND | ND | 44.28 | 0.06 | -17.45 | 0.17 |
| A166G | ND | ND | ND | ND | 44.88 | 0.26 | -16.85 | 0.31 |
| H170G | 44.91 | 0.16 | -0.13 | 0.16 | 61.22 | 0.27 | -0.51 | 0.27 |
| R209G | 44.39 | 0.18 | -0.64 | 0.18 | 63.36 | 0.09 | 1.63 | 0.09 |
| E259G | ND | ND | ND | ND | 45.13 | 0.16 | -16.60 | 0.23 |
| E780G,R783G | ND | ND | ND | ND | 44.24 | 0.17 | -17.49 | 0.24 |
| E786G,M787G | ND | ND | ND | ND | 43.12 | 0.16 | - 18.61 | 0.22 |
| F159G,R162G | ND | ND | ND | ND | 44.38 | 0.11 | -17.35 | 0.20 |
| A166G,H170G | ND | ND | ND | ND | 43.42 | 0.10 | -18.31 | 0.19 |
| F131G,E780G,R783G | 44.20 | 0.17 | -0.84 | 0.17 | 58.07 | 0.63 | -3.66 | 0.63 |
| R209G,E259G | 44.08 | 0.25 | -0.95 | 0.25 | 62.22 | 0.62 | 0.49 | 0.62 |
| F131G,E780G,R783G,E786G, M787G | 44.32 | 0.34 | -0.72 | 0.34 | 62.60 | 0.10 | 0.87 | 0.10 |
| A166C | 45.32 | 0.09 | 0.33 | 0.09 | 62.53 | 0.07 | 0.91 | 0.07 |
| F169C | 44.93 | 0.07 | -0.06 | 0.08 | 61.58 | 0.04 | -0.05 | 0.04 |
| L255N | 44.81 | 0.13 | -0.18 | 0.13 | 61.76 | 0.10 | 0.13 | 0.10 |
| A166C,F169C | 44.87 | 0.10 | -0.12 | 0.10 | 62.43 | 0.05 | 0.81 | 0.05 |
| A166C,F169C,L255N | 45.30 | 0.07 | 0.32 | 0.07 | 61.33 | 0.09 | -0.29 | 0.09 |
| A142D | 44.96 | 0.11 | -0.02 | 0.11 | 59.89 | 0.07 | -1.74 | 0.08 |
| E786K | 44.97 | 0.16 | -0.02 | 0.16 | 56.33 | 0.04 | -5.29 | 0.04 |
| A790S | 44.90 | 0.05 | -0.09 | 0.04 | 60.04 | 0.14 | -1.59 | 0.14 |
| E786K,A790S,L791C | 44.98 | 0.12 | -0.09 | 0.12 | 63.47 | 0.44 | 1.84 | 0.44 |
| L791C | 46.53 | 0.63 | -0.52 | 0.66 | 63.59 | 0.23 | 1.13 | 0.23 |
| A424S | 44.58 | 0.49 | -0.41 | 0.49 | 62.09 | 0.01 | 0.47 | 0.02 |
| G765S | 45.10 | 0.28 | 0.11 | 0.28 | 61.84 | 0.06 | 0.22 | 0.06 |
| E786K,A790S,L791C,A142D | 44.86 | 0.04 | -0.13 | 0.04 | 58.88 | 1.78 | -2.75 | 1.78 |

SD refers to standard deviation of triplicates (*n*=3). ND stands for not detected.

**Table S11.** Relative activity for single amino acid oligomerization mutants on WT.

| **Mutation** | **Activity** | |
| --- | --- | --- |
|  | **Relative (%)** | **SD (%)** |
| F131G | 19.3 | 5.8 |
| E780G | 43.9 | 11.2 |
| R783G | 4.8 | 3.3 |
| E786G | 23.5 | 8.1 |
| M787G | 1.6 | 3.8 |
| F159G | 5.1 | 2.0 |
| R162G | 12.0 | 4.2 |
| A166G | 10.0 | 4.1 |
| H170G | 54.2 | 13.7 |
| R209G | 54.2 | 14.4 |
| E259G | 16.2 | 5.8 |
| E780G,R783G | 24.6 | 8.6 |
| E786G,M787G | 8.8 | 4.8 |
| F159G,R162G | 5.4 | 3.3 |
| A166G,H170G | 4.2 | 2.6 |
| F131G,E780G,R783G | 13.8 | 7.0 |
| R209G,E259G | 33.9 | 11.6 |
| F131G,E780G,R783G,E786G,M787G | 46.8 | 14.1 |
| A166C | 134.8 | 55.3 |
| F169C | 166.2 | 59.4 |
| L255N | 104.9 | 37.6 |
| A166C,F169C | 96.5 | 34.6 |
| A166C,F169C,L255N | 73.7 | 29.5 |
| A142D | 118.6 | 43.9 |
| E786K | 72.3 | 22.8 |
| A790S | 123.4 | 42.0 |
| E786K,A790S,L791C | 39.8 | 12.9 |
| L791C | 65.7 | 22.5 |
| A424S | 80.8 | 25.3 |
| G765S | 70.9 | 23.4 |
| E786K,A790S,L791C,A142D | 69.5 | 27.2 |

SD refers to standard deviation of triplicates (*n*=3).

**Table S12.** Inputs and outputs from mass balances for the production of 1 kg of MANT-*N*-glucose.

| **Flows** | **Provider in openLCA** | **Allocation** | **Unit** | **UDP-Glc** | **WT** | **HPMN251** |
| --- | --- | --- | --- | --- | --- | --- |
| **Inputs** | | | | | | |
| Anthranilic acid | Market for anthranilic acid | GLO | kg | 0.486 | 1.018 | 0.486 |
| Acid catalyst | Market for cationic resin | RER | kg | 0.131 | 0.275 | 0.131 |
| Methanol | Market for methanol | GLO | kg | 3.852 | 8.064 | 3.852 |
| NH_4_Cl | Market for ammonium chloride | GLO | kg | 0.003 | - | - |
| Glucose | Market for glucose | GLO | kg | 19.834 | 0.379 | 0.262 |
| MgSO_4_ | Market for magnesium sulphate | GLO | kg | 0.282 | 0.005 | 0.004 |
| Yeast extract / Tryptone | Market for protein feed, 100 % crude | GLO | kg | 0.467 | 0.007 | 0.005 |
| NaCl | Market for sodium chloride, powder | GLO | kg | 0.042 | - | - |
| Buffer | Market for sodium phosphate | RER | kg | 11.478 | 4.726 | 2.004 |
| Sucrose | Market for sugar, from sugar beet | GLO | kg | 16.289 | 25.408 | 18.209 |
| Water | Market for water, deionized | EWS | kg | 352.192 | 337.037 | 161.666 |
| Brewer’s yeast | Market for fodder yeast | GLO | kg | 39.220 | 0.753 | 0.520 |
| FeSO_4_ | Market for iron sulphate | GLO | g | 0.268 | 0.005 | 0.004 |
| MnSO_4_ | Market for manganese sulphate | GLO | g | 0.436 | 0.008 | 0.006 |
| Enzyme | Market for enzymes | GLO | kg | 0.021 | 0.046 | 0.016 |
| DMSO | Market for dimethyl sulfoxide | GLO | kg | 19.506 | 40.826 | 19.506 |
| **Outputs** | | | | | | |
| Hazardous waste | Waste incineration with energy recovery | EWS | kg | 3.934 | 8.234 | 3.934 |
| Biowaste | Municipal incineration | GLO | kg | 42.363 | 0.768 | 0.530 |
| Wastewater | Wastewater, average, capacity 10^9^ L | EWS | L | 416.616 | 408.539 | 201.194 |
| MANT-*N*-glucose |  |  | kg | 1.000 | 1.000 | 1.000 |

EWS, Europe without Switzerland; RER, region of Europe; GLO, global. Energy inputs, infrastructure, solvent recovery, and downstream purification of intermediates and products were excluded from the system boundary, consistent with the early-stage nature of the process. Mass balances were based on Gharabli et al., (2025)^3^ considering MANT^4^, UDP^5,6^, and UDP-Glc synthesis^7^.

**Table S13.** Mass balance and costs for production of 1 kg of MANT-*N*-glucose.

| **Input** | **UDP-Glc** | | **WT** | | **HPMN251** | |
| --- | --- | --- | --- | --- | --- | --- |
|  | Mass (kg) | Cost (USD) | Mass (kg) | Cost (USD) | Mass (kg) | Cost (USD) |
| MANT | 0.536 | 5.361 | 1.122 | 11.220 | 0.536 | 5.361 |
| UDP-Glc | 3.246 | 275.585 | - | - | - | - |
| Buffer | 1.237 | 0.940 | 4.532 | 3.444 | 1.869 | 1.420 |
| Enzymes | 0.021 | 0.525 | 0.046 | 1.146 | 0.016 | 0.412 |
| DMSO | 19.506 | 29.258 | 40.826 | 61.239 | 19.506 | 29.258 |
| Water | 159.591 | 1.229 | 334.029 | 2.572 | 159.591 | 1.229 |
| Sucrose | - | - | 25.408 | 12.704 | 18.209 | 9.105 |
| UDP | - | - | 0.058 | 2.919 | 0.040 | 1.987 |
| Total | 184.138 | 312.899 | 406.021 | 95.244 | 199.768 | 48.772 |

**Table S14.** Results of calculation of endpoint impact using ReCiPe 2106 (H) for the production of 1 kg of MANT-*N*-glucose.

| **Impact category** | **Unit** | **UDP-Glc** | **WT** | **HPMN251** |
| --- | --- | --- | --- | --- |
| Land use | species·yr | 9.760E-07 | 2.018E-07 | 1.370E-07 |
| Global warming, Human health | DALY | 2.365E-04 | 1.158E-04 | 5.909E-05 |
| Marine ecotoxicity | species·yr | 1.606E-09 | 8.688E-10 | 4.199E-10 |
| Fine particulate matter formation | DALY | 2.796E-04 | 1.609E-04 | 8.446E-05 |
| Water consumption, Terrestrial ecosystem | species·yr | 6.126E-08 | 4.395E-08 | 2.492E-08 |
| Freshwater ecotoxicity | species·yr | 8.074E-09 | 4.306E-09 | 2.080E-09 |
| Terrestrial ecotoxicity | species·yr | 1.334E-08 | 6.780E-09 | 3.317E-09 |
| Water consumption, Human health | DALY | 9.551E-06 | 7.217E-06 | 4.091E-06 |
| Mineral resource scarcity | USD2013 | 3.158E-01 | 1.547E-01 | 7.342E-02 |
| Ozone formation, Terrestrial ecosystems | species·yr | 6.068E-08 | 3.480E-08 | 1.841E-08 |
| Global warming, Freshwater ecosystems | species·yr | 1.948E-11 | 9.545E-12 | 4.869E-12 |
| Human carcinogenic toxicity | DALY | 7.291E-05 | 4.139E-05 | 1.947E-05 |
| Fossil resource scarcity | USD2013 | 1.807E+01 | 1.973E+01 | 9.716E+00 |
| Freshwater eutrophication | species·yr | 4.873E-08 | 2.426E-08 | 1.198E-08 |
| Human non-carcinogenic toxicity | DALY | 5.080E-05 | 2.855E-05 | 1.376E-05 |
| Ozone formation, Human health | DALY | 4.164E-07 | 2.378E-07 | 1.260E-07 |
| Water consumption, Aquatic ecosystems | species·yr | 6.951E-12 | 2.047E-12 | 1.171E-12 |
| Marine eutrophication | species·yr | 3.279E-10 | 8.301E-11 | 5.515E-11 |
| Ionizing radiation | DALY | 6.560E-08 | 4.318E-08 | 2.190E-08 |
| Global warming, Terrestrial ecosystems | species·yr | 7.132E-07 | 3.495E-07 | 1.783E-07 |
| Terrestrial acidification | species·yr | 3.286E-07 | 1.863E-07 | 1.039E-07 |
| Stratospheric ozone depletion | DALY | 5.491E-07 | 1.512E-07 | 9.445E-08 |
| **Normalized impacts** | Pt | 1.54526E8 | 1.67133E8 | 8.22648E7 |

EWS, Europe without Switzerland; RER, region of Europe; GLO, global. Normalization of endpoint categories was done using World (2010) H/H as weighting set. The normalization allows direct comparison by assigning a single score (Pt) to each system.

**Table S15.** Results of central composite design on indoxyl glycosylation.

| **Treatment** | **pH** | **UDP** | **Sucrose** | **Temperature** | **Conversion (%)** | **SD** |
| --- | --- | --- | --- | --- | --- | --- |
| 1 | -1 | -1 | -1 | -1 | 17.33 | 4.78 |
| 2 | 1 | -1 | -1 | -1 | 21.75 | 0.71 |
| 3 | -1 | -1 | -1 | 1 | 15.63 | 1.56 |
| 4 | 1 | -1 | -1 | 1 | 20.25 | 1.34 |
| 5 | -1 | -1 | 1 | -1 | 16.42 | 2.82 |
| 6 | 1 | -1 | 1 | -1 | 33.75 | 3.98 |
| 7 | -1 | -1 | 1 | 1 | 18.32 | 2.35 |
| 8 | 1 | -1 | 1 | 1 | 24.78 | 3.15 |
| 9 | -1 | 1 | -1 | -1 | 23.11 | 3.36 |
| 10 | 1 | 1 | -1 | -1 | 27.24 | 3.76 |
| 11 | -1 | 1 | -1 | 1 | 21.26 | 1.34 |
| 12 | 1 | 1 | -1 | 1 | 26.03 | 0.67 |
| 13 | -1 | 1 | 1 | -1 | 32.45 | 6.47 |
| 14 | 1 | 1 | 1 | -1 | 32.72 | 1.83 |
| 15 | -1 | 1 | 1 | 1 | 29.09 | 4.45 |
| 16 | 1 | 1 | 1 | 1 | 45.48 | 1.62 |
| 17 | -2 | 0 | 0 | 0 | 12.85 | 3.04 |
| 18 | 2 | 0 | 0 | 0 | 35.07 | 9.47 |
| 19 | 0 | 0 | 0 | -2 | 19.12 | 3.43 |
| 20 | 0 | 0 | 0 | 2 | 11.10 | 0.28 |
| 21 | 0 | 0 | -2 | 0 | 18.36 | 3.8 |
| 22 | 0 | 0 | 2 | 0 | 27.93 | 4.43 |
| 23 | 0 | -2 | 0 | 0 | 15.00 | 2.21 |
| 24 | 0 | 2 | 0 | 0 | 22.27 | 0.74 |
| 25 | 0 | 0 | 0 | 0 | 21.16 | 2.72 |
| 26 | 0 | 0 | 0 | 0 | 25.81 | 2.92 |
| 27 | 0 | 0 | 0 | 0 | 25.80 | 1.47 |
| 28 | 0 | 0 | 0 | 0 | 26.79 | 3.92 |
| 29 | 0 | 0 | 0 | 0 | 27.10 | 2.69 |
| 30 | 2 | 1 | 0 | -2 | 37.63 | 1.47 |

UGT/SuSy mass ratio was fixed at 1. pH (-2 = pH of 5.0; -1 = pH of 5.75; 0 = pH of 6.5; 1 = pH of 7.25; 2 = pH of 8.0). UDP (-2 = 0.001 mM; -1 = 0.01 mM; 0 = 0.1 mM; 1 = 1 mM; 2 = 2 mM). Sucrose (-2 = 50 mM; -1 = 125 mM; 0 = 200 mM; 1 = 400 mM; 2 = 600 mM). Temperature (-2 = 30 °C; -1 = 37.5 °C; 0 = 45 °C; 1 = 52.5 °C; 2 = 60 °C). SD refers to standard deviation of triplicates (*n*=3). Predicted conditions to maximize conversion were UDP: 2mM, Sucrose: 600 mM, pH: 8, and temperature: 54 °C.

**Table S16.** Inputs and outputs from mass balances for the production of 1 kg of indican.

| **Flows** | **Provider in openLCA** | **Allocation** | **Unit** | **WT** | **Literature** | **HPMN251** |
| --- | --- | --- | --- | --- | --- | --- |
| **Inputs** | | | | | | |
| Phthalimide | Market for phthalimide | GLO | kg | 1.700 | 2.074 | 0.616 |
| Glucose | Market for glucose | GLO | kg | 0.281 | 0.347 | 0.209 |
| MgSO_4_ | Market for magnesium sulphate | GLO | kg | 0.004 | 0.005 | 0.003 |
| Yeast extract / Tryptone | Market for protein feed, 100 % crude | GLO | kg | 0.005 | 0.007 | 0.004 |
| Buffer | Market for sodium phosphate | RER | kg | 1.326 | 1.620 | 0.536 |
|  | Market for citric acid | GLO | kg | 0.540 | 0.659 | 0.195 |
| Sucrose | Market for sugar, from sugar beet | GLO | kg | 6.643 | 8.106 | 7.219 |
| Water | Market for water, deionized | EWS | kg | 99.267 | 121.158 | 36.808 |
| Brewer’s yeast | Market for fodder yeast | GLO | kg | 0.559 | 0.688 | 0.416 |
| FeSO_4_ | Market for iron sulphate | GLO | g | 0.004 | 0.005 | 0.003 |
| MnSO_4_ | Market for manganese sulphate | GLO | g | 0.006 | 0.008 | 0.005 |
| Enzyme | Market for enzymes | GLO | kg | 0.137 | 0.364 | 0.050 |
| **Outputs** | | | | | | |
| Biowaste | Municipal incineration | GLO | kg | 0.570 | 0.702 | 0.424 |
| Wastewater | Wastewater, average, capacity 10^9^ L | EWS | L | 108.890 | 133.326 | 44.631 |
| Indican |  |  | kg | 1.000 | 1.000 | 1.000 |

EWS, Europe without Switzerland; RER, region of Europe; GLO, global. Energy inputs, infrastructure, solvent recovery, and downstream purification of intermediates and products were excluded from the system boundary, consistent with the early-stage nature of the process. Mass balances were carried out considering UDP^5,6^ synthesis. Because the environmental impacts of producing indoxyl acetate are not yet established, phthalimide was selected as a proxy due to its chemical similarity and its classification as a fine chemical.

**Table S17.** Mass balance and costs for production of 1 kg of indican.

| **Input** | **WT** | | **Literature** | | **HPMN251** | |
| --- | --- | --- | --- | --- | --- | --- |
|  | Mass (kg) | Cost (USD) | Mass (kg) | Cost (USD) | Mass (kg) | Cost (USD) |
| Indoxyl acetate | 1.700 | 8.499 | 2.074 | 10.371 | 0.616 | 3.079 |
| Sucrose | 6.643 | 3.321 | 8.106 | 4.053 | 7.219 | 3.609 |
| UDP | 0.043 | 2.174 | 0.053 | 2.653 | 0.032 | 1.575 |
| Buffer | 1.720 | 0.172 | 2.100 | 0.210 | 0.623 | 0.062 |
| Enzymes | 0.137 | 3.437 | 0.364 | 9.107 | 0.050 | 1.245 |
| Water | 97.034 | 0.747 | 118.409 | 0.912 | 35.148 | 0.271 |
| Total | 107.278 | 18.351 | 131.106 | 27.306 | 43.687 | 9.841 |

**Table S18.** Results of calculation of endpoint impact using ReCiPe 2106 (H) for the production of 1 kg of indican.

| **Impact category** | **Unit** | **WT** | **Literature** | **HPMN251** |
| --- | --- | --- | --- | --- |
| Land use | species·yr | 7.167E-08 | 7.625E-08 | 6.109E-08 |
| Global warming, Human health | DALY | 2.079E-05 | 2.172E-05 | 1.159E-05 |
| Marine ecotoxicity | species·yr | 1.795E-10 | 1.940E-10 | 8.749E-11 |
| Fine particulate matter formation | DALY | 3.101E-05 | 3.333E-05 | 1.908E-05 |
| Water consumption, Terrestrial ecosystem | species·yr | 1.233E-08 | 1.379E-08 | 8.181E-09 |
| Freshwater ecotoxicity | species·yr | 9.120E-10 | 9.839E-10 | 4.418E-10 |
| Terrestrial ecotoxicity | species·yr | 1.555E-09 | 1.672E-09 | 7.739E-10 |
| Water consumption, Human health | DALY | 2.020E-06 | 2.264E-06 | 1.340E-06 |
| Mineral resource scarcity | USD2013 | 3.510E-02 | 3.964E-02 | 1.690E-02 |
| Ozone formation, Terrestrial ecosystems | species·yr | 7.468E-09 | 7.969E-09 | 4.428E-09 |
| Global warming, Freshwater ecosystems | species·yr | 1.714E-12 | 1.790E-12 | 9.548E-13 |
| Human carcinogenic toxicity | DALY | 7.977E-06 | 9.188E-06 | 3.722E-06 |
| Fossil resource scarcity | USD2013 | 2.126E+00 | 2.406E+00 | 1.058E+00 |
| Freshwater eutrophication | species·yr | 5.076E-09 | 5.466E-09 | 2.586E-09 |
| Human non-carcinogenic toxicity | DALY | 6.450E-06 | 6.998E-06 | 3.072E-06 |
| Ozone formation, Human health | DALY | 5.083E-08 | 5.409E-08 | 3.036E-08 |
| Water consumption, Aquatic ecosystems | species·yr | 6.117E-13 | 6.492E-13 | 4.107E-13 |
| Marine eutrophication | species·yr | 2.752E-11 | 2.965E-11 | 2.304E-11 |
| Ionizing radiation | DALY | 7.333E-09 | 7.918E-09 | 4.014E-09 |
| Global warming, Terrestrial ecosystems | species·yr | 6.274E-08 | 6.553E-08 | 3.495E-08 |
| Terrestrial acidification | species·yr | 3.923E-08 | 4.319E-08 | 2.794E-08 |
| Stratospheric ozone depletion | DALY | 3.776E-08 | 3.887E-08 | 3.257E-08 |
| Normalized impacts | Pt | 1.81575E7 | 2.05475E7 | 9.03406E6 |

EWS, Europe without Switzerland; RER, region of Europe; GLO, global. Normalization of endpoint categories was done using World (2010) H/H as weighting set. The normalization allows direct comparison by assigning a single score (Pt) to each system.

**Table S19**. List of primers used to generate the mutations evaluated in this study.

| **Mutation position** | | **Primer forward (5’ – 3’)** | **Primer reverse (5’ – 3’)** |
| --- | --- | --- | --- |
| **Original** | **pET28a+** |  |  |
| N23Y | N44Y | ACTGCCTAUAGGAATGAAATTTTGGCCCTT | ATAGGCAGUGAGGGTTTCATCAAGCCTCTCA |
| A66P | A87P | ATGGTCCGUTTGGAGAAGTCTTGAGATCTAC | ACGGACCAUCAGTGAGCTTCTGTCTGTTC |
| L86F | L107F | ATGGGTTGCUTTTGCTGTTCGTCCAAGACCT | AGCAACCCAUGGTGGCAAAACTATGGCTTCCT |
| Q111P | Q132P | AGTTGCCGCCUGCTGAGTACCTGCACTTC | AGGCGGCAACUCCTCAACAACAAGAGCGTGCACAT |
| V132M | V153M | ACTTTATGCUTGAGTTGGACTTTGAACCATTC | AGCATAAAGUTGCCATTAGAACTTCCGTCAA |
| S152Y | S173Y | AAGTATATUGGAAATGGTGTGCAATTCCT | AATATACTUGTTAAGAGTTGGGCGGGGG |
| V218L | V239L | ACACTGCCUCCTGAAACTCCCTACTCAGA | AGGCAGTGUGCCCAGATACTCCTCAGCTT |
| H229R | H250R | AGCGTAAGUTCCAGGAGATTGGTTTGGAG | ACTTACGCUCAAATTCTGAGTAGGGAGTTTCA |
| V410I | V431I | ATTGGAAACUACAGTGATGGAAACATTGTCG | AGTTTCCAAUAATCAGATCTGGCTTGCCTTGCA |
| Q528R | Q549R | ATCGTACCAUTTACTTCCCCCACACTGAAAC | ATGGTACGAUCAGCTCCAGGGGAGACAA |
| H534Y | H555Y | ACTTCCCCUATACTGAAACCAGCCGTAGG | AGGGGAAGUAAATGGTTTGATCAGCTCCAG |
| C563L | C584L | ATTCTGGUGCTGAAGGACCGCAGCAA | ACCAGAAUGTGTTCTTCATTCTCCACTGAG |
| L687C | L708C | ACTTGCGGCUGCCCAACATTCGCCACATGC | AGCCGCAAGUCATGGCCTCAACCACTGTCA |
| I755Y | I776Y | AATATTACUCTCAGAGGCTTCTCACTC | AGTAATATUGCCATGTGTACTTCTCTTCAATA |
| L796C | L817C | AAATGTGCUGAGTCTGTGCCCCTTGCTGC | AGCACATTUGCGGTACTTGAGAGCATAG |
| V47I | V68I | AGATTAUTGCTGAGTTTGAGGAAATCC | ATAATCUGGTGGTGTTGCAGGATGCC |
| A104E | A125E | ACGAACUTGTTGTTGAGGAGTTGCAAC | AGTTCGUGCACATTCACTCTCAGGTACT |
| E180D | E201D | ATTTCCUCAGGCTTCACAGCGTCAA | AGGAAAUCCAAAAGTGGGTGCAAGCTCTCCTT |
| G216R | G237R | ATCTGCGUACAGTGCCTCCTGAAACTC | ACGCAGAUACTCCTCAGCTTTCCTCAGA |
| N315M | N336M | AGATGGAGAUGCTCCATCGCATTAAGCA | ATCTCCATCUCCAAAGCACGAACTTGATCCA |
| A395M | A416M | ATGTTAUGCACGAGCTTGCCAAAGAGT | ATAACAUCCTCAGTGTAAGTTTCCAAGTAGG |
| L453F | L474F | AAAATTUGAAGAGAGATACCACTTCTCT | AAATTTUTTCCAGTAAATGTCGGATTCGG |
| A465V | A486V | ACAGTGGAUCTATTTGCCATGAACCACACA | ATCCACTGUGAATTGGCAAGAGAAGTGGTATC |
| F481Y | F502Y | AGTACCTAUCAGGAGATTGCTGGAAGC | ATAGGTACUGGTGATAATGAAATCTGTGTGGTT |
| V514P | V535P | ATCCGTUTGATCCAAAATTCAACATTGTCTC | AACGGAUCAATACCATGCACAACGCGGTAGA |
| L618R | L639R | AGGACCGUGAAGAAAAGGCCGAGATGA | ACGGTCCUTTGACTCCTTCCTCCTGTCT |
| G629L | G650L | ACCTGCUGATCGAGACCTACAAGTTGAA | AGCAGGUACATCTTCTTCATCTCGGCCTTT |
| V671F | V692F | AGCCTGCTUTTTACGAGGCTTTTGGTTTGAC | AAGCAGGCUGCACGAAAGCACCCCTGG |
| N694Y | N715Y | ATGCTATGGUGGTCCTGCTGAGATCATTG | ACCATAGCAUGTGGCGAATGTTGGCAAGC |
| N775K | N796K | ATGTGTCUAAACTTGACCGCCGTGAGAGCCG | AGACACAUGCTTCCAGAAGCCATAGACA |
| V132E | V153E | AGTTCAAAGUTGCCATTAGAACTTCCGTCAA | ACTTTGAACUTGAGTTGGACTTTGAACCATTC |
| S73Y | S94Y | ATCTCAAGACUTCTCCAAAGGCACCATCAG | AGTCTTGAGAUATACACAGGAAGCCATAGTTTTG |
| V132K | V153K | AGTTTAAAGUTGCCATTAGAACTTCCGTCAA | ACTTTAAACUTGAGTTGGACTTTGAACCATTC |
| A143S | A164S | AGCTTGCAUTGAATGGTTCAAAGTCCAACTCA | ATGCAAGCUTCCCCCGCCCAACTCTTA |
| T222L | T243L | AGGGCAGTUCAGGAGGCACTGTGCCCAGAT | AACTGCCCUACTCAGAATTTGAGCACAAGT |
| S250M | S271M | ATCATCUCAAGGACACGCTCCGCGTTGT | AGATGAUTCAACTTCTCTTGGATCTTCT |
| M317L | M338L | AGCAGCUCATTCTCCAAAGCACGAACT | AGCTGCUCCATCGCATTAAGCAACAA |
| V329T | V350T | ATTGGACAUTACCCCTCGTATTCTC | ATGTCCAAUCCTTGTTGCTTAATGC |
| G348N | G369N | AACTACTUGTAACCAACGTCTTGAGAAGGTG | AAGTAGTUCCTACTGCATCGGGGAG |
| V374L | V395L | AATTCCCUTCTCAGTTCTAAAGGGAACTCGAA | AGGGAATUCTGCGCAAGTGGATCTCAAGA |
| V374I | V395I | AATAATUCCCTTCTCAGTTCTAAAGGGAACT | AATTATUCGCAAGTGGATCTCAAGA |
| Y390F | Y411F | AGTAAAAGUTTCCAAGTAGGGCCAGACTT | ACTTTTACUGAGGATGTTGCCCACGAG |
| A395V | A416V | ACAACAUCCTCAGTGTAAGTTTCCAAGTAG | ATGTTGUCCACGAGCTTGCCAAAGAGTTGCA |
| R456K | R477K | ATTTCTCUTCCAATTTTTTCCAGTAAATGTCG | AGAGAAAUACCACTTCTCTTGCCAATTC |
| T490W | T511W | ACCCAGUCCTTGCTTCCAGCAATCTC | ACTGGGUTGGACAGTACGAATCTCACA |
| A622M | A643M | ATCTCCATCUTTTCTTCCAAGTCCTTTGACTCC | AGATGGAGAUGAAGAAGATGTACGGCCTG |
| M624R | M645R | ATCTTCTUACGCTCGGCCTTTTCTTCCAAGT | AAGAAGAUGTACGGCCTGATCGAGAC |
| V650L | V671L | AGACGGUTCATCTGCGATGAAATCCATCT | ACCGTCUGAGGAATGGAGAGCTCTACC |
| C660A | C681A | ATCACGCGGUAGAGCTCTCCATTCCTCACA | ACCGCGTGAUCGCGGACACCAGGGGTGCTTTC |
| R663K | R684K | ACCTTTGGUGTCGCAGATCACGCGGTAGA | ACCAAAGGUGCTTTCGTGCAGCCTGCTGTA |
| N694H | N715H | ACCATGGCAUGTGGCGAATGTTGGCAAGC | ATGCCATGGUGGTCCTGCTGAGATCATTG |
| Q745M | Q766M | ATACGCAUGAGACCAGCCTTTGAGATCT | ATGCGTAUTGAAGAGAAGTACACATGGC |
| Q754K | Q775K | ATTTTCCAUGTGTACTTCTCTTCAATACGCT | ATGGAAAAUTTACTCTCAGAGGCTTCTCA |
| H206S | H227S | ACGCTTUGGAGTGCATCTGGGTTTTGA | AAAGCGUTCTGAGGAAAGCTGAGGA |
| G216L | G237L | AGCAGAUACTCCTCAGCTTTCCTCAGA | ATCTGCUGACAGTGCCTCCTGAAACTC |
| Q403P | Q424P | AACTCTUTGGCAAGCTCGTGGGCAACA | AAGAGTUGCCGGGCAAGCCAGATCTGATT |
| S523P | S544P | AGGCGGGACAAUGTTGAATTTTGGATCAAAGACATC | ATTGTCCCGCCUGGAGCTGATCAAACCATTTACT |
| Q758E | Q779E | AGCCTTUCAGAGTAAATTTGCCATGTGTACTT | AAAGGCUTCTCACTCTCACCGGTGT |
| E18D | E39D | AGGGTATCAUCAAGCCTCTCACGGAGACT | ATGATACCCUCACTGCCAACAGGAATGAA |
| T21S | T42S | AGGGTTUCATCAAGCCTCTCACGGAGACT | AAACCCUCAGCGCCAACAGGAATGAAATTTTGG |
| E35V | E56V | ACGATCCUTGACAGAAGGGCCAAAATTTCA | AGGATCGUGGCCAAGGGCAAGGGCATCCT |
| H319L | H340L | AGGAGCAUCTCATTCTCCAAAGCACGAA | ATGCTCCUGCGCATTAAGCAACAAGGAT |
| Q528M | Q549M | ATGGTCATAUCAGCTCCAGGGGAGACAA | ATATGACCAUTTACTTCCCCCACACTGAAAC |
| T529S | T550S | ATGCTTUGATCAGCTCCAGGGGAGACA | AAAGCAUTTACTTCCCCCACACTGAAAC |
| T537K | T558K | ACGGCTTTUTTCAGTGTGGGGGAAGTAAATG | AAAAGCCGUAGGTTGACATCCTTCCACCC |
| C563G | C584G | ACGCCAAUGTGTTCTTCATTCTCCACTGAG | ATTGGCGUGCTGAAGGACCGCAGCAA |
| V658Y | V679Y | ATAGCGGUAGAGCTCTCCATTCCTCACA | ACCGCTAUATCTGCGACACCAGGGGTGCTT |
| T734S | T755S | AGTGGCTUGGGTCAAGCTTGCACTTCT | AAGCCACUGGGACAAGATCTCAAAGGC |
| F131G | F152G | ACGGAGUGCTTGAGTTGGACTTTGAAC | ACTCCGUTGCCATTAGAACTTCCGTCAACAA |
| E780G | E801G | ACCGCCGUGGAAGCCGCCGCTATCTTGAG | ACGGCGGUCAAGGTTAGACACATGCTTCCAGA |
| R783G | R804G | AGCCGCGGAUATCTTGAGATGTTCTATGC | ATCCGCGGCUCTCACGGCGGTCAAGGTTAGAC |
| E786G | E807G | ATCTTGGAAUGTTCTATGCTCTCAAGTA | ATTCCAAGAUAGCGGCGGCTCTCACGGC |
| M787G | M808G | AGGGATUCTATGCTCTCAAGTACCGCAAA | AATCCCUCAAGATAGCGGCGGCTCTCA |
| F159G | F180G | AAGGACUCAACCGTCACCTTTCTGC | AGTCCTUGCACACCATTTCCAATTGACTTGT |
| R162G | R183G | ACGGACACCUTTCTGCCAAACTCTTCCA | AGGTGTCCGUTGAGGAATTGCACACCATTTCCAA |
| A166G | A187G | AAAACTCUTCCACGACAAGGAGAGCTT | AGAGTTTUCCAGAAAGGTGACGGTTGAGGAATTG |
| H170G | H191G | AACTCTUCGGAGACAAGGAGAGCTTGCAC | AAGAGTUTGGCAGAAAGGTGACGGTTGAG |
| R209G | R230G | AAAAGCUGAGGAGTATCTGGGCACAGT | AGCTTTUCCCAGAACATGTTGGAGTGCATCTGG |
| E259G | E280G | AGCCCCUGACCCGTGCACCCTTGAG | AGGGGCUCCAAGAAGATCCAAGAGAAGTTGAAT |
| A166C | A187C | ACCTTTCTUGCAAACTCTTCCACGACAAGGA | AAGAAAGGUGACGGTTGAGGAATTGCACACCAT |
| F169C | F190C | AAACTCUGCCACGACAAGGAGAGCTTG | AGAGTTUGGCAGAAAGGTGACGGTTGAGGAA |
| L255N | L276N | AACGATCUTCTTGAGGCCCCTGACCC | AGATCGTUGAGAAGTTGAATTGACTCAAGGAC |
| A142D | A163D | ATGATGCCUTCCCCCGCCCAACTCTTA | AGGCATCAUTGAATGGTTCAAAGTCCAACTCAA |
| E786K | E807K | AAAATGUTCTATGCTCTCAAGTACC | ACATTTUAAGATAGCGGCGGCTCTCA |
| A790S | A811S | ATAGCCUCAAGTACCGCAAATTGGC | AGGCTAUAGAACATCTCAAGATAGCGGCG |
| L791C | L812C | ATGCTTGCAAGUACCGCAAATTGGCTGAGT | ACTTGCAAGCAUAGAACATCTCAAGATAGCGGCG |
| A424S | A445S | AGCCATAAAUTAGGTGTCACTCAGTGTA | ATTTATGGCUCAACAAAGAAGCGACAATGTTTCC |
| G765S | G786S | ACCAGCGUCTATGGCTTCTGGAAGCA | ACGCTGGUGAGAGTGAGAAGCCTCTGAGAGT |
| E780G/R783G | E801G/R804G | AGCCGCGGAUATCTTGAGATGTTCTATGC | ATCCGCGGCUTCCACGGCGGTCAAGGTTAGACACA |
| E786G/M787G | E807G/M808G | AGGATTCUATGCTCTCAAGTACCGCAAA | AGAATCCUCCAAGATATCCGCGGCTTCCA |
| F159G/R162G | F180G/R183G | AAGGACUCAACGGACACCTTTCTGCCAAACTCT | AGTCCTUGCACACCATTTCCAATTGACTTGT |
| A166G/H170G | A187G/H191G | AACTCTUCGGAGACAAGGAGAGCTTGCAC | AAGAGTUTTCCAGAAAGGTGTCCGTTGAGTCCTTG |
| A166C/F169C | A187C/F190C | AAACTCUGCCACGACAAGGAGAGCTTG | AGAGTTUGCAAGAAAGGTGACGGTTGAGGAATTG |
| E786K/A790S/L791C | E807K/A811S/L812C | ATGTTCTAUAGCTGCAAGTACCGCAAATTGGCT | ATAGAACAUTTTAAGATAGCGGCGGCTCTCA |
| L453F/V671F/S152Y/N775K/S523P/A395M/E780G | L474F/V692F/S173Y/N796K/S544P/A416M/E801G | ACCGCCGUGGCAGCCGCCGCTATCTTGAG | ACGGCGGUCAAGTTTAGACACATGCT |
| *Ac*SuSy-L637M/T640V | *Ac*SuSy-L658M/T662V | ATAAAGUGGTTGCCGGTGAACTGTATCG | ACTTTAUCCATCAGTGCACCCACCCAGCG |

**Table S20.** List of residues of active site, UDP binding site, tetrameric interface.

| **Tetramer interface** | |
| --- | --- |
| **Residue** | **Position** |
| N | 130 |
| F | 131 |
| E | 134 |
| D | 136 |
| T | 148 |
| L | 149 |
| K | 151 |
| S | 152 |
| F | 159 |
| R | 162 |
| H | 163 |
| A | 166 |
| F | 169 |
| H | 170 |
| Q | 205 |
| D | 256 |
| E | 259 |
| A | 260 |
| D | 262 |
| E | 392 |
| H | 425 |
| E | 780 |
| R | 783 |
| Y | 789 |
| **Active site** | |
| H | 285 |
| Y | 296 |
| P | 297 |
| D | 298 |
| T | 299 |
| G | 300 |
| G | 301 |
| Q | 302 |
| V | 303 |
| V | 304 |
| Y | 305 |
| R | 380 |
| H | 436 |
| K | 442 |
| K | 583 |
| E | 673 |
| F | 675 |
| G | 676 |
| L | 677 |
| T | 678 |
| E | 681 |
| **UDP binding site** | |
| L | 294 |
| G | 295 |
| M | 576 |
| A | 577 |
| R | 578 |
| V | 607 |
| S | 645 |
| Q | 646 |
| M | 647 |
| N | 648 |
| R | 649 |
| V | 650 |
| R | 651 |
| N | 652 |

This list was based on comparison between crystal structure of *At*SuSy^1^ and model of WT along with previously reported UDP binding site elucidation^8^.

**Table S21.** List of added and excluded mutations from consensus sequence design.

| **Excluded mutations from Fireprot predicted mutations** | |
| --- | --- |
| Mutation | Comment |
| A166S | At the tetrameric interface |
| Q205W | At the tetrameric interface |
| E259Q | Disruption of salt bridge |
| Y296L | Weaker hydrophobic interactions |
| L339I | Weaker hydrophobic interactions |
| I418L | Weaker hydrophobic interactions |
| T473A | Disruption of hydrogen bonds |
| T498F | Disruption of hydrogen bond |
| R539K | Disruption of hydrogen bond |
| F544L | Weaker hydrophobic interactions |
| M624I | Weaker hydrophobic interactions |
| S645A | At the UDP binding site |
| **Rationally added mutations** | |
| M317L | Stronger hydrophobic interactions |
| A465V | Stronger hydrophobic interactions |
| L687C | Disulfide bond formation |
| S152Y | Improved interactions at tetrameric interface |
| V132E | Hydrogen bond formation |
| V132K | Salt bridge formation |
| G216R | Hydrogen bond formation |
| V374I | Stronger hydrophobic interactions |
| Q528R | Salt bridge formation |
| C563L | Stronger hydrophobic interactions |
| M624R | Hydrogen bond formation |

**Table S22.** Selected ancestral sequences and their *in silico* predicted properties.

| **SuSy variant** | **Identity (%)** | **Similarity (%)** | **Surface charge** | **RMSD (Å)** | **Soluprot score** |
| --- | --- | --- | --- | --- | --- |
| WT | 100.00 | 100.00 | -16.8 | 0 | 0.697 |
| Ancestor 180 | 95.65 | 97.52 | -16.0 | 0.12 | 0.623 |
| Ancestor 165 | 90.07 | 94.17 | -20.1 | 0.18 | 0.688 |
| Ancestor 111 | 97.52 | 98.39 | -18.1 | 0.25 | 0.691 |
| Ancestor 101 | 93.06 | 95.79 | -19.1 | 0.27 | 0.708 |
| Ancestor 290 | 62.81 | 68.60 | -4.3 | 1.33 | 0.544 |
| Ancestor 294 | 72.06 | 74.54 | 1.7 | 1.54 | 0.623 |

Ancestor 165 indicates Anc165 as described in the main manuscript.

**Table S23.** *In silico* characterization of ProteinMPNN variants.

| **SuSy variant** | **Identity (%)** | **Similarity (%)** | **Surface charge** | **RMSD (Å)** | **Soluprot score** |
| --- | --- | --- | --- | --- | --- |
| WT | 100.0 | 100.0 | -16.80 | 0 | 0.697 |
| SuSy-T1 | 78.81 | 83.78 | -36.36 | 0.306 | 0.723 |
| SuSy-T2 | 78.69 | 84.99 | -41.26 | 0.303 | 0.663 |
| SuSy-T3 | 77.12 | 84.75 | -30.23 | 0.339 | 0.642 |
| HPMN | 94.9 | 95.65 | -16.14 | 0.116 | 0.746 |

SuSy T-1, T-2, and T-3 where designed through the non-conservative strategy, while HPMN was designed through the conservative strategy.

**Table S24.** *In silico* design of oligomeric mutations.

| **Mutant** | **Predicted effect** | **Molecular contacts** | **Predicted binding affinity (kcal mol^-1^)** | **Predicted dissociation constant (M)** |
| --- | --- | --- | --- | --- |
| **Interface A-B/C-D** | | | | |
| WT |  | 84 | -9.1 | 2.00E-07 |
| F131G | Disruption | 75 | -8.2 | 1.00E-06 |
| E780G | Disruption | 79 | -8.6 | 4.70E-07 |
| R783G | Disruption | 78 | -8.1 | 1.10E-06 |
| E786G | Disruption | 86 | -8.9 | 2.80E-07 |
| M787G | Disruption | 77 | -9.1 | 2.00E-07 |
| A142D | Enhancement | 86 | -9.3 | 1.40E-07 |
| E786K | Enhancement | 91 | -9.6 | 8.70E-08 |
| A790S | Enhancement | 84 | -10.5 | 1.90E-08 |
| L791C | Enhancement | 80 | -9.1 | 2.00E-07 |
| **Interface A-C/B-D** | | | | |
| WT |  | 76 | -9.6 | 8.70E-08 |
| F159G | Disruption | 72 | -9.2 | 1.80E-07 |
| R162G | Disruption | 68 | -8.6 | 4.70E-07 |
| A166G | Disruption | 74 | -9.2 | 1.90E-07 |
| H170G | Disruption | 70 | -9 | 2.50E-07 |
| R209G | Disruption | 70 | -9.2 | 1.70E-07 |
| E259G | Disruption | 68 | -8.8 | 3.40E-07 |
| A166C | Enhancement | 78 | -9.6 | 8.60E-08 |
| F169C | Enhancement | 72 | -9.6 | 8.60E-08 |
| L255N | Enhancement | 76 | -10.1 | 4.10E-08 |
| A166C,F169C | Enhancement | 76 | -9.6 | 8.50E-08 |
| A166C,F169C,L255N | Enhancement | 76 | -10.1 | 4.00E-08 |

Predicted dissociation constant at 25 °C.

**Table S25.** Solvent tolerance of discovered and engineered SuSy variants.

| **SuSy variant** | **Solvent (% v/v)** | **5** | | **10** | | **15** | | **20** | | **25** | |
| --- | --- | --- | --- | --- | --- | --- | --- | --- | --- | --- | --- |
|  | **Activity (%)** | **RA** | **SD** | **RA** | **SD** | **RA** | **SD** | **RA** | **SD** | **RA** | **SD** |
| WT | DMSO | 70.8 | 7.2 | 64.2 | 2.9 | 29.2 | 3.8 | 20.8 | 2.9 | 9.2 | 1.4 |
|  | MeOH | 95.0 | 3.5 | 41.3 | 1.8 | 19.2 | 11.5 | 3.3 | 2.9 | 13.3 | 11.8 |
|  | ACN | 85.0 | 6.6 | 9.2 | 1.4 | 5.0 | 2.5 | 10.8 | 18.8 | ND | ND |
| *Gm*SuSy-97 | DMSO | 93.8 | 6.6 | 72.8 | 7.2 | 53.7 | 4.0 | 39.5 | 4.7 | 31.5 | 2.1 |
|  | MeOH | 66.0 | 6.1 | 29.0 | 2.2 | 22.2 | 1.5 | 11.7 | 1.3 | 11.1 | 0.7 |
|  | ACN | 46.3 | 6.4 | 22.8 | 2.6 | 19.1 | 1.7 | 13.6 | 1.4 | 19.8 | 5.8 |
| Anc165 | DMSO | 137.5 | 10.0 | 97.5 | 13.9 | 89.3 | 10.2 | 86.0 | 4.5 | 72.7 | 2.5 |
|  | MeOH | 78.5 | 9.7 | 28.1 | 3.0 | 13.2 | 1.5 | ND | ND | 1.7 | 0.0 |
|  | ACN | 58.7 | 2.2 | 13.2 | 2.9 | ND | ND | ND | ND | ND | ND |
| HPMN251 | DMSO | 91.8 | 5.9 | 81.5 | 8.4 | 87.7 | 7.9 | 61.4 | 7.7 | 48.1 | 4.4 |
|  | MeOH | 69.9 | 4.8 | 37.3 | 3.3 | 23.9 | 11.7 | 22.8 | 30.0 | 1.9 | 0.6 |
|  | ACN | 58.4 | 4.4 | 14.9 | 17.3 | 1.5 | 1.1 | 4.1 | 1.6 | ND | ND |

Dimethyl sulfoxide (DMSO), methanol (MeOH), and acetonitrile (ACN). RA denotes relative activity, defined as the percentage of remaining activity compared to the activity without solvent. SD refers to standard deviation of triplicates (*n*=3).

**Table S26.** Protonation states used during the molecular dynamics simulations.

| **WT** | | **Anc165** | |
| --- | --- | --- | --- |
| **Residue** | **Protonation state** | **Residue** | **Protonation state** |
| 4 | ASH | 298 | ASH |
| 298 | ASH | 474 | ASH |
| 474 | ASH | C,D 173 | ASH |
| 96 | GLH | 96 | GLH |
| C,D 173 | GLH | 221* | GLH |
| 221 | GLH | 620* | GLH |
| 620 | GLH | A,D 780* | GLH |
| A,D 780 | GLH | 786 | GLH |
| 10 | HIP | 10 | HID |
| 44 | HIP | 23 | HIP |
| 45 | HIE | 37 | HIP |
| 103 | HIP | 45 | HIE |
| 117 | HIP | 117 | HIE |
| 163 | HID | 163 | HID |
| 170 | HIP | 170 | HIP |
| 176 | HIP | 176 | HIP |
| 185 | HIP | 185 | HIP |
| 206 | HIE | 186 | HIP |
| 229 | HIP | 229 | HIP |
| 285 | HID | 285 | HID |
| 319 | HIP | 319 | HIP |
| 359 | HIP | 359 | HIP |
| 361 | HIE | 361 | HIE |
| 396 | HIP | 425 | HIE |
| 425 | HIE | 436 | HIE |
| 436 | HIE | 458 | HIP |
| 458 | HIP | 472 | HIP |
| 472 | HIP | 497 | HIE |
| 497 | HID | 510 | HIP |
| 510 | HIP | 545 | HIP |
| 534 | HIP | 561 | HIE |
| 545 | HIP | 694 | HIP |
| 561 | HIP | 703 | HIE |
| 694 | HIP | 709 | HIP |
| 703 | HIE | 714 | HIP |
| 709 | HIP | 735 | HIP |
| 714 | HIP |  |  |
| 735 | HIP |  |  |
| 772 | HIP |  |  |

All aspartic and glutamic acid residues are deprotonated (ASP and GLU) unless otherwise specified in the table. The protonation states of *Gm*SuSy-97 are identical to those of WT. Some protonation states are chain-dependent; therefore, the relevant chain (A, B, C, or D) is specified. For protonation states marked with an asterisk (*), the protonated oxygen alternates between OE1 and OE2.

**Table S27.** Results of full 2^5^ factorial design of MANT glycosylation.

| **Treatment** | **U/S** | **pH** | **UDP** | **Sucrose** | **Temperature** | **Conversion (%)** | **SD** |
| --- | --- | --- | --- | --- | --- | --- | --- |
| CP | 0 | 0 | 0 | 0 | 0 | 29.2 | 0.3 |
| 1 | 1 | 1 | -1 | 1 | -1 | 22.7 | 0.2 |
| 2 | 1 | 1 | 1 | 1 | 1 | 31.7 | 0.4 |
| 3 | 1 | 1 | -1 | -1 | 1 | 12.7 | 0.8 |
| 4 | -1 | 1 | -1 | 1 | -1 | 25.2 | 0.3 |
| 5 | 1 | 1 | 1 | -1 | -1 | 19.9 | 0.1 |
| 6 | 1 | -1 | 1 | -1 | -1 | 27.2 | 0.5 |
| 7 | -1 | -1 | -1 | -1 | 1 | 24.6 | 0.6 |
| 8 | 1 | -1 | 1 | -1 | 1 | 35.2 | 0.1 |
| 9 | -1 | 1 | 1 | 1 | 1 | 35.3 | 0.5 |
| 10 | -1 | 1 | 1 | -1 | -1 | 21.8 | 0.1 |
| 11 | 1 | -1 | 1 | 1 | -1 | 30.9 | 0.2 |
| 12 | -1 | 1 | -1 | -1 | -1 | 18.4 | 0.1 |
| 13 | 1 | 1 | 1 | 1 | -1 | 25.7 | 0.9 |
| 14 | 1 | -1 | -1 | 1 | -1 | 26.6 | 0.1 |
| 15 | 1 | -1 | -1 | 1 | 1 | 33.8 | 0.5 |
| 16 | 1 | -1 | 1 | 1 | 1 | 41.8 | 0.5 |
| 17 | -1 | 1 | 1 | 1 | -1 | 28.3 | 0.4 |
| 18 | -1 | -1 | -1 | 1 | -1 | 28.3 | 0.1 |
| 19 | 1 | 1 | -1 | -1 | -1 | 19.1 | 0.3 |
| 20 | -1 | -1 | 1 | -1 | 1 | 32.6 | 0.2 |
| 21 | 1 | -1 | -1 | -1 | 1 | 22.7 | 0.2 |
| 22 | -1 | -1 | -1 | -1 | -1 | 23.7 | 0.3 |
| 23 | -1 | -1 | 1 | 1 | 1 | 41.1 | 0.9 |
| 24 | 1 | 1 | -1 | 1 | 1 | 23.6 | 0.4 |
| 25 | -1 | 1 | -1 | -1 | 1 | 13.4 | 0.4 |
| 26 | -1 | 1 | 1 | -1 | 1 | 23.5 | 0.4 |
| 27 | -1 | -1 | 1 | -1 | -1 | 24.7 | 0.1 |
| 28 | 1 | -1 | -1 | -1 | -1 | 22.1 | 0.2 |
| 29 | 1 | 1 | 1 | -1 | 1 | 21.9 | 0.4 |
| 30 | -1 | -1 | 1 | 1 | -1 | 27.7 | 0.1 |
| 31 | -1 | -1 | -1 | 1 | 1 | 34.2 | 0.3 |
| 32 | -1 | 1 | -1 | 1 | 1 | 25.1 | 0.5 |

U/S stands for UGT/SuSy molar ratio (-1 = U/S:1; 1 = U/S:2; 0 = U/S: 1.5). pH (-1 = pH of 7.0; 1 = pH of 8.0; 0 = pH of 7.5). UDP (-1 = 0.125 mM; 1 = 0.5 mM; 0 = 0.3125 mM). Sucrose (-1 = 100 mM; 1 = 300 mM; 0 = 200 mM). Temperature (-1 = 35 °C; 1 = 45 °C; 0 = 40 °C). SD refers to standard deviation of triplicates (*n*=3).

**References**

1.	Zheng, Y., Anderson, S., Zhang, Y. & Garavito, R. M. The Structure of Sucrose Synthase-1 from Arabidopsis thaliana and Its Functional Implications *. *Journal of Biological Chemistry* **286**, 36108–36118 (2011).

2.	Wu, R. *et al.* The Crystal Structure of Nitrosomonas europaea Sucrose Synthase Reveals Critical Conformational Changes and Insights into Sucrose Metabolism in Prokaryotes. *J Bacteriol* **197**, 2734–2746 (2015).

3.	Gharabli, H. *et al.* Enzymatic Glycosylation of Anthranilates for Enhanced Functionality. 2025.05.29.656899 Preprint at https://doi.org/10.1101/2025.05.29.656899 (2025).

4.	Yadav, G. D. & Krishnan, M. S. An Ecofriendly Catalytic Route for the Preparation of Perfumery Grade Methyl Anthranilate from Anthranilic Acid and Methanol. *Org. Process Res. Dev.* **2**, 86–95 (1998).

5.	Wu, H. *et al.* Metabolic engineering of Escherichia coli for high-yield uridine production. *Metabolic Engineering* **49**, 248–256 (2018).

6.	李豪 & 李克进. A kind of preparation method of uridine 5’-diphosphate. (2018).

7.	Schmölzer, K., Lemmerer, M., Gutmann, A. & Nidetzky, B. Integrated process design for biocatalytic synthesis by a Leloir Glycosyltransferase: UDP‐glucose production with sucrose synthase. *Biotech & Bioengineering* **114**, 924–928 (2017).

8.	Diricks, M. *et al.* Sequence determinants of nucleotide binding in Sucrose Synthase: improving the affinity of a bacterial Sucrose Synthase for UDP by introducing plant residues. *Protein Engineering, Design and Selection* proeng **30**, 143-150 (2016).
